# Intrinsic DNA sequence determinants and tissue-specific regulation of human replication origins

**DOI:** 10.64898/2025.12.28.696672

**Authors:** Marcell Veiner, Marina Salvadores, Iván Galván-Femenía, Fran Supek

## Abstract

The accurate duplication of the genome relies on the spatiotemporal control of DNA replication initiation, yet the determinants specifying mammalian origin locations remain elusive. By developing ORIFormer, a transformer-based neural network, we decode a complex, conserved DNA sequence grammar that accurately predicts initiation sites. This approach uncovers novel determinants, which we validate by demonstrating selection and molecular effects of motif-altering genetic variants. To characterize tissue-specific usage, we developed MuSAS, a statistical genomic method leveraging widespread mutational strand asymmetries, such as from APOBEC and mismatch repair deficiency, to map initiation zones across diverse somatic tissues. Integrating these modalities reveals that while intrinsic DNA sequence features establish a high-potential landscape of constitutive origins, tissue-specific usage is governed by local chromatin accessibility acting as a permissive switch. We suggest that human replication initiation is driven by a deterministic genetic code modulated by the epigenetic landscape, providing a unified framework for understanding genome copying.

## Introduction

DNA replication is a fundamental process in all living organisms, ensuring the accurate duplication of genetic material before cell division. In eukaryotic cells, replication initiates at specific sites (replication origins) whose selection and activation is regulated to maintain genome stability and coordinate replication with other cellular processes, such as transcription and chromatin organization (reviewed in^1,2^).

The advent of high-throughput sequencing technologies has enabled genome-wide mapping of replication origins in various species, including humans^3,4^. Various methods have been developed to identify narrow-resolution initiation sites (IS), or wider-resolution initiation zones (IZ), including short nascent strand sequencing (SNS-Seq)^5,6^, initiation site sequencing (Ini-Seq)^7^, and Okazaki fragment sequencing (OK-Seq)^8^. These studies have revealed that mammalian replication origin loci are not evenly distributed across the genome but are often associated with specific genomic features, such as transcription start sites, CpG islands, strand asymmetries, and G-quadruplex structures ^4–6,9^.

Despite the progress in mapping replication origins, the molecular mechanisms underlying their selection and regulation remain poorly understood. Previous studies have identified several DNA sequence motifs enriched at origins, embodied in the origin G-rich repeated elements (OGREs)^10^ and the related G-rich hyper-motifs^11^. However, these motifs are not detectable at many bona fide origin loci, and, conversely, many loci harboring a motif do not show origin activity. Thus, the known DNA sequence determinants only partially explain the observed distribution of initiation sites. It is less well known if there exist additional, more complex DNA sequence patterns that underlie replication origin activity; the alternative explanation is that DNA sequence requirements of origins in mammals are lax and that origin regulation occurs largely in a sequence-independent manner.

Moreover, it is less clear to what extent cell types differentially use a particular subset of origin loci, and what subsets these would be for various tissues. While many origins appear to be constitutively active across different cell types, there was variation noted across individual examples of human cell lines^6,12–14^, suggesting cell type as a possible explanation. Replication origin density predicts the domain-scale DNA replication timing (RT), and the RT programs do exhibit tissue-specific usage patterns^15,16^. This suggests that additional factors, such as chromatin structure and transcriptional activity, may play a role in regulating origin activation level and/or timing, thus bearing on the domain-scale replication timing programs (reviewed in^17,18^).

To address these outstanding questions related with DNA sequence determinants of human replication origins, and their tissue-specific usage by a global, systematic analyses of genomic data, we employed two strategies. Firstly, we developed a deep learning model, ORIFormer, based on transformer neural networks, to predict replication origin loci from DNA sequence. ORIFormer accurately detects initiation sites at sub-kilobase resolution, generalizes across experimental methods and species and identifies active origin DNA sequence motifs, corresponding to certain transcription factors.

Furthermore, we investigated the variation in origin usage across human tissues by analyzing somatic mutational patterns in >1000 human cancer genomes. DNA strand biases in mutations resulting from defective DNA polymerase epsilon are able to localize origin locations in individual examples of colon cancer^19,20^. Here, we generalize this principle to various mutational signatures and tissues, using a bespoke method MuSAS, allowing us to systematically catalogue origin usage across various human somatic tissues. We reveal a multitude of tissue-specific origins, associated with domain-scale RT shifts and with local gene activation.

Overall, by integrating ORIFormer to decode the DNA sequence and MuSAS to map tissue-specific usage, we establish a hierarchical model for human origin specification. Our analyses suggest that an origin’s intrinsic DNA sequence potential (ORIFormer score) decreases as the tissue-specific usage (MuSAS signal) increases. We propose that human constitutive origins are strongly determined by their underlying DNA sequence, while the tissue-specific origins are regulated by cell type-associated epigenetic switches at a potential origin locus rather than by strong DNA sequence determinants.

## Results

### Predicting replication origin activity from DNA sequence using neural networks

To better understand the role of the DNA sequence in replication origin usage at different genomic loci, we developed a transformer-based neural network called ORIFormer, which predicts replication origin activity from sequence (Fig. 1a). As input ORIFormer takes in a DNA sequence of size 1kb, then — inspired by the neural network design of Enformer^21^ — applies a series convolutional blocks and a pooling mechanism, followed by a multi-head attention (Transformer) layer, and a series of dense layers to predict the probability of DNA replication initiation in a given 1kb window.

**Figure 1.**
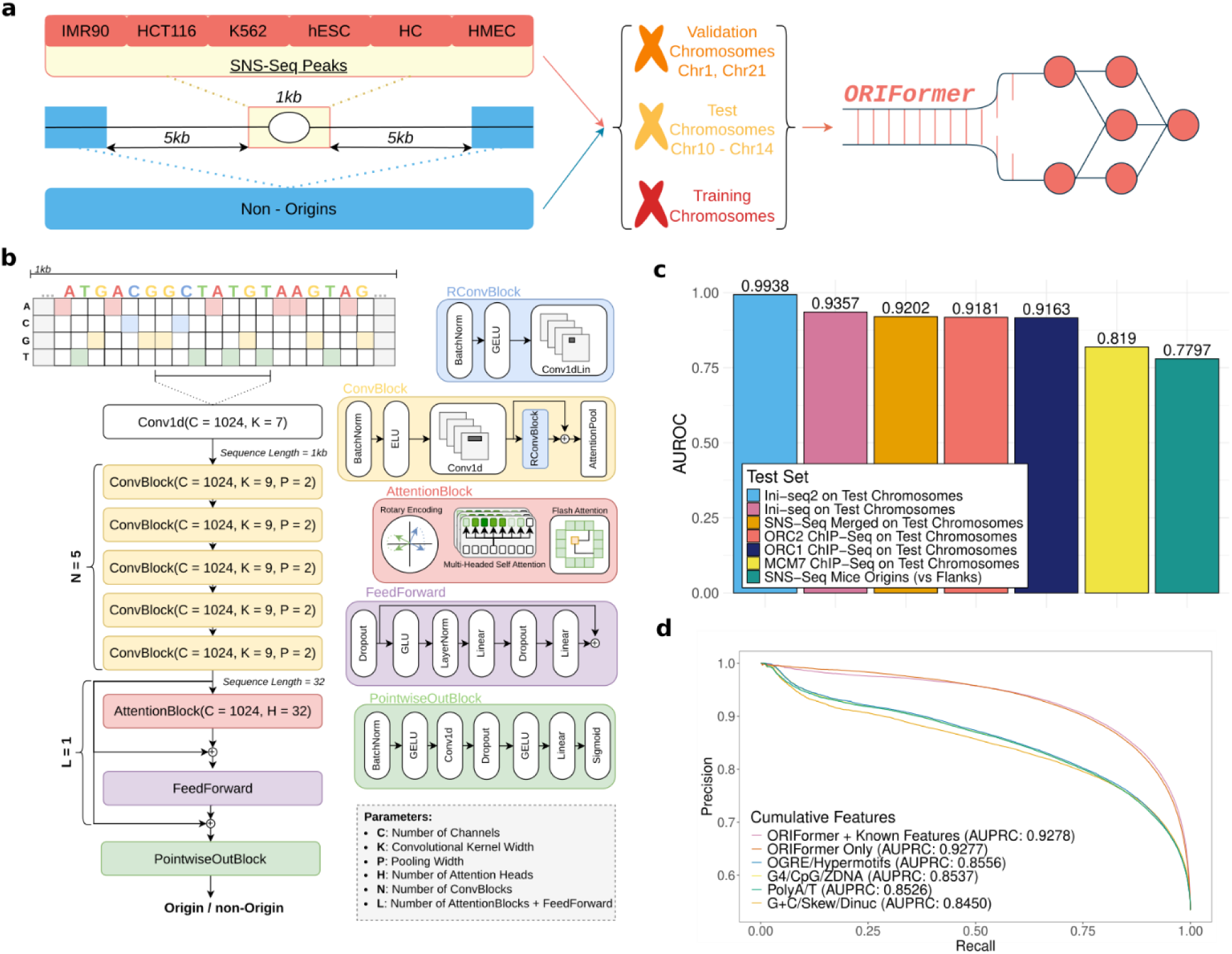
Overview of ORIFormer architecture and performance on DNA replication origin prediction. **(a)** Schematic of the dataset preparation used for training, validation, and testing of ORIFormer. **(b)** Detailed architecture of the ORIFormer model. Input nucleotide sequences of 1kb are first passed through a series of convolutional layers, which process sequence data into motifs and other higher level sequence features, which are then passed to an attention block which effectively combines these features. The output of ORIFormer is the probability of observing an active replication origin in the 1kb segment of the human genome. Key architectural parameters are listed at the bottom. **(c)** Comparison of the model’s Area Under Receiving Operator Curve (AUROC) scores across replication origins identified by different experimental assays (x-axis) on held out (test) chromosomes, and in mouse on SNS-Seq origins genomewide (green, right). **(d)** Precision-Recall curves for ORIFormer compared to other known predictive DNA sequence features. ORIFormer (dark orange line, AUPRC = 0.9277) learned all previously reported features alongside more information; there is no practical benefit (AUPRC = 0.9278) to supplementing ORIFormer with these extra, previously-known features (magenta line).

We observed that discrete DNA replication initiation sites (IS) identified by SNS-Seq^11^ and Ini-Seq2 assays^22^ can be predicted from a short sequence context (500bp on each side of the peak) at high accuracy, with AUC on held-out data >=0.92. However, broader DNA replication initiation zones (IZ) identified by OK-seq^8,23^ or RepliSeq^16^ do not have a sufficient density of DNA elements identifiable by our neural net, testing across various DNA context lengths up to 10kb (Fig. S1a, Fig. S1b). Therefore, we trained ORIFormer to predict IS, identified by 1kb windows centered on peaks in SNS-Seq, a widely applied assay to sequence RNA-primed nascent strands of DNA, with good reliability in latest iterations of the methodology^11^. To obtain a heterogeneous set of replication origins, we merged a total of 996,922 SNS-Seq peaks from 3 studies^6,11,24^ that covered a variety of cell lines, namely K562, IMR90, Hela, hESC, HMEC, CD34+ hematopoietic cells (HC), and HCT116, and multiple replicates.

To develop the ORIFormer architecture, we used Bayesian hyperparameter optimisation^25,26^, as implemented in Weights & Biases (W&B) tool; for sake of computational time, we applied this to a subset of high efficiency SNS-Seq and Ini-Seq2 origins (Fig. S2a). We identified 6 promising hyperparameter sets from W&B that corresponded to different model sizes, and in addition we manually curated 6 other hyperparameter sets. Models with these 12 neural net configurations were trained on the sequence of 1kb loci around the full set of 454,525 SNS-Seq peaks along the human genome. We note that many of these peaks were overlapping, and thus likely detected the same replication event; we leveraged this noise in the experimental measurements as a data augmentation strategy. Each of these models were then evaluated in predicting (i) SNS-Seq origins on unseen, held-out chromosomes 10-14, (ii) Ini-Seq2 origins on the same set of unseen chromosomes, and (iii) SNS-Seq identified origins in the mouse genome (Fig. S2b, Fig. S2c). The best performing neural net model (from hereon, ORIFormer) consists of ∼64 M parameters, and achieved AUC of 0.9202, 0.9938, and 0.9357 on held-out chromosomes, identifying SNS-Seq, Ini-Seq2, and Ini-Seq (from^7^) detected origins, respectively. ORIFormer also detects origin related protein binding sites from ChIP-Seq studies at good accuracy, namely 0.9163, and 0.9181 for ORC1 and ORC2, respectively, and 0.7893, 0.8619 for the two replicates for MCM7 binding sites. ORIFormer also predicted SNS-Seq identified replication origins at fair accuracy in the mouse genome (AUC=0.7797) without any further training (Fig. 1c, Fig. S3a), suggesting ORIFormer captures conserved DNA sequence determinants of replication origins across mammals.

In addition, model predictions correlated with the origin efficiency category, as defined in^11^, with mean ORIFormer prediction score of 0.932 for “Common” origins (previously defined as origins present in >=95% samples at high counts^11^), 0.879 for “Core” (for which 70-85% of replication events take place in most samples), 0.697 for “Stochastic” (hosting only 15-30% of replication activity with low activity across samples) and 0.218 mean prediction score for negative control non-origin regions (Fig. S3b). We then tested whether ORIFormer learned information not captured by previously reported DNA sequence determinants of origins. We considered G-quadruplex forming repeat sequences (G4s)^27^, G+C skew^22^, poly-A/T repeats^28^, Z-DNA motifs^22^, and a set of reported “hypermotifs” associated with SNS-seq origins^11^. Indeed, the area under the precision-recall curve (AUPRC) increased from 0.8556 to 0.9278 when adding ORIFormer predictions alongside known origin-related features (Fig. 1d), indicating that ORIFormer draws on novel sequence motifs. Conversely, adding these known DNA features alongside ORIFormer predictions did not meaningfully increase the AUPRC over the ORIFormer-only model (only by 0.0001), indicating that ORIFormer subsumed known origin-associated DNA features (Fig. 1d).

### DNA sequence determinants of human replication origins

We asked which DNA motifs were prioritized by ORIFormer. Our approach was to extract recurrent motifs, using STREME^29^, from the highest-ranking predicted set of 25,000 test sequences (i.e. from chromosomes held out during ORIFormer training). This yielded 176 motifs with nominal STREME *p* < 0.05 that are associated with high predicted origin activity (File S1). The most highly statistically supported motif M1 (Fig. 2a), GGGAGGC[CT]GAGGC[AG] (*p* < 1e-100), was found in 52% of the sequences (estimated using CentriMo^30^), and its localization exhibited a distinct peak close to centers of SNS-Seq origin loci (Fig. 2b). The second (2-GCTGGGA[CT]TACAGGCG, *p* < 1e-100, present in 49%), and third most significant motifs (3-GCACTCCAGCCTGGG[CT]GAC, *p* < 1e-100, observed in 57%) showed enrichment at distinct peaks within ∼250bs of the center (Fig. 2a, Fig. 2b). While these motifs do contain consecutive Gs / Cs, they do not fit the G-quadruplex motif definition^31^ nor a part thereof, since the G or C repetitions are interrupted or too short. The broadly mutually exclusive positioning of the top 3 motifs with respect to the SNS-seq peak location (Fig. 2b) suggest these motifs are a part of a commonly occurring broader DNA sequence arrangement favorable to replication initiation.

**Figure 2.**
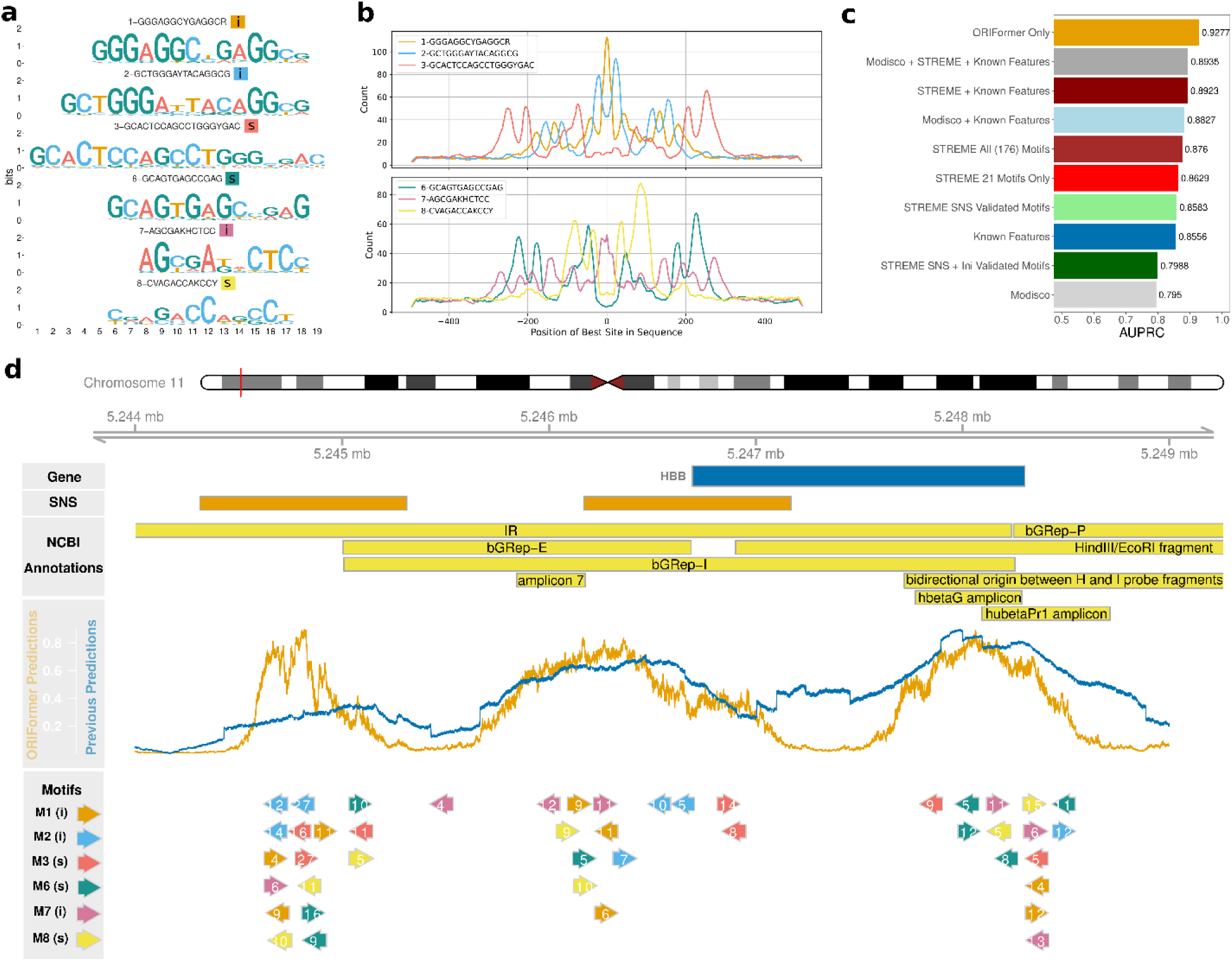
DNA motifs explain predicted replication origin activity. **(a)** Logos of the first 6 motifs (M1-M3 and M6-M8), which were discovered in a STREME enrichment analysis of ORIFormer-prioritized DNA sequences, and that additionally validate via in silico mutagenesis at experimentally measured origin loci (“s” validated via ISM in SNS-Seq origins, while “i” denotes they validated in both SNS-Seq and Ini-Seq2 origins). Colors as in panel (b). **(b)** Positional distribution of these motifs in SNS-seq peaks (0 denotes origin center). **(c)** Area Under the Precision-Recall Curve (AUPRC) achieved by different sets of motifs, compared to ORIFormer predictions (dark orange) and Known Features (blue). **(d)** Example replication origin locus near the promoter region of the hemoglobin subunit beta (HBB) gene at chr11.p15.4. Yellow bars highlight replication initiation annotations from NCBI^46^. ORIFormer predictions (orange line) are contrasted with logistic regression predictions using previously-known origin-associated features (blue line) at basepair resolution. Motif instances of the top motifs from panel (a) are shown by colored arrows, with the log odds scores and orientation of the motif matches and.

Next, we asked whether these motifs correspond to known motifs using TOMTOM^32^, querying the HOCOMOCOv11 core database for human^33^, where we found various significant matches to reported TF binding motifs (Table S1). Namely the top-ranking motif M1 GGGAGGC[CT]GAGGC[AG] had 17 matches (TOMTOM E < 2.0 and p < 0.05), of which the most significant was the Zinc Finger Protein 770 (ZNF770) motif with all 14 nucleotides overlapping (*p* = 1.23e-10, E = 4.92e-08, *q* = 9.64e-08) (File S2). We are not aware of known roles of ZNF770 in initiation of DNA replication; instead this protein has a predicted role in regulation of transcription by RNA polymerase II^34^. As reassurance, we also found the ORIFormer motif M1 to match to ZNF770 motif preferentially using a different tool (MACRO-APE^35^) in the updated HOCOMOCO v13 database; top similarity-score for ZNF770 motifs with IDs 0.P.B (similarity = 0.142) and 1.P.B (similarity = 0.128), exceeding similarity = 0.074 for the next best match (CPXCR1). Next, the ORIFormer motif M2-GCTGGGA[CT]TACAGGCG had no TOMTOM matches in the database at stringency E < 2.0. ORIFormer motif M3-GCACTCCAGCCTGGG[CT]GAC matched, at tentative significance, to binding sites of TEAD1 (E = 0.2) and NKX21 (E = 1.8); these however were not reliably replicated using the MACRO-APE search (not shown), which instead suggested ZNF560 for motif M3 (similarity = 0.182). Further, the majority of the ORIFormer motifs identified here contained significant TOMTOM matches (E < 2.0) to protein binding motifs, of which the top 3 are listed in Table S1.

These motifs were identified by analyzing highly ORIFormer-ranked replication origin DNA regions, but they could also be found in peaks from the SNS-seq assay (a random selection of 5,000 peaks analyzed; File S3). The top 3 ORIFormer motifs M1, M2 and M3 were detected in 36%, 33%, and 42% of the SNS-Seq peak sequences, respectively. 27-29% of the SNS peak sequences contained two, and 24% contained all 3 of the top 3 motifs. This result was reinforced by Spaced Motif Analysis (SpaMo)^36^, revealing that motif 2 and 3 significantly co-occur near motif 1 (E = 0.00, E = 5.02e-108) with an optimal gap size of 5 and 25 nucleotides respectively (File S4). This and above analyses suggest that the top 3 motifs we identified form parts of a local DNA sequence constellation, observable at a 100-500 nt scale, that enables or facilitates DNA replication initiation sites.

As illustrative examples, we inspected well-studied replication origin loci near the *HBB* and *PRPF4B* genes^37,38^. First, we predicted origin activity at each position in a 3.5kb and 5kb window focusing at the TSS of corresponding transcripts (respectively) using ORIFormer. We observed that ORIFormer predictions track better with known origin locations by SNS-seq, than predictions based on previously-known DNA sequence features (Fig. 2d, Fig. S9). One peak in the promoter region of the beta-globin *HBB* gene (Fig. 2d) (locus from hold-out test chromosome 11) in particular is missed when relying only on previously reported features, but ORIFormer correctly predicts origin activity potential in the region. The queried regions obtained many instances of top motifs highlighted in (Fig. 2a), co-localizing with peaks in predicted origin activity, and reported SNS-Seq peaks. The *HBB* promoter locus also contains a number of annotated initiation events^39–42^, overlapping with our predictions (Fig. 2d), and validating an ORIFormer predicted peak that was missed by SNS-Seq.

### Validation of origin-associated DNA motifs with analysis and experimental data

To further support the utility of various newly-identified DNA motifs for predicting origin loci, we used their motif counts in a DNA segment as variables in a logistic regression model for predicting SNS-Seq peaks on test chromosomes. We noted that the first 21 DNA motifs captured by ORIFormer were highly predictive of origin activity, achieving higher AUPRC (0.863 vs 0.856) than all previously reported origin-associated features (Fig. 2c), while the remaining 155 motifs explained only an additional 0.013 AUPRC points (Fig. 2c).

To support which motifs were relevant to ORIFormer predictions, we performed in silico mutagenesis (ISM) (described e.g. in^43^) at sampled motif instances from the ORIFormer top-ranked SNS-Seq and Ini-Seq2 loci, from test chromosomes held-out during ORIFormer training; top 3 instances are shown in (Fig. S4). The ISM infers the importance of every nucleotide at the sites contained within the motif, and the importances can be shown as a sequence logo (Fig. S4, Fig. S5c). In general, motif instances at example loci such as *HBB* and *PRPF4B* had clear sequence logos (Fig. 2d, Fig. S9). To assess relevance of ISM motifs to ORIFormer predictions across many loci, we compared the motif logos (pooled over 100 loci processed by ISM, with FDRs established from randomly-sampled motifs) (Fig. S5a, Fig. S5b, Fig. S5c), against the above-described logos from the STREME analysis of ORIFormer outputs (File S1). At FDR <= 10%, 64 out of the 176 original STREME motifs were validated via ISM applied to examples of actual SNS-Seq loci. As further independent support, 19 STREME motifs out of these 64 also validated via ISM of examples of Ini-Seq2 loci (Table S1, Fig. S5c). Crucially, top-ranking ORIFormer motif M1, which matched to known ZNF770 binding motif (see above), and 2nd-ranking motif M2 were validated by ISM of SNS-Seq and Ini-Seq2 peaks, while the motif M3 was ISM-validated in SNS-Seq loci. Motifs M4 and M5 did not validate in ISM, while M6 (E = 9.0e-521, observed in 55%), and M8 (E = 1.3e-399, observed in 41%) did ISM-validate in both groups and motif M7 (E = 6.9e-431, observed in 59%) only validated in SNS-Seq origins (sequences shown in Fig. 2a). Therefore, DNA motifs in addition to top-3 M1-M3 are likely to determine replication origin usage.

As a complementary approach to using STREME on the top ORIFormer predictions, we applied TF-MoDISco^44^, which extracted 14 motifs (Fig. S6), each of which could be matched (via TOMTOM) to top motifs from original STREME analysis. The 1st ORIFormer motif, corresponding to ZNF770 binding site motif, was validated herein with TF-MoDISco; interestingly, the link to ZNF770 protein binding was additionally strengthened, since the ISM reconstructed 3 additional nucleotides flanking the ORIFormer motif M1 that do match the ZNF770 site motif (Fig. S6). Overall, TF-MoDISco extracted motifs were less predictive of origins genome-wide than original STREME motifs (AUPRC 0.795 vs 0.876) (Fig. 2c), supporting the STREME-based analysis as the primary motif discovery method.

Furthermore, to mechanistically implicate proteins whose binding motifs were identified via ORIFormer analysis, we analyzed ChIP-seq tracks available in ENCODE^45^ of transcription factors (TF) that matched our ORIFormer motif list via TOMTOM or MACRO-APE. Out of the 66 combined matches, 26 had ChIP-seq experimental data available (Fig. S6), where most corresponded to proteins matching ORIFormer motifs M1 and M9. This analysis assessed local protein binding profiles at top ORIFormer predicted loci (considering only origins located on held-out test chromosomes) that also overlapped known SNS-Seq peaks. We ranked the matching TFs by two criteria: (a) enrichment of ChIP-seq signal at origin locus center vs. origin flanks (at 0.5kb distance), and (b) we also considered enrichment against control genomic loci, sampled from the same domain (within 50kb) (Fig. 3a, Fig. S8a). Considering our ORIFormer motif M1, the ChIP-seq of ZNF770 protein in particular showed strong log2 enrichment both against control regions (0.54), suggesting enrichment at origin loci, and against origin flanks (0.34), suggesting preferential binding at origin loci centers (Fig. 3a).

**Figure 3.**
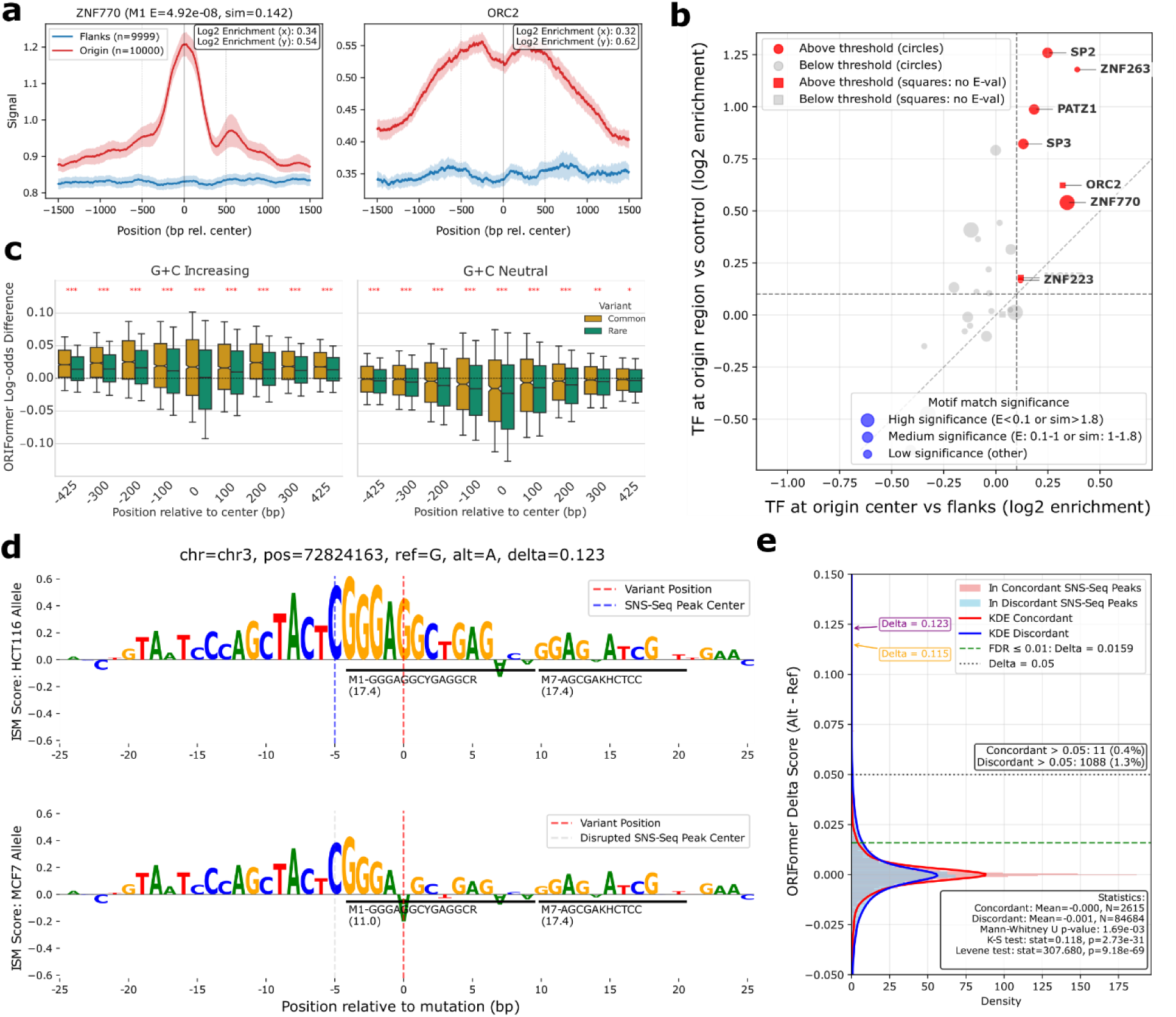
Transcription factor binding association and genetic variant effects on replication origins. **(a)** ChIP-seq signal enrichment profiles for the selected transcription factor (TF) ZNF770 binding, and ORC2 as a positive control, at replication origins (red) versus flanking regions (blue) across 3kb windows. Shaded areas show 95% confidence intervals. Enrichment scores indicate signal fold-change at origin centers relative to flanks (y-axis) and control regions (x-axis). ChIP-seq for other candidate TFs identified via ORIFormer motifs are shown in Supplementary Fig. S7 **(b)** ChIP-seq enrichments of various TFs at origin loci from test chromosomes determined by SNS-seq. Each point represents a TF with motif identified via ORIFormer analyses, and having ChIP-seq data available, with point sizes indicating closeness of match for ORIFormer motifs M1 etc. to the known TF binding motif in HOCOMOCO, based on E-values or similarity scores. Squares represent positive control factors (MCM7, ORC1, ORC2) where significance metrics for motifs do not apply since these factors were chosen by known role in replication initiation. Values along the axes denote log2 enrichment values for each TF as in the legend of panel a. Dashed lines mark 0.1 threshold for log2 enrichment **(c)** Position-dependent genetic variant effects on origin activity. Boxes show distributions of ORIFormer log-odds differences (Alt - Ref) for SNVs binned by relative position within 1kb origin regions, stratified by mutation category (G+C increasing or neutral; G+C decreasing is shown in Supplementary Fig. S11a) and variant frequency (common or rare in gnomAD population database). Notches indicate 95% C.I. Whiskers are 0.5 x IQR. **(d)** In silico mutagenesis (ISM) of an example origin-disrupting G>A variant in the MCF7 cell line genome. **(e)** Distribution of origin activity changes (by ORIFormer Delta) for variants in concordant SNS-seq replication origins in MCF7 and HCT116 cell lines (red) versus MCF7-HCT116 discordant (blue) SNS-seq origins. Density histograms compare delta scores of the genetic variants (Alt - Ref probability). Dashed lines indicate key thresholds by effect size (delta) or FDR; delta scores of individual high-origin impact variants from panels (d) and another variant in Supplementary Fig. S13 are shown as arrows. KDE: kernel density estimate.

Another strong association was observed with the binding of ZNF263 with log2 ChIP-seq enrichment 1.18 against control loci, and 0.39 enrichment against peak flanks (Fig. S8a). This TF is another, albeit less confident, match to ORIFormer motif M1 via TOMTOM (E = 1.48). Further TFs with notable binding signals are shown on (Fig. 3a, Fig. S6), where the next most confident association (by enrichment against flanks) is of motif M1 with protein SP2 (E = 0.56) (log2 enrichments 1.26 and 0.25). Overall, out of 27 tested proteins, 6 had some degree of support in this ChIP-seq analysis (having both log2 enrichments > 0.1), with ZNF770 being the only high-quality match among them (based on TOMTOM E-value) (Fig. 3b). Unsupported proteins by ChIP-seq tended to correspond to less-confident motif matches (Fig. S8a) to M1, or to other lower-ranked ORIFormer motifs. As positive controls, we included replication origin-associated proteins ORC2 and MCM7. The ChIP-seq data for these proteins had a lower resolution (Fig. S8b), consequently we compared central enrichment against origin flanks at distance 1.5kb. ORC2 exhibited notable enrichment (0.32 vs flanks, 0.62 vs control) and MCM7 more modestly so (0.12 and 0.18). Remarkably, enrichment of the newly-identified ZNF770 protein in ChIP-Seq was stronger than both positive controls; we do note a part of this difference could be due to lower resolution of the ORC2 and MCM7 data. Taken together, we provide convergent evidence to prioritize several DNA binding proteins, prominently the transcription factor ZNF770, associated with active human replication origins bearing stereotyped DNA motifs.

### Selection on replication origin motifs in the human population

As evidence of functionality of the DNA motifs determining DNA replication origin loci, we hypothesized they would be under negative selection in population variants. As a test, we compared rare (<0.1% MAF) and common variants (>1%) from GnomAD 4.0^47,48^ (Fig. S10a, Fig. S10b). To prevent confounding, we further stratified the variants into those increasing G+C content (henceforth denoted as [A|T]>[G|C]), those decreasing G+C content (denoted [G|C] > [A|T]) and preserving G+C content ([A|T] > [T|A], [G|C] > [C|G]).

To investigate whether these variants can disrupt origin activity, we selected those intersecting top predicted (inferred high-activity) origins by ORIFormer, implying they had to be enriched in the DNA motifs tested above. Next, we calculated variant effects on origin activity using ORIFormer. Variants increasing G+C content generally obtained positive ORIFormer log-odds delta scores, consistent with top-ranked origin DNA motifs being G+C rich (Fig. S10c). Rare variants of this type increased predicted origin activity significantly less (p < 0.0001) than common variants of the same types (Fig. S10c). Variants with no effect on G+C content had ORIFormer deltas closer to 0, with rare variants being modestly more disruptive than common variants (p < 0.0001). Variants that decrease G+C content all had disruptive effects on ORIFormer-predicted replication origin activity; again mirroring the trends above, the rare transversion mutations (G+C content-decreasing G > T and C > A) had significantly lower ORIFormer log-odds deltas than common mutations of the same type, while only minor differences were observed between common and rare transition mutations (Fig. S10c). Overall, within each of the 3 G+C strata of variants (G+C content increasing, preserving, or decreasing) we noted signatures of negative selection against variants that disrupt replication origin activity as assessed by ORIFormer. This analysis does not formally rule out positive selection for variants that generate origin-like DNA motifs.

We further studied short indels with length less than 100bps. Shorter deletions (<5 bp) had overall neutral ORIFormer deltas, with expectedly wider spread of effect sizes than SNVs (Fig. S10c). Larger (at least 5bp long) deletions and insertions had effects on ORIFormer predicted activity in line with the indel effect on G+C content (Fig. S10c); variant frequency distribution suggested a negative selection signal against origin disruption with insertions.

### Variant impact is concentrated at key origin-associated motifs

Furthermore, we asked if variants have different effects on ORIFormer-predicted origin activity when positioned at the origin locus center, comparing rare versus common population variants to assess selection (Fig. 3c). G+C content reducing variants were disruptive in every position, however G+C preserving and G+C increasing mutations were more deleterious towards origin locus centres than in 425 nt flanks (Fig. 3c, Fig. S11a). Furthermore, we infer that rare variants were significantly more disruptive than common variants, especially towards the origin centres (p<0.001 for all mutation types) and less so towards origin flanks (Fig. 3c), supporting negative selection assessed via ORIFormer functional impact.

This increased variant impact towards origin center is likely due to overlap with key motif instances: SNVs which overlap top 10 ORIFormer motifs had significantly higher ORIFormer deleteriousness than variants not overlapping any motifs inside origin regions (mean log-odds difference of −0.009). Effects were strongest for motifs M1 (mean of −0.065), corresponding to ZNF770, and M7 (mean of −0.078) (Fig. S11a, Fig. S11b). Rare variants were more ORIFormer-disruptive than common variants in 7 out of 10 top motifs, a signature of negative selection (Fig. S11a).

### Motif disruption with genetic variants and origin silencing in cell lines

To support that ORIFormer can identify causal effects on replication origin activity, we considered existing genetic variation in cell lines with SNS-seq data available as a “natural experiment”. Namely, we compared MCF7 versus HCT116 cell line genomes, whereby using ORIFormer, we prioritized variants that are predicted to disrupt origin activity, and asked if these indeed correspond to differential SNS-Seq signal. 84,684 variants resided in discordant replication origins (SNS peak observed in one cell line but not the other); their distributions of ORIFormer delta scores exhibited broader variability (s.d. = 0.019) than variants in concordant SNS-Seq origins (s.d = 0.010), with markedly heavier tails (Fig. 3e). Discordant SNS-seq origins showed a significantly longer tail of highly disruptive variants (Kolmogorov-Smirnov *p* = 2.7e–31; Levene’s test *p* = 9.2e–69), with 1.3% of variants predicted to be high-effect by ORIFormer (1,088/84,684), compared with only 0.4% (11/2,615) in concordant origins (Fig. 3e). These variants exhibited ORIFormer delta scores more than 0.05, corresponding to an empirical FDR threshold well below 1% (Fig. 3e).

Next, we utilized this pre-existing genetic variation to test the hypothesis that ORIFormer-identified DNA motifs have causal roles in replication origin activity. This test is based on considering whether these 1,088 variants disrupted replication origin activity in MCF7 (which exhibited no SNS-seq peaks at those loci, compared with HCT116) by interfering with the instances of top ORIFormer motifs. Indeed, variants with high ORIFormer delta (>=0.05), disrupted the binding of ORIFormer motif associated TFs significantly more than variants with ORIFormer deltas close to 0 (Fig. S12ab). Across all TFs matching ORIFormer motifs M1-M10 combined, high-effect variants by ORIFormer delta showed a mean FIMO^49^ score change of 1.23 (std=6.887, n=1,957) compared to 0.006 (std=2.815, n=16,796) for neutral variants (Mann-Whitney p<0.0001), confirming these motifs jointly encapsulate the ORIFormer model. The fraction of variants with high activity effect was 10.4% (1,957/18,753) across all MCF7-HCT116 discordant origins (Fig. S12a, Fig. S12b, Fig. S12c).

In particular, variants disrupted the binding motif of ZNF770, our top match to ORIFormer motif M1, as well as other M1-matching TF motifs (Fig. S12a, Fig. S12b). We observed n=135 (5.2%) ORIFormer high-effect variants within ZNF770 motifs at discordant SNS-seq origins, showing a mean FIMO score change Δ=2.41 (s.d.=3.046), compared to n=2,441 ORIFormer neutral variants with a mean change of Δ=-0.002 (s.d.=2.121) (Mann-Whitney U test, p<0.0001). Significant disruption patterns were also observed for other key motifs, some matching ORIFormer M1: SP2 (n=17, mean Δ=8.98), SP3 (n=17, mean Δ=8.76), and motifs PRDM6 (n=355, mean Δ=1.87), MEF2B (n=280, mean Δ=11.34) matching ORIFormer M9. For all these binding site motifs, the ORIFormer-neutral variants had mean FIMO Δ<=0.17 with all p<0.001 for significant differences for the FIMO motif match between ORIFormer high-effect and neutral variants. Considering more permissive criteria, in total 15 out of 31 analyzed binding motifs matching M1-M10 showed significant differences at p<0.05 in FIMO-disruption between high-effect and neutral variants (Fig. S12a) within the MCF7/HCT116 discordant SNS-seq peaks. We note that FIMO Δ should not be used to rank motifs; rather the significant difference in this test serves as a binary indicator that the motifs are not just correlative markers, but likely functional drivers of replication initiation in these cell lines.

The two discordant origins highlighted on (Fig. 3d, Fig. S13) showcase G+C-decreasing variants G > A, and C > T in the genome of MCF7, with highly disruptive ORIFormer delta scores, 0.123 and 0.115, respectively (FDR < 1%). Both variants disrupt the motif instance of ORIFormer M1, and thus the potential for binding of key motifs (Fig. S12a), disrupting the nearby origin activity (Fig. 3d, Fig. S13). Moreover, both instances are directly adjacent to ORIFormer motif M7 instances, highlighting one possible arrangement of the sequence constellation that defines origins. These results together suggest that ORIFormer, and the DNA motifs identified via ORIFormer analysis, can be used to infer a causal effect of genetic variation on replication origin activity, and that such variation is selected in human populations.

### Clustered ORIFormer prediction loci mark broader initiation zones

To examine how ORIFormer predictions fit into the previous observations/models of broad initiation zones (IZ) with diffuse origin activity, versus the initiation sites (IS) with focal origin activation, we considered the reconciling model where IZs are constituted of local clusters of ISs at varying activity levels. To use ORIFormer’s predictions to test this IZ/IS combined model, we used logistic regression, differentiating between IZs (of 100kbs) identified by high resolution Repli-Seq (HR Repli-Seq, from H1 embryonic stem cells^13^), and control regions of the same size and genomic distribution; the discriminatory feature is the number of initiation sites — either SNS-seq observations or ORIFormer predictions — overlapping the region. Indeed, the presence of SNS-Seq peaks were predictive of an IZ (Fig. S14), compared to randomly-placed peaks of the same number and length (0.752 vs 0.553 mean AUROC) (Fig. 4a). Remarkably, sites predicted by ORIFormer with a strong score (prediction > 0.9) were even better at classifying HR Repli-Seq IZs than experimentally measured SNS-Seq peaks, with a mean AUROC of 0.768 (Fig. 4a). Both SNS-Seq or ORIFormer peaks outperformed measured OK-Seq IZs overlapping the region (0.679 mean AUROC). IS distribution along IZs was not random, with both SNS-Seq and also the confident ORIFormer peaks clustering near the ends (+- 4kb) of OK-Seq IZs (HeLa S3 Cells), consistent with previous reports^50^ (Fig. 4c). Therefore, ORIFormer predictions support the model where the observed IZs, at least those from the high-resolution Repli-Seq assay, are local clusters of ISs with enrichment at IZ borders.

**Figure 4.**
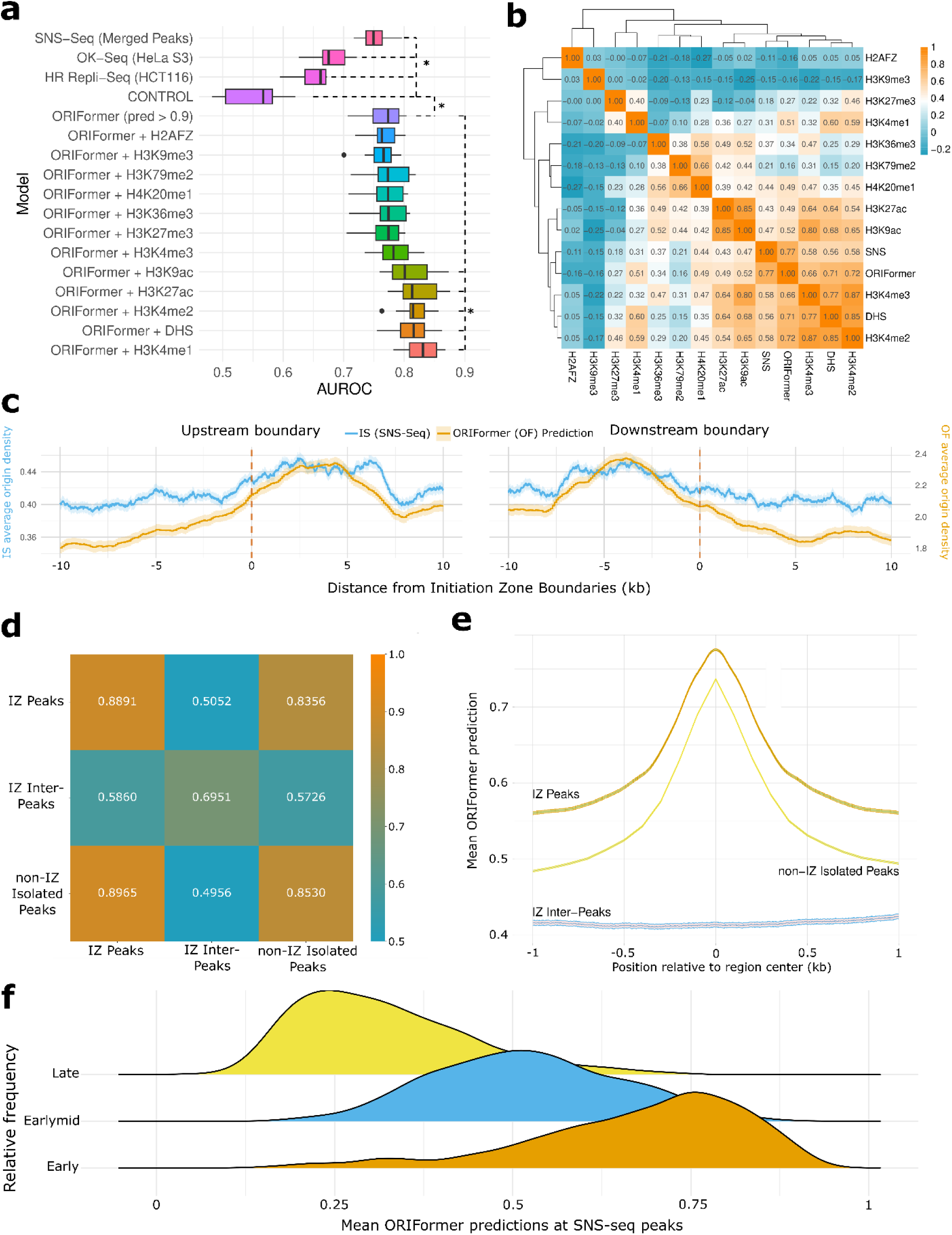
ORIFormer predictions and epigenomic tracks mark Initiation Zones (IZs), which are defined by multiple initiation sites, also determining replication timing. **(a)** Area under the ROC curve (AUROC) of logistic regression models identifying IZs experimentally determined in High Resolution Repli-Seq (HR Repli-Seq, in H1 cells). Features provided to the models are based on the local density of the features in overlapping regions, e.g. the SNS-seq model predicts IZ activity from the number of SNS-Seq peaks the IZ intersects. The CONTROL model uses the same number of randomly selected regions across the genome as the SNS-seq model. ORIFormer peaks are defined as ORIFormer score > 0.9. Epigenomic tracks use the sum of the ChIP-seq or DHS signal, instead of peak density. “+” means features were combined. “*” denotes a one-sided t-test p < 0.05. **(b)** Pearson correlation matrix of all origin and epigenomic features. **(c)** Density of SNS-Seq peaks (initiation sites, IS; blue), and confident ORIFormer predictions (> 0.9; OF, orange), +- 10kb of the upstream (left) and downstream (right) boundaries (dashed lines) of OK-Seq IZs from HeLa S3 cells. **(d)** AUROC of specialized ORIFormer models trained to detect, from DNA sequence, the SNS-Seq peaks inside IZs (top row), SNS-Seq inter-peak regions inside IZs (middle row), and SNS-Seq peaks outside of IZs (bottom row) evaluated on the test set of other models. **(e)** ORIFormer prediction around within-IZ SNS-Seq peaks (dark orange), outside of IZ SNS-Seq peaks (yellow), inter-peaks inside IZs (blue). Origin/Inter-peak regions centered at position 0 on the x-axis, line and shaded region show mean and 95% CI across loci. **(f)** Early replication timing IZs (by HR Repli-Seq) have more efficient origins, i.e. higher ORIFormer predictions averaged across all SNS-Seq peaks inside the IZ (x axis).

Although this suggests that the presence of an IZ would be, indirectly, determined by DNA sequence — which determines ISs — we note (see above, and Fig. 1) that ORIFormer does not directly predict IZ loci at high accuracy. We interpret this as that the IZ, in addition to a local clustering of ISs, may have additional characteristics that define the IZ. We asked if the IZs are defined in part by epigenetic features; indeed, including epigenomic tracks in the model further increased the mean AUC by 0.068 (Fig. 4a). Namely, DNase I Hypersensitive Sites (DHS) and active promoter and enhancer marks such as H3K27ac, H3K4me1/2/3, and K3K9ac, were the most predictive of HR Repli-Seq IZs, complementary with ORIFormer predictions. Other histone marks, most notably H2A.Z, H3K27me3, and H3K9me3, were less predictive (Fig. 4b).

Next, we generated customized ORIFormer models to test whether there is sequence specificity of ISs within IZs, or if the DNA sequence is similarly suitable for replication initiation in the whole IZ^8^. We found that SNS-Seq peaks whether inside or outside of zones can be predicted confidently by ORIFormer (0.889 and 0.853 AUC) (Fig. 4d). Evaluating the model trained on peaks inside IZ, on SNS-Seq peaks outside of IZ obtained a modestly lower performance of 0.836, suggesting that ISs inside IZs were somewhat more defined sequence-wise, but containing the same general features as the ISs outside IZs. Consistently with the above, ORIFormer predictions around detected SNS-Seq peaks were notably higher than in the regions between IS, whether in IZ or outside IZ (Fig. 4e). The DNA sequence model trained to detect inter-IS regions within IZs achieved a lower performance of 0.695. Overall, IS have strong DNA sequence determinants and IZs are defined largely via a local clustering of ISs and apart from that they have weaker DNA sequence determinants but may be defined epigenetically instead.

IZs with earlier DNA replication times (RT) tended to be denser with SNS-Seq and with ORIFormer predicted peaks (Fig. S15). Consistent with this, the mean ORIFormer prediction across an IZ varied systematically with RT profile (Fig. 4f). The RT profiles, which vary on the scale of roughly 500 kb to 1 Mb, were proposed to arise largely from different local density of origins^51^ and indeed our data is broadly consistent with that. Additionally, the mean ORIFormer prediction score for early replication origins was significantly greater than for early-mid origins (one-sided t-test, p < 2.2e-16), which in turn was greater than for late origins (p < 2.2e-16), suggesting that the origin activity level encoded in the DNA sequence, and not only their density^51^, associates with early RT of the domain. Late RT origins had weak DNA sequence determinants (ORIFormer scores in Fig. 4f), which implies they are regulated by other means.

### Detecting initiation zones de-novo via somatic mutation asymmetry

Together, our analyses indicate that many origins are specified by a conserved DNA sequence code, implying they would be largely shared across cell types. However replication programs are not uniform: RT of certain chromosomal domains (by some accounts, about half of the genome^16^) varies across tissues, implying that only a part of the origin usage is “hard-wired” in cis, whereas additional regulation layers must act in a cell type-specific manner. To dissect them, we next turned to an orthogonal readout that reports DNA replication fork direction and initiation events directly in primary human tissues: mutational strand biases in cancer genomes. As a striking individual example, tumors with impaired proofreading of the replicative DNA polymerase epsilon (POLE), characterized by hypermutation signature SBS10^52,53^, have previously been used to identify replication origins from switches in DNA strand bias polarity of mutations in 20 colon cancers^19,20^. Here, we developed a methodology that generalizes this principle to all mutational signatures and various tumor types beyond colorectal, enabling the identification of tissue-specific replication origin usage. Indeed several other signatures, such as APOBEC mutagenesis and DNA mismatch repair (MMR) deficiency, were reported to have DNA replication strand biases^53,54^, and thus represent an untapped resource for identifying replication origin loci in human somatic tissues.

We tested which mutational signatures would be useful for our purpose by systematically assessing their DNA strand asymmetry switches at known replication origin loci. As a negative control, we considered signal of transcription strand asymmetry, known to occur mechanistically independently of DNA replication and affects a distinct set of mutational processes^53^. For each mutational signature separately (and further separating the 6 mutation types within each signature), in each tissue, we computed a Mutational Strand Asymmetry Score (MuSAS, see Methods) at: (i) a consensus, high-confidence set of known origin loci supported in overlapping experimental datasets^20^, and (ii) as controls, transcription start sites (TSS) (Fig. 5a, stratified by cancer type in Fig S16a). Using randomized data, we established MuSAS thresholds for significant strand asymmetry for each combination of signature, mutation type and tissue. After having applied additional filtering (e.g. removal of artifact signatures; see Methods), we retained a subset of signatures that consistently exhibited strand asymmetry at known origins and could be used for de novo identification of DNA replication origin loci (Fig. 5a, Fig. S16). As a negative control, we checked the same signatures for transcription strand bias switches at TSS loci and did not find notable signal; as expected, we do observe TSS-colocated strand switches for the tobacco-smoking or UV-light associated signatures (Fig. 5a).

**Fig 5.**
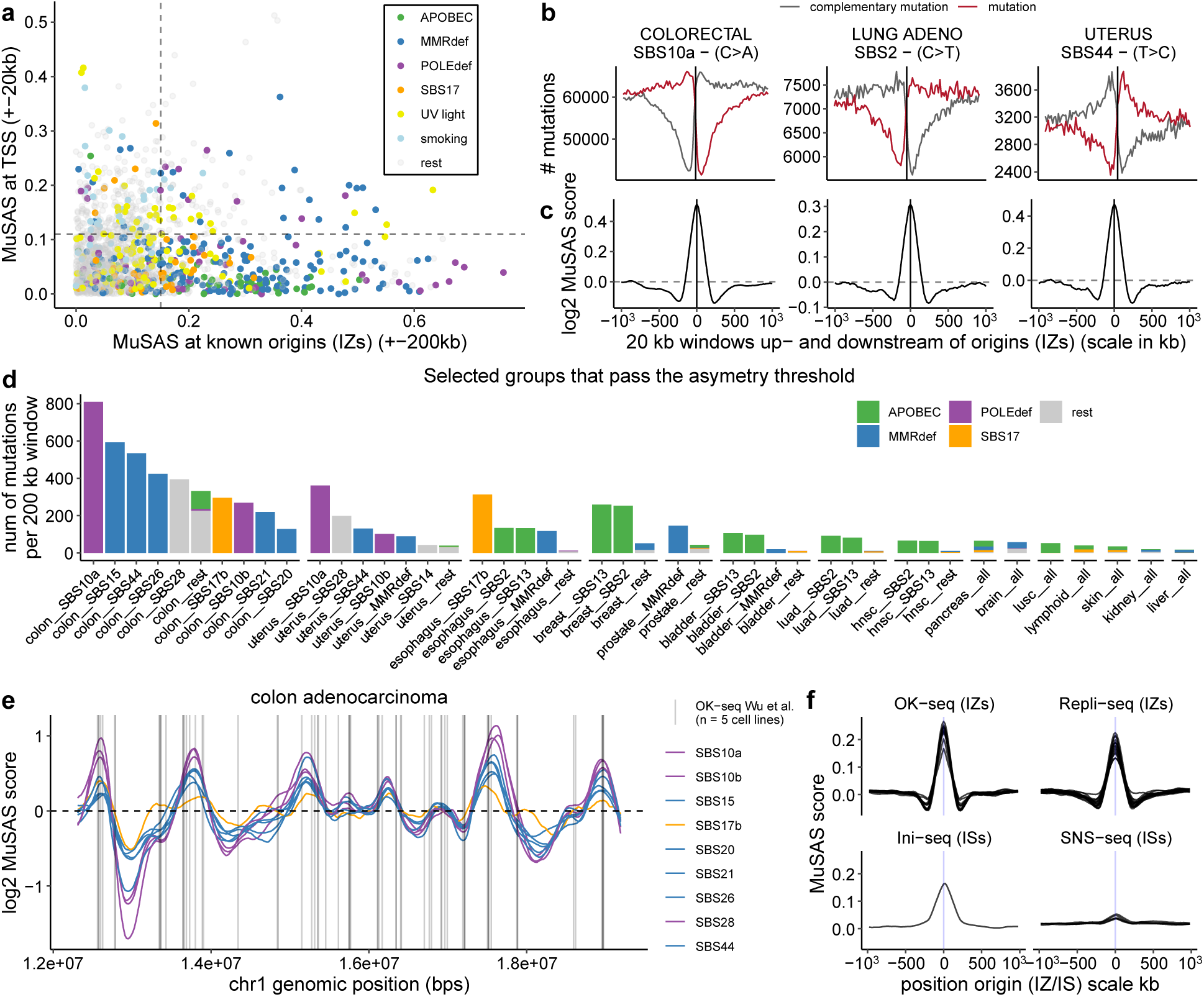
Mutational strand asymmetries identify replication initiation zones across diverse mutational signatures. **(a)** MuSAS score, measuring switches in strand asymmetry of somatic SNVs, at known origin loci (initiation zones, IZs) vs at transcription start site (TSS) loci for all mutational signatures (see color key for selected examples of SBS signatures), tissue, and mutation type (e.g. T>C, T>G, etc.) combinations. Dashed lines mark the MuSAS thresholds obtained from randomized data: (y-axis) 90% of randomized data at TSS below threshold (for removing transcriptional asymmetry); and (x-axis) 95% of randomized data at origins below threshold (for selecting asymmetry vs non-asymmetry, above and below the threshold respectively). **(b, c)** Examples of local mutation densities (local count data, b) and MuSAS scores (c) at origins for mutational signature of DNA polymerase ε (POLE) deficiency (SBS10a) in colon cancer, APOBEC mutagenesis (SBS2) in lung cancer, and MMR deficiency (SBS44) in uterus cancer. **(d)** Total number of mutations in our dataset available, per 200kb windows, in the mutational signature (SBS)-tissue combinations deemed useful for having sufficient origin-associated strand bias (panel **a**, x axis) for reliably calculating MuSAS. **(e)** MuSAS score profiles across an example chromosome 1 segment in colon adenocarcinoma for the different mutational (SBS) signatures. Known OK-seq origin loci (IZs) from 5 cell lines from ref.^23^ (except blood and brain) marked with vertical lines. **(f)** Average local MuSAS score profile, across cancer types, shown at origin positions (IS or IZ) obtained by different experimental assays.

In addition to the POLE mutational signatures (SBS10) demonstrated to localize origins^19^, mismatch repair-deficient (MMRd) signatures (e.g. SBS44, SBS26 and others) and APOBEC-related signatures (SBS13, SBS2) were effective for identifying replication origins (Fig. 5b). This is consistent with the known replication DNA strand biases of APOBEC and MMRd signatures^53,54^, where the two may in fact be mechanistically linked^55^. Another mutational signature we find broadly useful for its replication asymmetry^54^ is SBS17 (in variations SBS17a and SBS17b), a signature of uncertain aetiology. In the context of gastrointestinal cancers, it has been hypothesized to arise from oxidative damage to the free nucleotide pool (8-oxo-dGTP)^56^. These various instances of mutational signatures have considerably broader applicability to diverse tissues than the largely colon and uterus-specific SBS10^20^

To detect replication initiation loci de-novo, our methodology calculates MuSAS in 20 kb windows across the genome using WGS data and searches for the local maxima (peaks) in mutation asymmetry; as expected, we observed that MuSAS peaks at known replication origins and decreases bidirectionally (Fig. 5c). Since most tumors (75%) in our dataset have less than 5 SNV mutations/Mb, we pooled mutational data across individual tumor genomes to increase statistical power, where mutations remained stratified by cancer type. Within each cancer type, mutations from mechanistically related mutational mechanisms (e.g. SBS2 and SBS13 for APOBEC; SBS44 and SBS26 for MMRd) were grouped. This grouping is summarized in Fig. 5c, along with the final mutation counts. The statistical power of MuSAS applied to the current set of mutations allows to locate replication origin loci at 10kb-100kb scale, therefore the detected events would correspond to IZs, rather than to ISs.

We calculated joint MuSAS profiles for each mutational signature mechanistic group and identified replication IZs as local maxima for 15 tissues (example of colon in Fig. 5d). The mean number of IZs detected by tissue is 3280 with fairly small variations between tissues (Fig. S17a), with the lowest number of IZ identified in breast (n = 3112) and the highest in prostate (n = 3477). These loci of mutational bias strand switches, corresponding to IZs^20^, that were identified via MuSAS allows a comparison between somatic tissues in the replication origin usage.

We validated MuSAS by comparing it to various experimental datasets (Fig. 5f). MuSAS shows the strongest agreement with OK-seq, consistent with the resolution of the mutation-bias method, and moderate agreement with high-resolution origin loci (SNS-seq and Ini-seq). As an assessment of MuSAS accuracy, we noted that MuSAS IZs have a similar overlap with OK-seq IZs (23-32% peaks match with various OK-seq experiments in^23^), as the overlap between OK-seq and experimental IZs identified with a different assay, high-resolution Repli-seq^13^ (26-57% peaks match with OK-seq experiments peaks matching) (Fig. S17b). By this measure, the intra-method agreement between different OK-seq studies^8,23^ is broadly in the same range (30%-45% across pairs of experiments) (Fig. S17b). Thus, MuSAS asymmetry of somatic mutations can identify the IZs that OK-seq identifies, at comparable robustness to the experimental assay.

In total, we used the MuSAS method to map origins of replication across 15 tissues. For 8 out of 15 tissues there are two or more mutational signatures available (which can be considered as pseudoreplicates, useful to assess consistency of MuSAS signal in that tissue).

### MuSAS detects tissue-specific usage of DNA replication initiation zones

To assess tissue specific usage of IZs estimated via MuSAS mutation asymmetry signal, we first computed pairwise correlations between MuSAS profiles (broadly corresponding to genomic IZ likelihoods) across all mutational signature-tissue groups. Hierarchical clustering revealed that MuSAS profiles grouped primarily by tissue type, rather than by mutational signature (Fig. 6a), supporting that MuSAS captures tissue-specific patterns of replication IZ activity, while mutational signatures all reflect the same upstream signal. For downstream analyses, we retained only tissues where different mutational signatures from the same tissue cluster together by the MuSAS signal, and removed low-mutation-count, noisy profiles (e.g., lymphoid and brain), resulting in a final set of seven tissues with robust MuSAS profiles.

**Fig 6.**
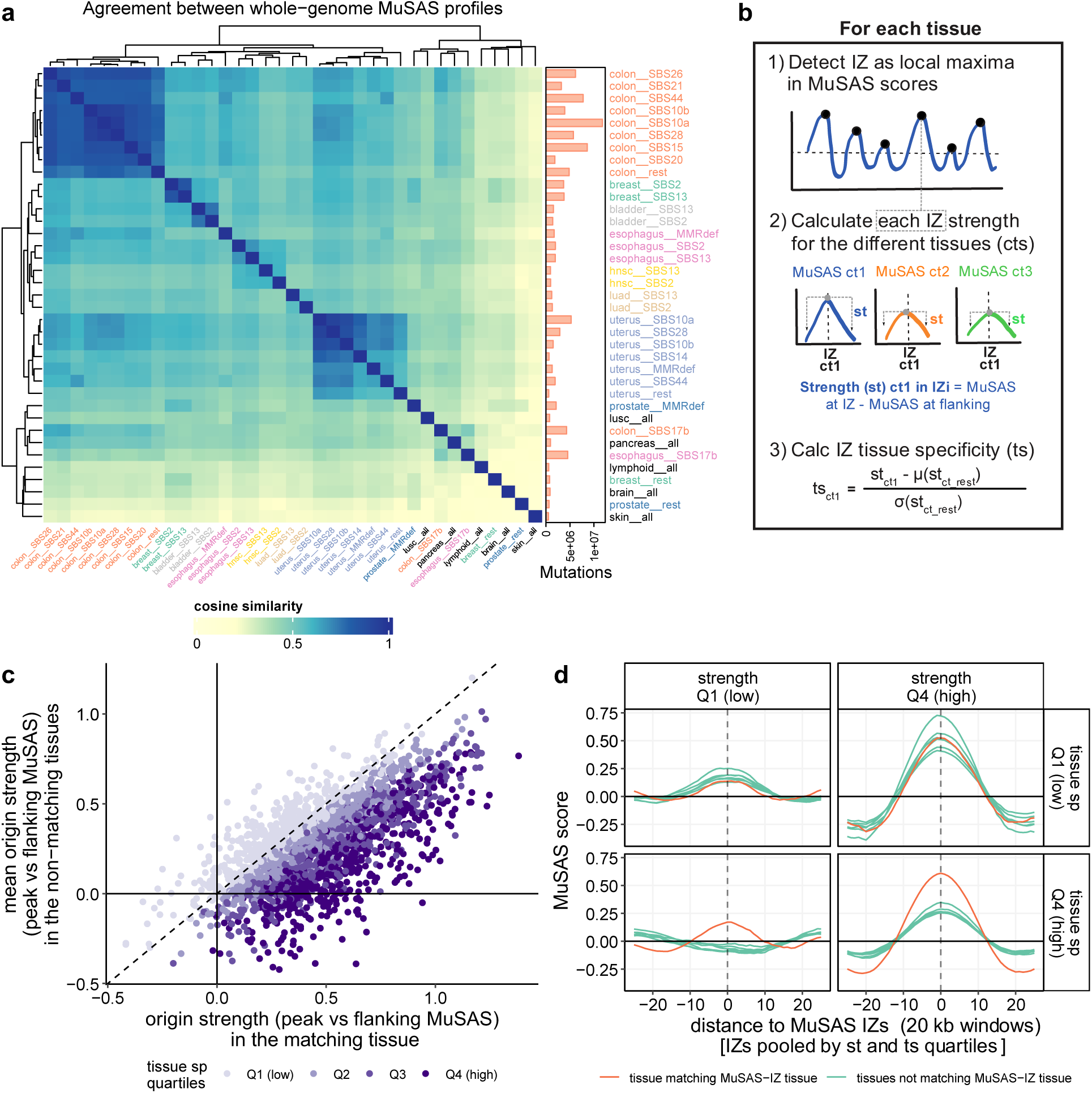
Identification and classification of tissue-specific replication origins across human cancers. **(a)** Cosine similarity between MuSAS mutation asymmetry profiles (20kb window resolution, determined across the genome) of tissue-mutational signature combinations, showing those with >500k useful mutations for MuSAS analysis (see the lateral bar chart). **(b)** Summary of the steps for method to assess tissue-specificity of initiation zones (IZ) via MuSAS. “ct”, cancer type, “st”, strength, “ts”, tissue-specificity. **(c)** Overall positive association between IZ strength in the matching tissues (here, colon shown on the x-axis, and the IZ strength in other tissues (mean across the rest of the tissues on the y-axis). Dots are IZs detected in colon, colored by tissue specificity (Z-score between IZ strength in the matched versus in the other tissues, see panel b, step 3). Q4 = most tissue specific. **(d)** Local MuSAS score profile across tissues at the identified IZs (in colon) and their flanking regions, when subdividing the IZs based on strength (columns) and tissue specificity (rows). X-axis unit is a 20kb window, thus the x axis spans −500 kb to +500 kb from the IZ.

To explore differences in origin usage across these tissues, we called discrete IZs as local MuSAS asymmetry signal peaks, independently for each tissue (Fig. 6b and Methods). For each IZ, we quantified its strength, defined as the difference between the MuSAS score magnitude at the peak (± 40 kbs) and the surrounding flanking regions (± 300 to 500 kbs), calculated separately for the tissue where it was detected and for all other tissues (Fig. 6b). This revealed that the IZ strength in a given tissue correlates with its strength in other tissues (Fig. 6c), indicating that many origins are broadly active across multiple tissues.

Additionally, we assessed tissue specificity by calculating a z-score for each IZ, comparing its MuSAS strength in the tissue of detection to the background distribution of MuSAS strengths in all other tissues (Fig. 6b). These tissue-wise z-scores identified tissue-specific IZs in principle independently of their overall strength; even origins with moderate-strength MuSAS peaks can show high tissue specificity (Fig. 6c).

By jointly considering the MuSAS strength and tissue-specific score, IZs can be classified into constitutive (active across multiple tissues, which can have higher or lower strength) and tissue-differential (showing strong bias in a specific tissue) categories. We interpret MuSAS peak strength as a proxy for origin efficiency or frequency of activation during tumor evolution, with stronger peaks reflecting more frequently used IZs.

We applied simple quartile-based thresholds, considering top-ranking 25% of origins by Z-score to exhibit a tissue-differential component, as an operational definition (tissue sp Q4 (fourth quartile), Fig. 6d). While our analysis of MuSAS profiles supports a continuous spectrum of tissue-specificity (rather than partitioning into entirely constitutive versus entirely tissue-specific subsets, Fig. 6cd), applying a quartile cutoff facilitates the study of general trends in usage of origins across the 7 human tissues with sufficient mutational signal.

These tissue-differential IZs reflect higher relative usage in a particular tissue, not exclusive activity; the peak strength in a tissue is correlated to the peak strength in the other tissues (high correlation in Fig. 6c). The tissue-differential origins with low overall strength (Quartile 1) may show a MuSAS score peak only in the matching tissue, while in stronger origins (Quartile 4), all tissues exhibit some usage (moderate effect peaks), but the MuSAS magnitude in the matching tissue remains higher (Fig. 6d).

### Tissue specific IZs are enriched in their matching tissue specific chromatin accessibility

Replication origins have been previously shown to preferentially localize to regions of open chromatin^8,57–59^, for instance near active gene promoters. Consistent with this, overall stronger IZs (MuSAS strength Q3-4) showed a clear enrichment of ATAC-seq peaks^60^ at the MuSAS peak center positions compared to flanking regions across all cancer types (Fig. S18ab). For example, this ATAC-seq signal enrichment was between 2 to 2.4-fold in colon, breast, bladder, lung and uterus, for the highest strength Q3-Q4 origins (Fig. S18ab).

We hypothesized that tissue-specific chromatin accessibility facilitates tissue-specific IZ activation. To test this, we studied if tissue-enriched origins are more active in regions with accessible chromatin in the corresponding tissue, which are less accessible in other tissues. Specifically, we assessed the overlap between MuSAS-identified origin loci and ATAC-seq peaks from TCGA tumors, stratified by cancer type^60^. This revealed that tissue-enriched origins (tissue sp Q4) showed greater enrichment of ATAC-seq peaks in the matching tissue relative to other tissues (colon shown in Fig 7ab). Specifically, for MuSAS-IZs with the highest strength (“st Q4”) and tissue specificity (“ts Q4”) we found tissue-specific chromatin accessibility enrichment of 3.4 (ATAC-seq signal fold-change in matching cancer type) vs 2.3 (fold-change in non-matching cancer type), 2.9 vs 2.0, 2.3 vs 1.7, 2.0 vs 1.8 and 3.4 vs 2.3 in colon, bladder, breast, lung adenocarcinoma, and uterus respectively (Fig. 7c-e, Fig. S18c).

**Fig 7.**
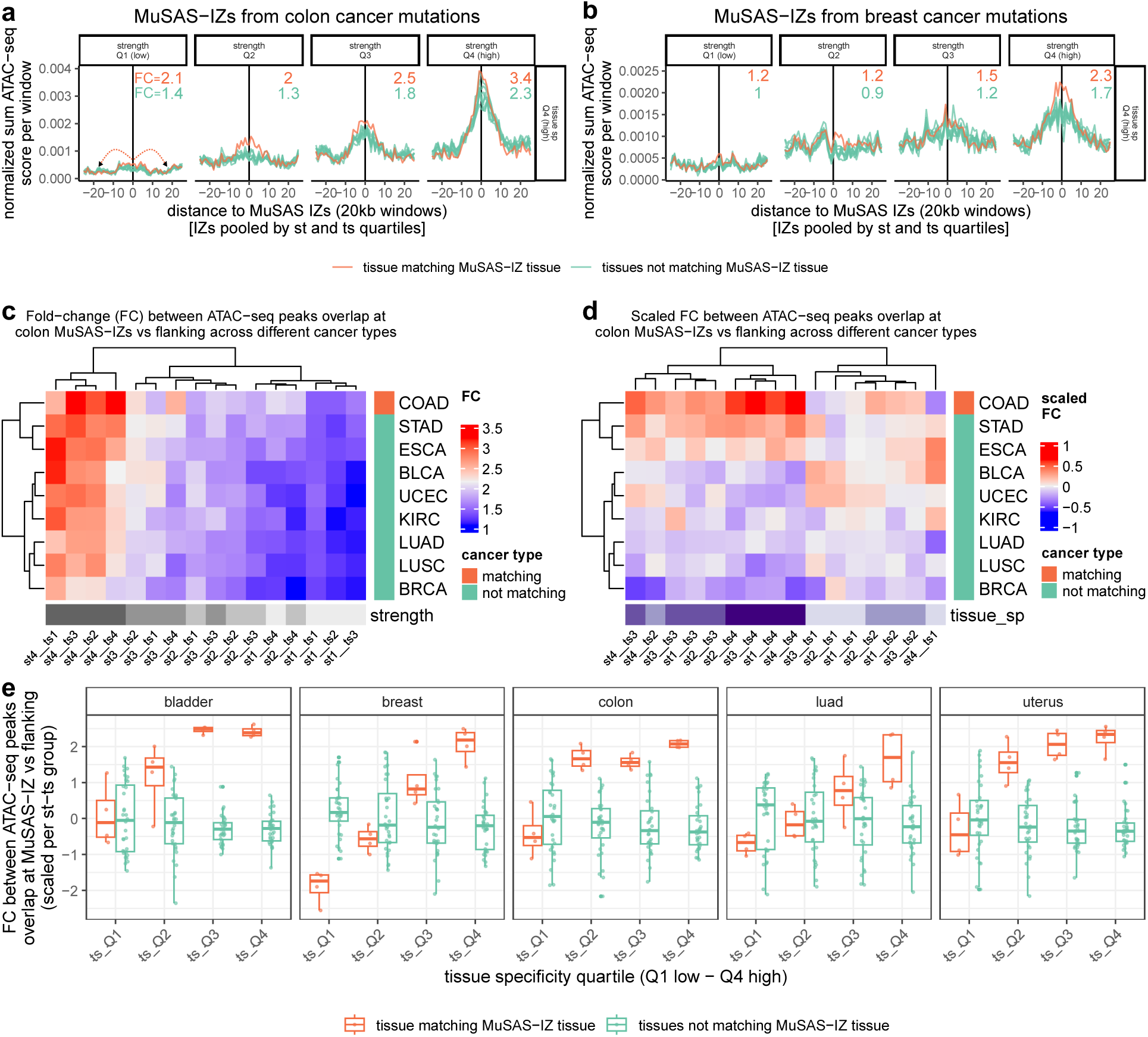
Tissue-specific replication origins localize to regions of tissue-specific chromatin accessibility. **(a)** Normalized frequency of ATAC-seq peaks (number of peaks overlapping the window divided by the number of ATAC-seq peaks in the sample) at the colon IZs (summed across IZ loci) and flanking windows across different cancer types (matching cancer type, COAD, in red). The IZs are grouped in 16 categories according to their strength (columns) and tissue specificity (the high tissue specificity, top quartile Q4 by Z-score is shown here. Numbers within each panel show the fold-change (FC) between ATAC-seq intensity at the IZ center normalized to the flanks, separately for the matching tissue, and the mean of the non-matching tissue. **(b)** Same as panel **a**, but for breast cancer IZ and breast ATAC-seq signal. **(c)** Fold-change (FC) between normalized frequency of ATAC-seq peaks at IZs (MuSAS peaks) center, versus flanking regions, for colon-IZs subdivided into 4 quartiles of strength (st1-st4) and 4 quartiles of tissue-specificity (ts1-ts4), total 16 categories. Each row represents the chromatin accessibility (ATAC-seq) of a different tissue, positively associating with origin strength). **(d)** Same as panel **c**, but FC is scaled across columns of the heatmap, adjusting for the effect of the IZ strength, emphasizing that ATAC-seq has strongest signal in matching tissue (here, colon) and related tissues (stomach, STAD, and esophagus, ESCA), in all IZ strength quartiles; other tissues shown in Supp Fig. S18. **(e)** Scaled ATAC-Seq FC values (as ones in panel **d**), shown for 5 tissues, separating by tissue-specificity quartile, and by matching vs non-matching tissue (each dot is one IZ strength category, from st1 to st4).

Thus, open chromatin may promote higher IZ activity in a tissue-specific manner, and several patterns support this mechanistic link: (i) IZs with higher MuSAS strength generally overlap more with accessible chromatin across all tissues, consistent with a relationship between origin efficiency and chromatin openness (Fig. 7c); (ii) tissue-enriched IZs show higher accessibility in their corresponding tissue, supporting the idea that tissue-specific chromatin states enable differential origin usage (Fig. 7d); and (iii) related tissues can show partial enrichment, for example, stomach (STAD) and esophagus (ESCA) share some accessibility signal at colon MuSAS origin loci, suggesting that chromatin similarity may permit similar origin activity across parts of the digestive tract (Fig. 7e).

Together, these observations are consistent with a model in which open chromatin facilitates replication origin activation, and tissue-specific differences in chromatin accessibility contribute to tissue-differential origin usage.

### Active chromatin features and DNA sequence motifs jointly influence origin usage

To explore how the DNA sequence interacts with epigenetic determinants of replication origin activity, we integrated MuSAS-identified IZs with DNA sequence-based predictions from ORIFormer. While MuSAS mutational asymmetry analysis provides IZs, at 10-100kb resolution, as detected by e.g. OK-seq or high-resolution Repli-Seq assays, ORIFormer infers ∼1kb resolution origin activity from DNA sequence (corresponding to IS as detected by e.g. SNS-seq or Ini-Seq2 assays). Comparing these complementary approaches enables us to dissect the contributions of DNA sequence and chromatin context to origin usage, further considering them in light of tissue-specificity.

We first examined whether ORIFormer predictions are enriched at MuSAS-identified IZs. Averaging ORIFormer scores (1 kb step size) across 500 kb regions centered on MuSAS peaks revealed a strong localized enrichment (Fig. 8a, Fig. S20), confirming that MuSAS-detected IZs coincide with regions of higher sequence-intrinsic IS density. Next, we stratified MuSAS IZs by tissue specificity to test whether DNA sequence contributes differentially to constitutive versus tissue-enriched IZs. ORIFormer scores averaged across each MuSAS IZ (30 kb window, with 1 kb step size) were higher for constitutive origins (low tissue-specificity, z-score Q1) compared to tissue-enriched origins (high tissue-specificity, z-score Q4) (Fig. 8b, Fig. S20). This indicates that constitutive origins are more strongly determined by DNA sequence, whereas tissue-specific origins are less so, suggesting chromatin configuration is needed for driving tissue-specific origins.

**Figure 8.**
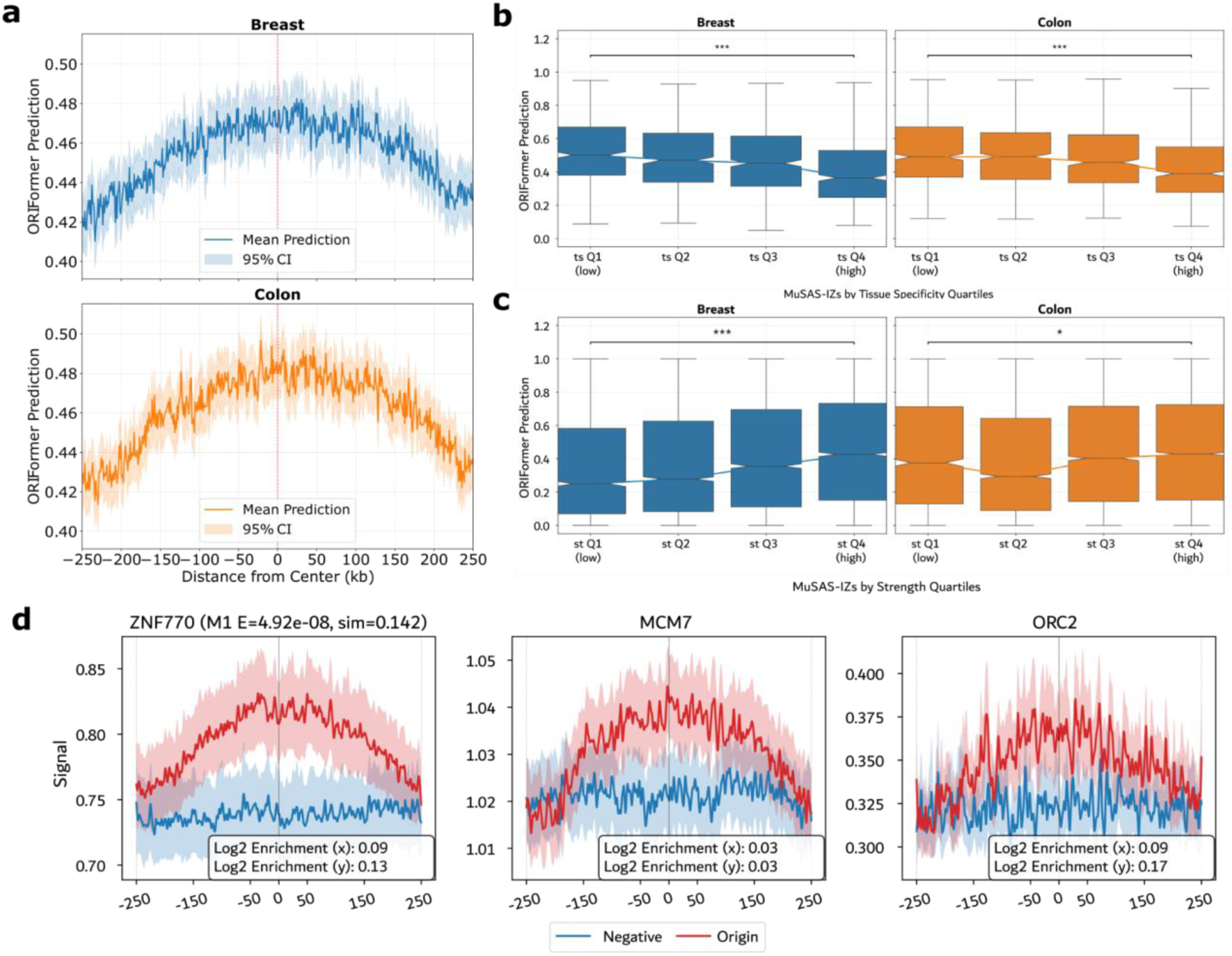
MuSAS IZs show high ORIFormer scores and enrichment in key motifs. **(a)** Rolling window analysis of ORIFormer predictions across MuSAS-detected IZs in breast (top) and colon (bottom) show elevated IS activity centered on IZs. Mean predictions ± 95% CI are shown for each 1kb window. **(b)** ORIFormer predictions stratified by MuSAS tissue-specificity (ts Q1, low, to Q4, high) at breast-detected IZs and colon-detected IZs. ORIFormer predictions (based on DNA sequence) decrease as MuSAS tissue specificity signal (based on mutation accumulation) increases (Mann-Whitney U test; *p<0.05, ***p<0.001). Lines connect group medians. Data pooled across all MuSAS IZ strength levels, the full boxplots are presented in Supplementary Fig. S20. Box plots show median (line), interquartile range (box), and whiskers extending to 1.5× IQR. **(c)** ORIFormer predictions stratified by MuSAS IZ strength levels (st Q1 to Q4), within the MuSAS high tissue-specificity IZs (ts Q4) for breast and colon cancers. ORIFormer IS predictions show modest increases with MuSAS IZ strength. **(d)** Smoothed ChIP-seq signal profiles for ORIFomer motif M1-associated factor ZNF770, as well as replication initiation associated factors MCM7, and ORC2, across MuSAS-detected IZs in colon cancer. Enrichment can be observed at MuSAS IZ centers (“Origin”) compared to non-origin, flanking genomic regions (“Negative”). Signals averaged over 500kb windows ± SEM shown. Log2 enrichment of ORIFormer signals stated in the plot indicate: (x) IZ center vs. upstream edge within IZs; (y) IZ center vs. non-origin, flanking regions center. All analyses use 30kb windows around MuSAS signature centers (panels b-c) or 500kb windows (panel d).

We then asked whether ORIFormer scores correlate with origin efficiency as determined by MuSAS strength. Stronger IZs (strength Q4) showed only a modest increase in sequence-based ORIFormer prediction compared to weaker ones (strength Q1) (Fig. 8c, Fig. S21). By contrast, chromatin accessibility showed a striking association: ATAC-seq peaks were strongly enriched in high-efficiency origins but nearly absent from low-efficiency ones (Fig. 7ab). Thus, this suggests that the DNA sequence determines a “baseline” origin potential at a locus: it is a necessary but not sufficient factor. The local chromatin configuration is the factor distinguishing highly efficient from less efficient origins; this holds true in general and in particular with respect to the tissue-specific context.

Finally, we tested the link between the candidate origin regulators prioritized via ORIFormer (Fig. 3ab) to the active IZs detected via mutational asymmetry. We analyzed ChIP-seq profiles for transcription factors corresponding to our top ORIFormer sequence motifs, centered on MuSAS IZs in colon cancer (Fig. 8d, Fig. S22). The ZNF770 (the factor matching our top-ranked motif M1) exhibited a robust, broad enrichment peaking at the center of the mutational asymmetry switch. This was mirrored by other M1-compatible factors (SP2, PATZ1, SP3) which also showed mid-point enrichment of ChIP-seq signal comparable to or higher than canonical factors MCM7 and ORC2 (Fig. S22). This provides independent validation that the DNA motifs prioritized by ORIFormer analyses indeed recruit their cognate proteins to the genomic loci where replication initiation generated mutational scars.

## Discussion

The precise nature of mammalian DNA replication origins has been a subject of long-standing debate, characterized by two apparently conflicting views: the site-specific model, supported by high-resolution initiation mapping (e.g. SNS-seq, Ini-seq), and the zonal model, supported by replication fork directionality profiling (e.g. OK-seq, Repli-seq)^3,8,11,13,22^; reviewed^61^. Our study reconciles these perspectives by a unified model where the DNA sequence establishes a high-resolution landscape of replication initiation potential, while the epigenetic context regulates the broad zonal activation of these initiation sites in a tissue-specific manner. Various previous studies implicated epigenetic contexts in controlling DNA replication^14,57,59,62^.

Contrary to the prevailing view that mammalian origins lack highly-specific sequence determinants or are governed only by lax requirements for DNA sequence, our deep learning approach reveals that human replication origins are defined by a rigorous, albeit complex, combinatorial DNA sequence code. ORIFormer successfully learned these determinants to predict initiation sites with high accuracy across species and experimental assays. Central to this sequence constellation defining origin loci is the identification of novel motifs, most notably the binding site for ZNF770 (motif M1). Beyond computational prediction, the functional importance of these motifs is evidenced by their significant enrichment in ChIP-seq signals at active initiation zones and by the signature of negative selection against motif-disrupting variants in the human population; this is consistent with previous reports of constraint at the evolutionary and population scales^38,63^. Together with our observation of variant effects on SNS-seq signal across cell line genomes, this strongly supports that the human genome contains a “hard-coded” replication program^4,11,63^ that has been evolutionarily conserved to a certain extent.

While DNA sequence defines the potential for initiation — and in our analysis the sequence code for origins was clarified — our data further supports that chromatin state acts as the switch that selects which origins fire in a given cell type^14,57^. We distinguish between two classes of origins: those with strong intrinsic sequence determinants (high ORIFormer scores) and are active across most tissues, and those which rely on permissive chromatin environments to facilitate activation and tend to be tissue-specific. This model explains why tissue-specific origins often coincide with tissue-specific open chromatin marks (ATAC-seq) and why they tend to exhibit weaker sequence motifs compared to their constitutive counterparts. The local chromatin environment might act by lowering the energetic barrier for binding of factors necessary for licensing or firing at these suboptimal sites, granting the flexibility required for developmental plasticity.

To map this tissue-specific usage, we generalized the usage of mutational strand asymmetry to localize origins, previously restricted to rare POLE-mutant tumors^19,20^, to a broader set of mutational processes. Because widespread signatures, including those resulting from APOBEC enzymes (SBS2/13) and mismatch repair deficiency (MMRd), exhibit consistent replication strand biases^53,54^, we draw on them to map out replication IZs de novo. This methodological advance allows for the “historical” mapping of replication dynamics across diverse somatic tissues using standard WGS sequencing data, reducing the need for experimental assays. We note a conceptual similarity to statistical analysis of nucleotide composition skews in the reference genome, which draws on germline mutations accumulated during evolutionary time^64,65^, however here covering a multitude of somatic tissues. While limited by mutation density to (currently) 20kb resolution, compared to the base-pair resolution of ORIFormer, MuSAS provides a robust, aggregate view of how the replication program is executed in active somatic tissues, with the caveat that cancerous cells might have altered replication programs.

In summary, our findings argue against a mostly stochastic model of mammalian DNA replication. Instead, we propose a deterministic hierarchy: a genetic foundation of precise, conserved DNA sequences (in part, including ZNF770 binding site motif) establishes the baseline competence for initiation, while an epigenetic “software” of chromatin accessibility modulates this landscape to define tissue-specific replication programs. By integrating deep learning with the mutation historical record of cancer genomes, we provide a framework to bridge the gap between the initiation site-level origin specification and the initiation zone-organization of the DNA replication program in human cells.

## Methods

### Sources of alternative human replication origin sets

In our study, we downloaded and analyzed several datasets of human replication origins. In each case we downloaded bed or equivalent files describing regions of replication events. Datasets not listed here were downloaded from original sources as described in Table number S2 of ref.^20^. Where relevant, loci in mitochondrial DNA and random contigs have been omitted. The following replication initiation datasets were analyzed:

- Ini.seq2.Guilbaud: Ini-Seq2 peaks from ref.^22^ were downloaded from Gene Expression Omnibus (GEO) with accession number GSE186675 for hg38.
- OK.Seq.HeLa.S3: OK-Seq origins for HeLa S3 cells from ref.^50^ were downloaded from GEO with accession number GSE193547 for hg19.
- HR.Repli.seq: High Resolution Repli-Seq origins for H1 and HCT116 were downloaded from ref.^13^.
- SNS.seq.Martin: SNS-Seq peaks from ref.^58^ were downloaded from DeOri database (https://tubic.org/deori/) only found K562 and MCF7 in hg19.
- SNS.seq.Picard: Data from ref.^6^ was downloaded from (http://pbil.univ-lyon1.fr/members/fpicard/oriseq/) for cell lines Hela, IMR90, K562 in hg19.
- ChIP.seq.ORC1.Dellino: ChIP-Seq Peaks for ORC1 from ref.^66^ in hg18 coordinates were downloaded in a BED format from GEO dataset GSM922790.
- ChIP.seq.ORC2.Miotto: Genomic coordinates of ChIP-Seq Peaks for ORC2 in hg19 were downloaded from the manuscript supplement Table S1 from ref.^67^.
- ChIP.seq.MCM7.Sugimoto: Genomic coordinates of ChIP-Seq Peaks for MCM7 in hg19 (exp1 and exp2) from ref.^68^ were downloaded from GEO dataset GSE107248.
- Bubble.seq.Mesner: Bubble-Seq defined origin positions from ref.^3^ were downloaded in a bedGraph format from GEO dataset GSE38809 (GM_combined_RD_bubbles file) in assembly hg18.
- SNS.seq.Akerman: Genomic coordinates of SNS-Seq peaks in hg38 were downloaded from the manuscript supplement of ref.^11^ Table number S1. Split into core (Q1 + Q2) and stochastic (Q3-Q10) oris. Common ORI locations taken from Table S1, Tab 1dCell_type_specific_and_common from region entitled Common origins (present in at least 95% of all samples at > 50 counts)
- Ini.seq.Langley: Genomic coordinates of Ini-Seq peaks from ref.^7^ were downloaded from the manuscript supplement (Table number S3) (determination by SICER at E = 10e−5) in hg19.
- SNS.seq.HCT116: SNS-Seq peaks from HCT116 in hg19 downloaded from GEO with ascension number GSE173451, from study ^24^.
- mORI.preds: Predicted coordinates (hg19) by mORI, taken from Table number S2 of ref.^20^.
- mORI.locs.50: Median-aggregated positions (hg19) of ORIs present in at least 50% of the methods as listed in Table number S3 of ref.^20^.
- mORI.locs.75: Median-aggregated positions (hg19) of ORIs present in at least 75% of the methods as listed in Table number S3 of ref.^20^.
- mORI.locs.90: Median-aggregated positions (hg19) of ORIs present in at least 90% of the methods as listed in Table number S3 of ref.^20^.
- mutORI.Bulk, Nucleotide.comp.skew.Huvet, OK.seq.Wu, OK.seq.Petryk, Repli.seq.Hansen: All locations (hg19) taken from Table number S3 of ref.^20^.

### Constructing training and test datasets for ORIFormer

We merged SNS-Seq peaks from different studies to obtain a better representation of tissues, namely a total of 996,922 SNS-Seq peaks from 3 studies^6,11,24^ with cell lines: K562, IMR90, Hela, hESC, HEMC, CD34+ Hematopoietic Cells (HC), and HCT116, and multiple replicates. Beds originally in hg38 coordinates were lifted over to hg19 using the UCSC Liftover tool (https://genome.ucsc.edu/cgi-bin/hgLiftOver).

Regions were then iteratively filtered, dropping the majority of the overlaps by selecting randomly between regions, yielding 649,531 origin regions on standard chromosomes, and thus removing about 35% of origins. To construct non-origin regions, these 649,531 segments were masked from the genome, along with any region containing unmapped nucleotides, and the remaining regions of the genome were split into 1kb regions, with a moving window of 500bps, yielding almost equal 639,692 non-origin regions. Both classes were then divided into training, validation, and test sets by chromosomes, setting aside chromosomes 10 - 14 for final testing, chromosomes 1 and 21 for validation during training, while the rest to train the models. This yielded 931,872 samples for training, 110,806 for validation, and 246,545 for testing.

In addition, 205,881 SNS-Seq peaks for mice (using assembly GRCm38), and 4,614 Ini-Seq2 peaks on test chromosomes in human were kept for final testing, for this equal number of non-origin regions from the flanks of 5 to 10kb up or downstream of each region randomly.

### ORIFormer design

ORIFormer takes as inputs DNA sequences of 1,000 nucleotides and outputs a scalar value between 0 and 1. Its design follows the design of Enformer^21^ as implemented in (https://github.com/lucidrains/enformer-pytorch) in PyTorch^69^, and comprises a series of convolutional with skip connections which enables the network to learn motifs followed by multi-headed attention with rotary encoding to learn important motif interactions.

The convolutional part of ORIFormer begins with a single 1D convolutional layer of kernel size 7, stride and dilation of 1, and padding to keep the input shape. This layer is followed by a series of 5 convolutional blocks. In our design, a convolutional block consists of a batch normalization layer followed by an ELU activation^70^, a 1D convolutional layer with stride 1, dilation 1, kernel size 9, and padding that maintains the original input shape. Output from this layer is then fed to a batch normalization layer, and GELU activation^71^, and then to a 1×1 convolutional layer, with the same stride, dilation and padding, which is then added to the output of the previous block in a residual manner. The output is sent to a pooling layer based on the attention mechanism^72^ which halves the input length. After 5 such convolutional blocks, with fixed number of channels (C = 1024), and kernel size K = 9, the output is fed to the attention block.

We use Rotary Position Embedding (RoPE)^73^ to inject positional information into the architecture, as implemented at (https://github.com/lucidrains/rotary-embedding-torch). Which together with transposed outputs from the convolutional blocks after a layer normalization step, are then fed to a single multi-head attention layer, with 32 heads, 1024 channels, and 32 key and value dimensions. Dropout with p = 0.05 is then applied to the attention scores, and 0.25 to the positional encodings. We use native flash attention as provided by PyTorch to speed up the computation. The attention mechanism is followed by a feedforward block, composed of a GLU activation function, layer normalization step, dropout with p = 0.05, and a linear layer with C = 1024 neurons.

This block is followed by a final sequential head, which consists of a batch normalization layer, GELU activation, a 1×1 convolutional block with 1024 channels, a dropout layer with p = 0.25, GELU activation, and a single neuronal layer with sigmoid activation that provides predictions.

### Model training

ORIFormer was trained on a single GPU of type NVIDIA GeForce RTX 3090, with runtime of 8d 3h. We used the AdamW optimizer^74^ with a batch size of 16, learning rate 5e-6 and weight decay rate of 9e-4, using binary cross entropy loss. The model was trained for a total of 100 epochs but reached optimum at epoch 40 which corresponded to where the loss calculated on the validation set was lowest, and model weights from that moment were reloaded as per early stopping (tolerance was set to 10 epochs).

During training, a data augmentation strategy of randomly reverse complementing half of the training samples was used, thus learning the inherent symmetry of the genome. In addition, we stochastically applied random shifting of regions by up to 100bps, as SNS-Seq peaks are not fixed. To ensure that the model is robust to class imbalance, we randomly sampled from the training set at each iteration, to match the number of samples in both classes. Model hyperparameters and metrics were logged using Weights and Biases^26^ for all developed models including ORIFormer, and can be browsed online at (https://wandb.ai/irb-gds/ORI).

### Hyperparameter search

We used Bayesian hyperparameter search as available in Weights and Biases, in what is called a sweep, with 42 model instances and the goal of minimizing binary cross entropy loss on the validation set. We defined the search space for each parameter, namely for the number of channels throughout the model, number of convolutional blocks, dilation factors, and kernel sizes, number of transformer blocks and attention heads, various dropout probabilities across the model, as well as augmentation and training parameters (for specific parameter values see the sweep on Wandb (https://wandb.ai/irb-gds/ORI). For purposes of computational efficiency, in this task, we defined a smaller training and testing set based on highly efficient origin loci from SNS-Seq and Ini-Seq2. Namely, we considered Ini-Seq2 origins, and flanking regions as non-origins as training and validation, while SNS-Seq peaks labeled as “common”, and corresponding non-origin flanking regions were held out for testing. We further tested 6 promising parameter sets from this procedure that corresponded to model sizes of about 2.8, 27, 33, 64, 85, and 100 million parameters, by training them on the full training set. This set of parameters was complemented by 6 additional manually curated (to increase model architecture diversity) parameter sets, corresponding to model sizes of 1, 11, 15, 21, 29, and 47 million parameters. The best performing parameters set across the 3 evaluation datasets (64M) was chosen as ORIFormer.

### Calculating the predictive power of known DNA features associated with replication origins

Predicted G4 and ZDNA regions in the human genome (hg19) have been downloaded from the NonB database^75^. The CpGi regions have been queried from UCSC using the ucscTableQuery function from rtracklayer with table = “cpgIslandExt”. The features related to G4/ZDNA/CpGi represent the relative overlap of the ORI region with the corresponding predicted feature.

In total the following features have been computed:

- Central.GC.content: G + C content in central 1kb
- Upstream.flank.GC.content: G + C content in [−10kb, −500bp] from peak
- Downstream.flank.GC.content: G + C content in [500bp, 10kb] from peak
- GC.skew: Compute GC Skew in 100bp sliding windows of step size 1, then divide the difference between the maximum and minimum skew, by the difference in the position of the maximum and minimum.
- AT.skew: Same for AT.
- AT.dinuc.proportion: Proportion of “AT” in the ori region
- GC.dinuc.proportion: Same for “GC“
- AC.dinuc.proportion: Same for “AC“
- TG.dinuc.proportion: Same for “TG“
- PolyA8.presence: Presence or Absence of “AAAAAAAA“
- PolyT8.presence: Same for Ts
- G4.coverage: Percentage of ori region covered by G4 predicted regions (from Non-B)
- ZDNA.coverage: Percentage of ori region covered by Z-DNA predicted regions (from Non-B)
- CpGi.coverage: Percentage of ori region covered by Z-DNA predicted regions (from UCSC)
- OGRE.motif: Presence of the Regex: [ACGT][ACGT]GG[ACGT][ACGT]GGGGGG[ACGT]GGG[ACGT]GGGGG[ACGT][ACGT]
- Hypermotifs 1-10: Presence of motifs reportred in Supplementary Figure 3b according to the following regex in order

- GG[GA][GAC]GGGG
- [GAT][GA][TG]GGG[CTA]G[GT][GA][GAC][CGT]
- CCCCTCCCCCA[CT]
- [TGA]G[CTA]G[GT]GGG[TCA]G
- [CAT]CTCCC[AC]C[AG][CG][ACT][TGCA]
- [GA][GT]GGG[TC]GTG[TG][CGT][CTG]
- [TCA][GA][GA]GG[AG][AG]G[GA][GA][CA][ACTG]
- GCCA[AC]C[GA]CCC[AC]C[TCA]
- [TAC][GAC][AG]GG[TA]CA
- [ATG]GGG[TC]G[TG]G[GT][CT]

The corresponding AUPRC values have been calculated from a fitted logistic regression by iteratively adding each (set) of features to the model. Variable importances have been obtained from the final model with all features using the caret package.

### Motif enrichment analysis

We dropped overlapping SNS-Seq peaks on test chromosomes, and ranked them by ORIFormer predictions, taking the top 25,000 sequences (of size 1kb) as our positive set. Similarly, non-origin regions were filtered and ranked, taking confidently predicted 25,000 non-origin regions as our control sequences. We used STREME^29^ to find motifs enriched in our positive set over the control sequences with minimum motif length of 7, and maximum of 30. All other parameters were the default values. STREME was run in R version 4.1.3. via the memes package version 1.11.0.

### Validating motifs via *in silico mutagenesis*

Instances of the motifs identified by STREME were found using FIMO (as implemented in tangermeme^76^ version 0.2.0, in Python 3.9.13) on the top 25,000 SNS-Seq peaks used for discovery (with a threshold of 1e-04). Hits were filtered to contain only those that appeared in the central [−100, +100] nucleotide positions around origin peaks, and which did not have mismatches in the central 5 nucleotides in the motif instance. Then for each instance of the motifs, we changed each nucleotide (e.g. A) to all other nucleotides (e.g. to C, G, or T), noting the maximum difference in predictions using the reference vs alternative nucleotides as the ISM score of that position/nucleotide. We repeated this for 100 motif instances, then averaged across them to obtain an attribution map for each motif we tested. Although these attribution maps have the same shape, and contain similar information, they are not PWMs, as the weights at a position do not sum to 1, and values can be negative.

To determine if a motif’s attribution map is similar to its original PWM, we calculated the mean Pearson correlation of the 4 nucleotide channels across all positions. We note that this method worked better than relying on the dot product. This similarity score was then compared to an empirical distribution of similarity scores comparing the attribution map to random PWMs, generated using a Dirichlet distribution, and the background nucleotide frequencies obtained from the top 25,000 SNS-Seq peaks. P-values were computed as the proportion of random similarity scores higher than the similarity score of the actual PWM. FDRs were calculated using Benjamini-Hochberg (BH).

This process was repeated with the top 10,000 Ini-Seq2 peaks (across all chromosomes) obtaining a different set of p-values. FDRs using BH were calculated by relying on the number of motifs that passed through the first FDR threshold of 10% on SNS-Seq peaks.

### Generating attribution maps and obtaining motifs via TF-MoDISco

*In silico mutagenesis* as described above were performed at each position along the top ranked 10,000 non-overlapping SNS-Seq peaks using tangermeme^76^. Motifs were obtained from ISM scores using TF-MoDISco^44^ available in modisco-lite at (https://github.com/jmschrei/tfmodisco-lite) (version 2.2.0). TF-MoDISsco was run using a maximum metacluster size of 1,000,000, flank size of 5, sliding window size of 25, trim window size of 19, and target seqlet FDR of 5%. All other parameters were at default values.

### Additional motif analysis tools

To quantify the presence of a motif in a set of sequences we relied on CentriMo^30^ as available in the meme suite webpage (https://meme-suite.org/meme/). To determine optimal spacing between a motif and a set of secondary motifs we used SpaMo^36^ available on the same website. Lastly, to search for matches in reported TF motif databases, we used TOMTOM^32^ via the command line, meme version 5.0.5, using the HOCOMOCOv11_core_HUMAN set as a database. In addition, we used the web interface for MACRO-APE^35^, with HOCOMOCO (H13CORE). We also used FIMO as implemented in tomtom-lite for motif searches^77^.

### Population genomics analysis

Population variants from GnomAD v.4.0. were downloaded from the website (https://gnomad.broadinstitute.org/news/2023-11-gnomad-v4-0/). Variants with allele frequency > 0.01 were labeled as common, while ones with < 0.001 as rare, yielding around 602M rare and 17M common variants. These were intersected with SNS-Seq replication origins to calculate the proportion of variants inside replication origins. Variants within top predicted replication origins on test chromosomes were used for variant effect prediction using ORIFormer. For indels, we only kept variants with size less than 100bps. Predictions of the replication origins with the reference and alternative alleles were computed, with the difference in logit scores as the variant effect.

### ChIP-Seq signal analysis at replication origins

To evaluate transcription factor binding enrichment at replication origins, we performed aggregate signal analysis using ChIP-seq data from 29 transcription factors. All regions were standardized to 3000 bp windows centered on replication origin coordinates. For each transcription factor, we compared ChIP-seq signal profiles between replication origin regions (labeled as 1) and matched flanking control regions (labeled as 0). Control regions were generated by randomly sampling genomic loci within ±50 kb of each origin center, maintaining a minimum distance of 1 kb from the origin and ensuring no overlap with any annotated replication origin. Signal intensities were extracted from bigWig files (fold change over control) or bedGraph files (in the case of ORC1) and averaged across all regions within each group. Mean signal profiles were computed with 95% confidence intervals using standard error of the mean, and enrichment was quantified as the log2 ratio of signal intensity at the center of origins versus flanking regions. Most datasets represent fold change over control values, providing normalized ChIP-seq enrichment signals.

GEO Accession Number by motif: GSE96190 (ATF3), GSE91415 (CUX1), GSE127656 (GATA3), GSE91861 (KLF1), GSE105407 (KLF4), GSE91633 (MAZ), GSM2863239 (MCM7), GSE127481 (MEF2B), GSE105659 (NFIC), GSE92195 (PATZ1), GSE106058 (PRDM6), GSE96235 (RXRB), GSE92014 (SP1), GSE02070 (SP2), GSE91528 (SP3), GSE96195 (TEAD1), GSE136477 (VEZF1), GSE92194 (WT1), GSE105804 (ZNF223), GSM935532 (ZNF263), GSE127634 (ZNF331), GSE127474 (ZNF449), GSE106059 (ZNF560), GSE127547 (ZNF770), GSM922790 (ORC1), GSM1717888 (ORC2).

### Genetic variation in cell line data

WGS data of cell lines MCF7 with SRA ID SRR8652105, and HCT116 with ID SRR8639145 were downloaded using SRA Toolkit (https://github.com/ncbi/sra-tools) in BAM format. Variants in both cell lines were called using Strelka2^78^ in somatic mode, using the other cell line as a normal sample. This method yielded around 1M SNVs, and 100k indels for MCF7 when compared to HCT116, and 1.5M SNVs, 500k indels in the HCT116 genome when compared to MCF7. The obtained variants were intersected with SNS-Seq centers (250bp around peak) from the respective cell lines^24,58^. Variant effect score for each variant was obtained using ORIFormer, measuring the difference in predicted probability on origin activity at the respective peak. We selected variants with large variant effects (5%) change in predicted probability, that were supported in one SNS-seq set, but not the other (meaning it is only present in one of the cell lines), and with motif instances in the vicinity of the variant. The effect of the variant was compared against all variants not-overlapping any motif instances, with statistical significance determined via one-sided t-test. The effect of the mutation is visualized by performing *in silico mutagenesis* at its neighborhood (+- 25bps on each side) and weighing each nucleotide by the respective score.

### Predicting initiation zones via IS density and epigenomic tracks

Epigenomic data was downloaded from ENCODE portal^45^ (https://www.encodeproject.org/) in narrowpeak file format using genomic coordinates hg19, for H1 embryonic stem cells. We downloaded and processed epigenomic datasets for DHS (file accession number: ENCFF001WFV), H2AFZ (ENCFF255HQX), H3K27ac (ENCFF883OGQ), H3K27me3 (ENCFF319KQT), H3K36me3 (ENCFF502LBQ), H3K4me1 (ENCFF838WSB), H3K4me2 (ENCFF105OFF), H3K4me3 (ENCFF902EBG), H3K79me2 (ENCFF270HKF), H3K9ac (ENCFF453PVM), H3K9me3 (ENCFF291YBE), and H4K20me1 (ENCFF585CSY).

We processed the High Resolution Repli-Seq data to identify Initiation Zones (IZs) in H1 embryonic stem cells^13^, standardizing their lengths to 100kb, to maintain consistency across regions, i.e. taking ±50 kb around the IZ midpoint. To generate a set of negative controls (nIZs), we randomly selected genomic regions of the same length that did not overlap with any known IZs. We focused our analysis on test chromosomes 10 to 14.

SNS-Seq peaks and high-confidence ORIFormer-predicted initiation sites (prediction >= 0.9) were integrated to enhance the identification of IZs. For each IZ and nIZ region, we extracted features by calculating the number of overlaps with SNS peaks, ORIFormer predictions, and the downloaded epigenomic datasets. Additionally, we computed the summed signal intensities (signalValue) of the epigenomic marks within these regions to capture the enrichment of specific histone modifications and accessible chromatin (as DHS).

Using these features, we trained logistic regression models to distinguish IZs from nIZs, using the above features. Model training and validation were performed using an 80/20 split for training and testing datasets, respectively, ensuring class balance between IZs and nIZs. We conducted ten replicates of this process to ensure robustness and account for variability in random sampling. Model performance was evaluated using the Area Under the Receiver Operating Characteristic Curve (AUROC), providing a quantitative measure of the models’ ability to discriminate between IZs and nIZs. Statistical significance between different models was assessed using one-sided paired t-tests, with p-values adjusted for multiple comparisons using the false discovery rate (FDR) method. This statistical analysis allowed us to determine whether the inclusion of ORIFormer predictions and specific epigenomic features significantly improved the predictive accuracy of our models.

### WGS somatic mutation collection and processing

We collected whole genome sequencing (WGS) somatic mutations from 7 different cohorts as described in^79^. The dataset included: 1950 WGS from PCAWG, 724 from TCGA, 2932 from ICGC (non-PCAWG, non-TCGA, non-TARGET), 4823 from Hartwig Medical Foundation (HMF), 1147 from CPTAC, 552 from MMRF COMMPASS, and 105 from the DECIDER project.

Sample-level metadata, including MSI status, tumor purity, ploidy, smoking history, and gender, were obtained from project data portals or supplementary publications. Cancer type labels were harmonized across datasets.

### Fitting mutational signatures to mutations

SNV calls with low mappability were filtered out according to the CRG 75 alignability track^80^ SigprofilerMatrixGenerator was used to generate the SBS-96 count matrices for all the samples (46). Then, we used SigProfilerAssignment tool^81^ to assign mutational signatures to all the mutations by using the COSMIC v3.3 mutational signatures as reference^82^. Iteratively, we classified each mutation to the most likely causal mutational signature by sampling a multinomial distribution of the vector of probabilities of each mutational signature per sample and mutation type (Decomposed_MutationType_Probabilities.txt output file from SigProfilerAssignment).

### Calculating mutational strand asymmetry score (MuSAS)

We developed a **MUtational Strand Asymmetry Score (MuSAS)** that can be calculated at any position of the genome. For each position, MuSAS is calculated by counting the number of mutations (stratified by mutation type) in both the flanking regions upstream and downstream of the position.

For origins the flanking regions considered are 200kb windows on each side, while for TSS the flanking regions are 20kb windows on each side. For each mutation type, the MuSAS score is calculated as the sum of mutations from one mutation type (for example, C>T) upstream, and its complementary (like G>A) downstream. This sum is then compared to the same alteration (C>T) downstream and its complementary (G>A) upstream. The formula is as follows:

*MuSAS* = *log*2 ((# *muts*_anchored [*upstream*] + # *muts*_*complementary* [*downstream*]) / (# *muts*_anchored [*downstream*] + # *muts*_*complementary* [*upstream*]))

A MuSAS score of 0 means that there is no asymmetry and the furthest the score diverges from 0, both positively or negatively, the more the asymmetry. The direction of the score (positive or negative) is arbitrary as it will depend on which mutation type we consider as anchored mutation and which as complementary.

To align the direction, we calculated the MuSAS score for each tissue, mutational signature and mutation types separately. We assign for each tissue, signature and mutation type which change is set as anchored mutation and which change is set as complementary, to obtain a positive MuSAS score at known origins (see below). After this alignment step, mutations from multiple signatures can be pooled while maintaining consistent directionality. This alignment only standardizes the sign and does not affect peak locations.

### Selecting the signatures with mutational strand asymmetry

We calculated the MuSAS score for every combination of tissue, mutational signature and mutation type at (i) known origins and (ii) at transcription start sites (TSSs). For the origins, we selected the origins that are represented in at least 50% of samples in the consensus from ref.^20^, removed the mutORI.Bulk and Nucleotide.comp.skew.Huvet and calculated the origin position as the median across the rest of the methods. For the origins the flanking window used is ±200kb. For the TSSs, we downloaded them from the Table browser and used flanking windows of ±20kb. For the TSS, we orient the flanking windows in relation to the gene being in the positive or negative strand.

As a control, for every tissue–signature–substitution combination separately, we shuffle the genomic coordinates across mutations in the dataset while preserving the tissue–signature–substitution information. We performed 100 permutations and for each iteration, we calculated the MuSAS score for each mutational signature, tissue and mutation type for both origins and TSSs. Based on the randomized data, we selected two thresholds: 0.15 for origins (95% of randomized data below) and 0.11 for TSSs (90% of randomized data below). We selected as useful combinations, those tissue-mutational signature-substitution combinations that had asymmetry at origins (above the randomized threshold) and did not have transcription asymmetry (below the randomized threshold).

In addition, we excluded signatures labeled as artifacts in the COSMIC database, as well as signatures that exhibit transcriptional strand asymmetry more than once (e.g., SBS7 and related UV-induced signatures), to ensure that only biologically meaningful asymmetry signals were retained.

### Determining the flanking windows size

From ref.^20^, we downloaded a comparison across origins detected with several experimental methods. We calculated the distance between origin locations across each method. The mean difference between origin positions was: 2,468,016 bps (Nucleotide.comp.skew.Huvet), 281,967 bps (OK.seq.Petryk), 314,691 bps (OK.seq.Wu), 288,764 bps (Repli.seq.Hansen) and 539,611 bps (mutORI.Bulk). The mean distance between the 5 methods is 356,258.2 bps. Based on this, we selected ±200 kb as a biologically justified and sufficiently broad origin-centered window.

### Merging mutations to increase statistical power

We computed MuSAS by merging mutations from tumors with the same tissue type and mutational signature. Because strand directionality is harmonized beforehand, aggregated mutations retain a consistent asymmetry signal.

Groups with high mutation burden were analyzed individually. Groups below a minimum threshold (50k mutations) were merged hierarchically: first by signatures of shared aetiology (e.g., APOBEC, MMR); and if still insufficient, by merging all useful signatures (asymmetric at known origins, according to our thresholds) within a tissue (naming this combined group tissue_all). The final grouping scheme is summarized in (Fig. 5c).

### Determining initiation zones (IZs) using MuSAS

We tiled the genome into 20 kb windows and computed the MuSAS score at each window. Windows with insufficient mutation counts were excluded. For each tissue, MuSAS values were log₂-transformed, sign-aligned, and analyzed separately for each chromosome arm, excluding centromeric regions.

MuSAS profiles were smoothed using a centered moving average with a window size of 20 bins (corresponding to 400 kb) and step size of one bin. Initiation zones (IZs) were defined as local maxima in the smoothed profile, requiring each peak to be the maximum within a ±10-bin neighborhood (±200 kb). Peak positions were identified independently for each tissue and represent the points where directional mutation asymmetry is most pronounced.

### Overlapping of MuSAS with experimental methods

We compiled 52 replication origin datasets obtained using diverse experimental methods (described in methods section: Sources of Alternative Human Replication Origins). For each dataset, we calculated the MuSAS score (averaged across cancer types) at the reported origin positions and their flanking regions.

For pairwise comparisons between MuSAS-derived IZs and experimental origins, we standardized genomic intervals as follows: for every experimental origin and every MuSAS-defined IZ (per tissue), we computed the midpoint of the interval and extended it symmetrically by ±25 kb to generate a uniform 50 kb window. Overlap analyses were then performed using these standardized windows to ensure comparability across datasets.

### Calculating tissue specificity in IZs

For each tissue, smoothed MuSAS profiles were generated genome-wide and replication initiation zones (IZs) were identified as local maxima in the MuSAS signal (as described above). For every IZ, we quantified: (i) IZ strength, defined as the MuSAS value at the peak relative to its flanking background, and (ii) tissue specificity, calculated as a z-score comparing the IZ strength in the tissue of detection to the distribution of strengths for the same genomic locus across all other tissues.

To categorize origins, both the IZ strength and the z-score distributions were divided into quartiles, yielding 16 IZ classes (4 strength quartiles × 4 tissue-specificity quartiles).

### Chromatin accessibility at IZs

Chromatin accessibility at IZs and flanking regions was quantified using TCGA ATAC-seq peak calls in primary human cancers^60^. We downloaded the files in All cancer type-specific peak sets [ZIP] from https://gdc.cancer.gov/about-data/publications/ATACseq-AWG

For each ATAC dataset (each dataset is a cancer type), we generated a series of windows spanning the IZ center and ±25 flanking 20 kb bins (±500 kb total). For each window, we computed the number of overlapping ATAC peaks, as well as the mean, median, and summed ATAC peak score, requiring a minimum overlap of ≥252 bp. Each window was annotated with the corresponding IZ category (from the 16-class system), allowing accessibility quantification per category and per tissue.

## Code availability

All the code related to ORIFormer, MuSAS, downstream analyses, and figure creation are available in the GitHub repository at the following link: https://github.com/Vejni/ORI

## Ackowledgements

M.V. was suppored by a fellowship from the “la Caixa” Foundation (ID 100010434, with fellowship code B006197), M.S. by an Spanish government FPU fellowship, and F.S. was funded by the ICREA Research Professor program and the University of Copenhagen. Work in the lab of F.S. is supported by an ERC StG “HYPER-INSIGHT” (757700), an ERC Consolidator Grant “STRUCTOMATIC” (101088342), Horizon2020 RIA project “DECIDER” (965193), Horizon Europe project “LUCIA” (101096473), Spanish government project “REPAIRSCAPE”, CaixaResearch project “POTENT-IMMUNO” (HR22-00402), a Novo Nordisk Fonden “Start Package” grant, the Danish Cancer Society grant “AI-DRIVERS” and a DFF Project2 (5243-00072B), the SGR funding of the Catalan government, and the Severo Ochoa Centers of Excellence award of the Spanish government to the IRB Barcelona. This publication and the underlying study have been made possible partly on the basis of the data that Hartwig Medical Foundation and the Center of Personalized Cancer Treatment (CPCT) have made available to the study. The results published here are in whole or part based on data generated by the TCGA Research Network (https://www.cancer.gov/tcga).

## Author contributions

M.V. and M.S. collected data, devised methods, implemented code, performed analyses, interpreted data and visualized results. I.G.F. prepared data and advised on analyses. F.S. devised methods, interpreted data and supervised the study. M.V., M.S. and F.S. drafted the manuscript. All authors read and approved the manuscript.

## References

1. Prioleau, M.-N. & MacAlpine, D. M. DNA replication origins—where do we begin? Genes Dev. 30, 1683–1697 (2016).

2. Fragkos, M., Ganier, O., Coulombe, P. & Méchali, M. DNA replication origin activation in space and time. Nat. Rev. Mol. Cell Biol. 16, 360–374 (2015).

3. Mesner, L. D. et al. Bubble-seq analysis of the human genome reveals distinct chromatin-mediated mechanisms for regulating early- and late-firing origins. Genome Res. 23, 1774–1788 (2013).

4. Besnard, E. et al. Unraveling cell type–specific and reprogrammable human replication origin signatures associated with G-quadruplex consensus motifs. Nat. Struct. Mol. Biol. 19, 837–844 (2012).

5. Cadoret, J.-C. et al. Genome-wide studies highlight indirect links between human replication origins and gene regulation. Proc. Natl. Acad. Sci. 105, 15837–15842 (2008).

6. Picard, F. et al. The Spatiotemporal Program of DNA Replication Is Associated with Specific Combinations of Chromatin Marks in Human Cells. PLoS Genet. 10, e1004282 (2014).

7. Langley, A. R., Gräf, S., Smith, J. C. & Krude, T. Genome-wide identification and characterisation of human DNA replication origins by initiation site sequencing (ini-seq). Nucleic Acids Res. gkw760 (2016) doi:10.1093/nar/gkw760.

8. Petryk, N. et al. Replication landscape of the human genome. Nat. Commun. 7, 10208 (2016).

9. Cayrou, C. et al. Genome-scale analysis of metazoan replication origins reveals their organization in specific but flexible sites defined by conserved features. Genome Res. 21, 1438–1449 (2011).

10. Cayrou, C. et al. New insights into replication origin characteristics in metazoans. Cell Cycle 11, 658–667 (2012).

11. Akerman, I. et al. A predictable conserved DNA base composition signature defines human core DNA replication origins. Nat. Commun. 11, 4826 (2020).

12. Massey, D. J. & Koren, A. High-throughput analysis of single human cells reveals the complex nature of DNA replication timing control. Nat. Commun. 13, 2402 (2022).

13. Zhao, P. A., Sasaki, T. & Gilbert, D. M. High-resolution Repli-Seq defines the temporal choreography of initiation, elongation and termination of replication in mammalian cells. Genome Biol. 21, 76 (2020).

14. Smith, O. K. et al. Distinct epigenetic features of differentiation-regulated replication origins. Epigenetics Chromatin 9, 18 (2016).

15. Ryba, T. et al. Evolutionarily conserved replication timing profiles predict long-range chromatin interactions and distinguish closely related cell types. Genome Res. 20, 761–770 (2010).

16. Hansen, R. S. et al. Sequencing newly replicated DNA reveals widespread plasticity in human replication timing. Proc. Natl. Acad. Sci. 107, 139–144 (2010).

17. Rhind, N. & Gilbert, D. M. DNA Replication Timing. Cold Spring Harb. Perspect. Biol. 5, a010132–a010132 (2013).

18. Marchal, C., Sima, J. & Gilbert, D. M. Control of DNA replication timing in the 3D genome. Nat. Rev. Mol. Cell Biol. 20, 721–737 (2019).

19. Shinbrot, E. et al. Exonuclease mutations in DNA polymerase epsilon reveal replication strand specific mutation patterns and human origins of replication. Genome Res. 24, 1740–1750 (2014).

20. Jaksik, R., Wheeler, D. A. & Kimmel, M. Detection and characterization of constitutive replication origins defined by DNA polymerase epsilon. BMC Biol. 21, 41 (2023).

21. Avsec, Ž., et al. Effective gene expression prediction from sequence by integrating long-range interactions. Nat. Methods 18, 1196–1203 (2021).

22. Guilbaud, G. et al. Determination of human DNA replication origin position and efficiency reveals principles of initiation zone organisation. Nucleic Acids Res. 50, 7436–7450 (2022).

23. Wu, X. et al. Developmental and cancer-associated plasticity of DNA replication preferentially targets GC-poor, lowly expressed and late-replicating regions. Nucleic Acids Res. 46, 10532–10532 (2018).

24. Mas, A. M. et al. ORC1 binds to cis-transcribed RNAs for efficient activation of replication origins. Nat. Commun. 14, 4447 (2023).

25. Shahriari, B., Swersky, K., Wang, Z., Adams, R. P. & De Freitas, N. Taking the Human Out of the Loop: A Review of Bayesian Optimization. Proc. IEEE 104, 148–175 (2016).

26. Biewald, L. Experiment Tracking with Weights and Biases. (2020).

27. Valton, A.-L. et al. G4 motifs affect origin positioning and efficiency in two vertebrate replicators. EMBO J. 33, 732–746 (2014).

28. Tubbs, A. et al. Dual Roles of Poly(dA:dT) Tracts in Replication Initiation and Fork Collapse. Cell 174, 1127–1142.e19 (2018).

29. Bailey, T. L. STREME: accurate and versatile sequence motif discovery. Bioinformatics 37, 2834–2840 (2021).

30. Bailey, T. L. & Machanick, P. Inferring direct DNA binding from ChIP-seq. Nucleic Acids Res. 40, e128–e128 (2012).

31. Maizels, N. G4 motifs in human genes. Ann. N. Y. Acad. Sci. 1267, 53–60 (2012).

32. Gupta, S., Stamatoyannopoulos, J. A., Bailey, T. L. & Noble, W. S. Quantifying similarity between motifs. Genome Biol. 8, R24 (2007).

33. Kulakovskiy, I. V. et al. HOCOMOCO: towards a complete collection of transcription factor binding models for human and mouse via large-scale ChIP-Seq analysis. Nucleic Acids Res. 46, D252–D259 (2018).

34. Camon, E. The Gene Ontology Annotation (GOA) Database: sharing knowledge in Uniprot with Gene Ontology. Nucleic Acids Res. 32, 262D –266 (2004).

35. Vorontsov, I. E., Kulakovskiy, I. V. & Makeev, V. J. Jaccard index based similarity measure to compare transcription factor binding site models. Algorithms Mol. Biol. 8, 23 (2013).

36. Whitington, T., Frith, M. C., Johnson, J. & Bailey, T. L. Inferring transcription factor complexes from ChIP-seq data. Nucleic Acids Res. 39, e98–e98 (2011).

37. Okada, N. & Shimizu, N. Dissection of the Beta-Globin Replication-Initiation Region Reveals Specific Requirements for Replicator Elements during Gene Amplification. PLoS ONE 8, e77350 (2013).

38. Murat, P. et al. DNA replication initiation shapes the mutational landscape and expression of the human genome. Sci. Adv. 8, eadd3686 (2022).

39. Aladjem, M. I. et al. Participation of the Human β-Globin Locus Control Region in Initiation of DNA Replication. Science 270, 815–819 (1995).

40. Henning, K. A. et al. Human artificial chromosomes generated by modification of a yeast artificial chromosome containing both human alpha satellite and single-copy DNA sequences. Proc. Natl. Acad. Sci. 96, 592–597 (1999).

41. Buzina, A. Initiation of DNA replication at the human -globin 3’ enhancer. Nucleic Acids Res. 33, 4412–4424 (2005).

42. Wang, L. et al. The Human β-Globin Replication Initiation Region Consists of Two Modular Independent Replicators. Mol. Cell. Biol. 24, 3373–3386 (2004).

43. Novakovsky, G., Dexter, N., Libbrecht, M. W., Wasserman, W. W. & Mostafavi, S. Obtaining genetics insights from deep learning via explainable artificial intelligence. Nat. Rev. Genet. 24, 125–137 (2023).

44. Shrikumar, A., et al. Technical Note on Transcription Factor Motif Discovery from Importance Scores (TF-MoDISco) version 0.5.6.5. Preprint at 10.48550/ARXIV.1811.00416 (2018).

45. Sloan, C. A. et al. ENCODE data at the ENCODE portal. Nucleic Acids Res. 44, D726–D732 (2016).

46. NCBI. LOC107133510 origin of replication at HBB.

47. Karczewski, K. J. et al. The mutational constraint spectrum quantified from variation in 141,456 humans. Nature 581, 434–443 (2020).

48. Chao, K. & gnomAD Production Team. gnomAD v4.0. (2023).

49. Grant, C. E., Bailey, T. L. & Noble, W. S. FIMO: scanning for occurrences of a given motif. Bioinformatics 27, 1017–1018 (2011).

50. Liu, Y., Wu, X., d’Aubenton-Carafa, Y., Thermes, C. & Chen, C.-L. OKseqHMM: a genome-wide replication fork directionality analysis toolkit. Nucleic Acids Res. 51, e22–e22 (2023).

51. Gindin, Y., Valenzuela, M. S., Aladjem, M. I., Meltzer, P. S. & Bilke, S. A chromatin structure-based model accurately predicts DNA replication timing in human cells. Mol. Syst. Biol. 10, 722 (2014).

52. Australian Pancreatic Cancer Genome Initiative et al. Signatures of mutational processes in human cancer. Nature 500, 415–421 (2013).

53. Haradhvala, N. J. et al. Mutational Strand Asymmetries in Cancer Genomes Reveal Mechanisms of DNA Damage and Repair. Cell 164, 538–549 (2016).

54. Tomkova, M., Tomek, J., Kriaucionis, S. & Schuster-Böckler, B. Mutational signature distribution varies with DNA replication timing and strand asymmetry. Genome Biol. 19, 129 (2018).

55. Mas-Ponte, D. & Supek, F. DNA mismatch repair promotes APOBEC3-mediated diffuse hypermutation in human cancers. Nat. Genet. 52, 958–968 (2020).

56. the Oesophageal Cancer Clinical and Molecular Stratification (OCCAMS) Consortium et al. Mutational signatures in esophageal adenocarcinoma define etiologically distinct subgroups with therapeutic relevance. Nat. Genet. 48, 1131–1141 (2016).

57. Cayrou, C. et al. The chromatin environment shapes DNA replication origin organization and defines origin classes. Genome Res. 25, 1873–1885 (2015).

58. Martin, M. M. et al. Genome-wide depletion of replication initiation events in highly transcribed regions. Genome Res. 21, 1822–1832 (2011).

59. Dao, F.-Y., Lv, H., Fullwood, M. J. & Lin, H. Accurate Identification of DNA Replication Origin by Fusing Epigenomics and Chromatin Interaction Information. Research 2022, 2022/9780293 (2022).

60. Corces, M. R. et al. The chromatin accessibility landscape of primary human cancers. Science 362, eaav1898 (2018).

61. Hyrien, O., Guilbaud, G. & Krude, T. The double life of mammalian DNA replication origins. Genes Dev. 39, 304–324 (2025).

62. Turner, J. L. et al. Master transcription-factor binding sites constitute the core of early replication control elements. EMBO J. 44, 4499–4524 (2025).

63. Massip, F. et al. Evolution of replication origins in vertebrate genomes: rapid turnover despite selective constraints. Nucleic Acids Res. 47, 5114–5125 (2019).

64. Chen, C.-L. et al. Replication-Associated Mutational Asymmetry in the Human Genome. Mol. Biol. Evol. 28, 2327–2337 (2011).

65. Touchon, M. et al. Replication-associated strand asymmetries in mammalian genomes: Toward detection of replication origins. Proc. Natl. Acad. Sci. 102, 9836–9841 (2005).

66. Dellino, G. I. et al. Genome-wide mapping of human DNA-replication origins: Levels of transcription at ORC1 sites regulate origin selection and replication timing. Genome Res. 23, 1–11 (2013).

67. Miotto, B., Ji, Z. & Struhl, K. Selectivity of ORC binding sites and the relation to replication timing, fragile sites, and deletions in cancers. Proc. Natl. Acad. Sci. 113, (2016).

68. Sugimoto, N., Maehara, K., Yoshida, K., Ohkawa, Y. & Fujita, M. Genome-wide analysis of the spatiotemporal regulation of firing and dormant replication origins in human cells. Nucleic Acids Res. 46, 6683–6696 (2018).

69. Paszke, A., et al. PyTorch: An Imperative Style, High-Performance Deep Learning Library. Preprint at 10.48550/ARXIV.1912.01703 (2019).

70. Clevert, D.-A., Unterthiner, T. & Hochreiter, S. Fast and Accurate Deep Network Learning by Exponential Linear Units (ELUs). Preprint at 10.48550/ARXIV.1511.07289 (2015).

71. Hendrycks, D. & Gimpel, K. Gaussian Error Linear Units (GELUs). Preprint at 10.48550/ARXIV.1606.08415 (2016).

72. Chen, F., Datta, G., Kundu, S. & Beerel, P. Self-Attentive Pooling for Efficient Deep Learning. Preprint at 10.48550/ARXIV.2209.07659 (2022).

73. Su, J., et al. RoFormer: Enhanced Transformer with Rotary Position Embedding. Preprint at 10.48550/ARXIV.2104.09864 (2021).

74. Loshchilov, I. & Hutter, F. Decoupled Weight Decay Regularization. Preprint at 10.48550/ARXIV.1711.05101 (2017).

75. Cer, R. Z. et al. Non-B DB v2.0: a database of predicted non-B DNA-forming motifs and its associated tools. Nucleic Acids Res. 41, D94–D100 (2012).

76. Schreiber, J. tangermeme: A toolkit for understanding cis-regulatory logic using deep learning models. Preprint at 10.1101/2025.08.08.669296 (2025).

77. Schreiber, J. Tomtom-lite: Accelerating Tomtom enables large-scale and real-time motif similarity scoring. Preprint at 10.1101/2025.05.27.656386 (2025).

78. Kim, S. et al. Strelka2: fast and accurate calling of germline and somatic variants. Nat. Methods 15, 591–594 (2018).

79. Salvadores, M. & Supek, F. Cell cycle gene alterations associate with a redistribution of mutation risk across chromosomal domains in human cancers. *Nat*. Cancer 5, 330–346 (2024).

80. Derrien, T. et al. Fast Computation and Applications of Genome Mappability. PLoS ONE 7, e30377 (2012).

81. Díaz-Gay, M. et al. Assigning mutational signatures to individual samples and individual somatic mutations with SigProfilerAssignment. Bioinformatics 39, btad756 (2023).

82. Tate, J. G. et al. COSMIC: the Catalogue Of Somatic Mutations In Cancer. Nucleic Acids Res. 47, D941–D947 (2019).

