## Supplementary Figures S1-S22 for "Intrinsic DNA sequence determinants and tissue-specific regulation of human replication origins"

### Supplementary Materials for “Intrinsic DNA sequence determinants and tissue-specific regulation of human replication origins”

#### Tables:

- Table S1: Table lists each motif from STREME, along with p-values estimated from the logistic regression, ISM validations, along with the FDR corrections, and the first 3 matches of each STREME motif using TOMTOM (HOCOMOCO v11), if any.

#### Additional Files:

- File S1: STREME results / output html.
- File S2: TOMTOM Matches for motif 1-GGGAGGC[CT]GAGGC[AG].
- File S3: Positional Distributions of each motif using CentriMo on randomly selected 5,000 SNS-Seq peaks.
- File S4: Spacing analysis of 1-GGGAGGC[CT]GAGGC[AG] in combination with other motifs using CentriMo on randomly selected 5,000 SNS-Seq peaks.

#### Figures:

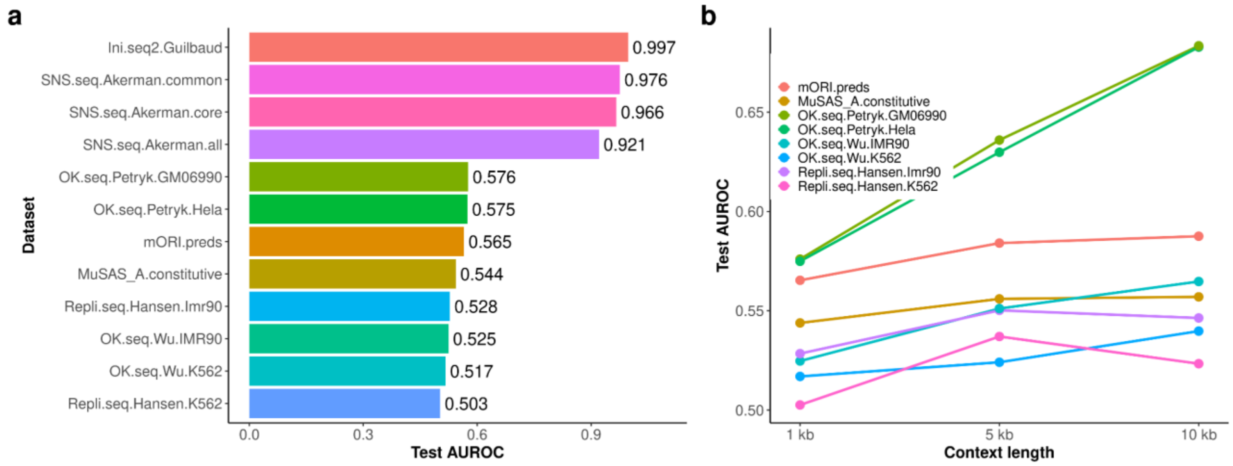


**Figure S1. The predictability of different replication origin datasets and context sizes.** **(a)** Hold out test performance (Area Under the Receiver Operating Characteristic, AUROC) scores of different versions of the ORIFormer architecture trained on different datasets of replication origins (with 1kb input length). **(b)** Performance comparison (AUROC) of models across different context lengths (1 kb, 5 kb, and 10 kb) to predict origin activity for selected datasets.

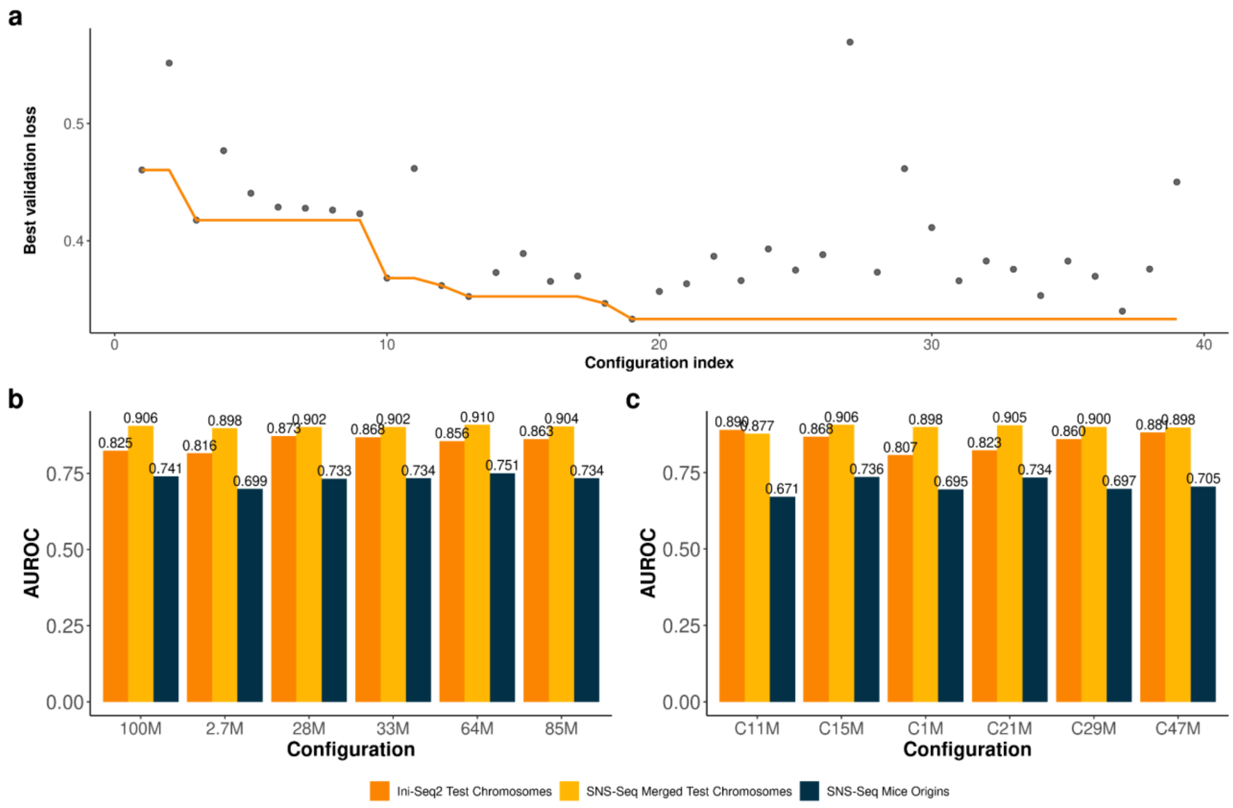


**Figure S2. Hyperparameter search for ORIFormer.**

**(a)** Panel a shows the hyperparameter optimization process, where each point represents a configuration's achieved validation loss, and the orange line tracks the best (minimum) validation loss achieved up to each configuration index. **(b - c)** Test performances (AUROC) for different parameter search configurations across four datasets after full training: Ini-Seq2 (left, orange bar), and SNS-Seq (middle, yellow bar) on Test Chromosomes, and SNS-Seq Mice Origins (rightmost bar, dark blue). Panel (b) shows top performing configurations of different sizes from the bayesian search, while panel (c) shows hand designed parameter configurations. Numbers in the names of models (x-axis) denote the number of parameters in millions (M).

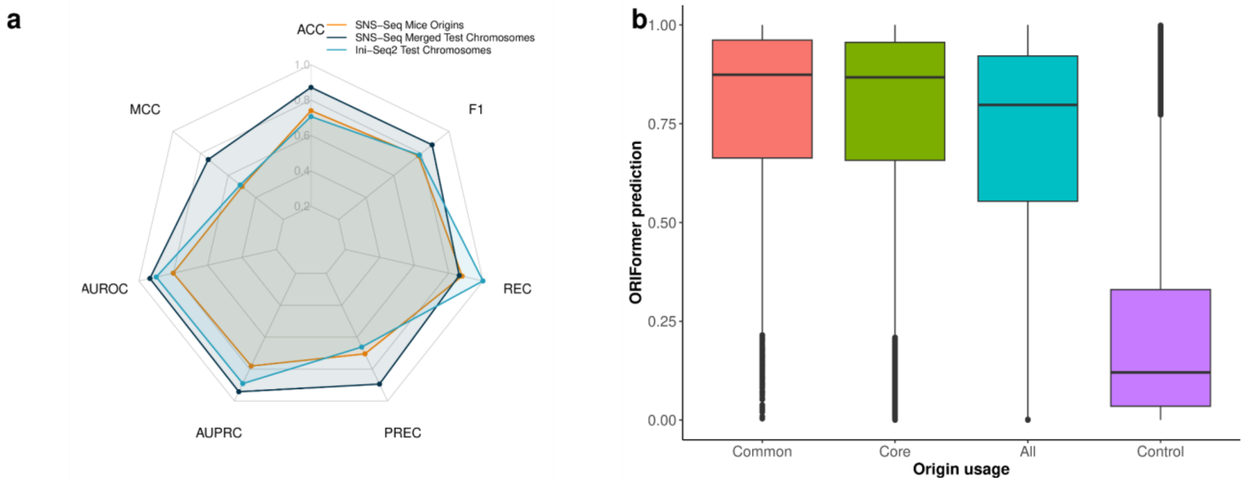


**Figure S3. ORIFormer generalises well and prioritises higher efficiency origins.**
**(a)** Radar plot comparing the performance of ORIFormer on three datasets (SNS-Seq Mice Origins, SNS-Seq, and Ini-Seq2 on Test Chromosomes) across several metrics, including AUROC, AUPRC, F1, Matthews correlation coefficient (MCC), accuracy (ACC), Precision (PREC), and Recall (REC). **(b)** ORIFormer predictions on SNS-Seq peaks of test chromosomes in different efficiency categories. Common (or constitutive) origins are ones that should be present in every cell type and cell cycle, while core origins are ones that harbour 80% of replication events (including common origins). All includes all previous types, plus stochastic origins which may remain dormant in some tissues and cell cycles. Controls are non-origin regions of the test chromosomes.

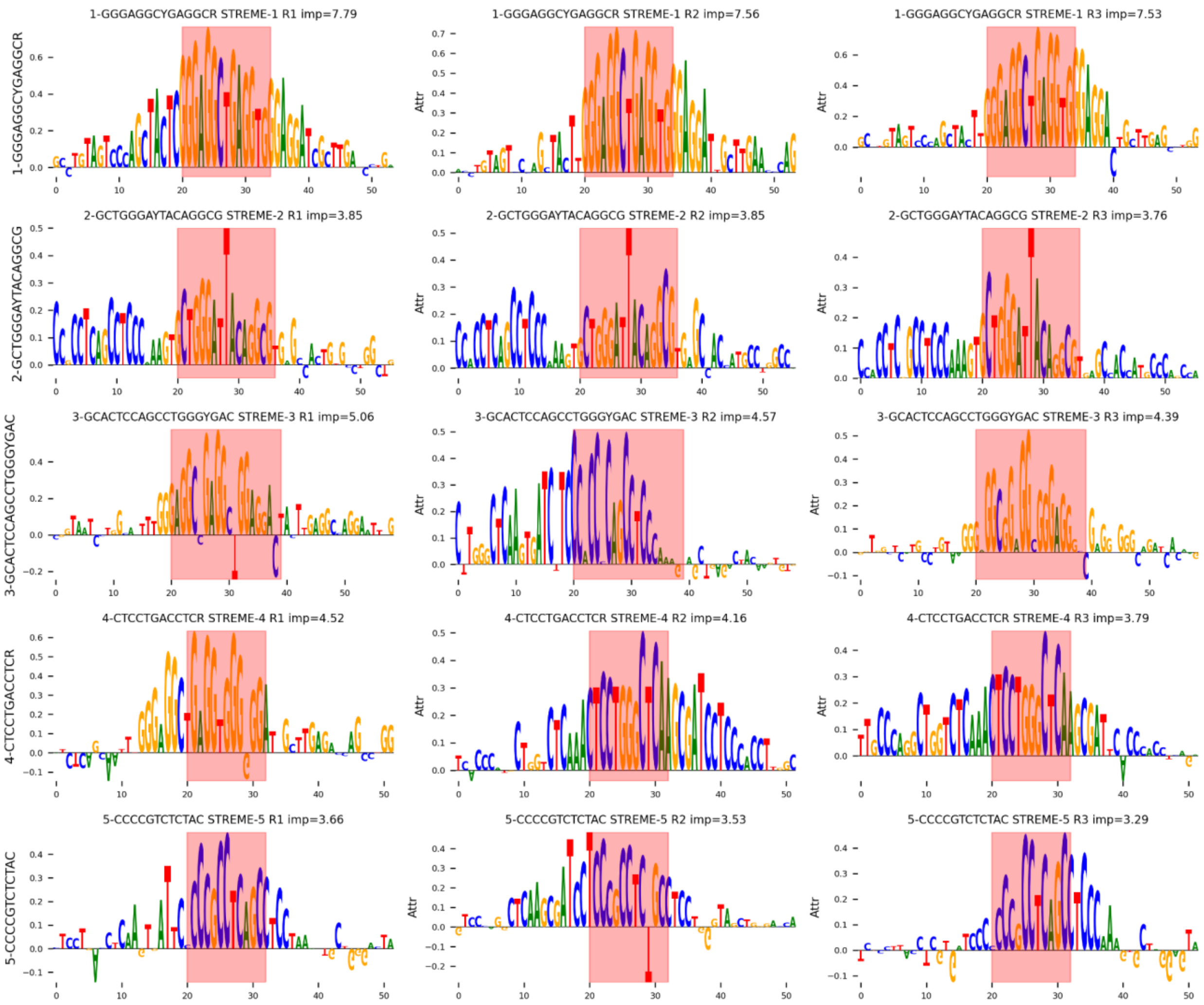


**Figure S4. Attribution-based sequence logos for top-ranking motif instances.** Each row represents a different transcription factor binding motif, with motif names shown on the left y-axis. For each motif, the three highest-importance instances (ranked R1, R2, R3) are displayed as sequence logos colored by integrated sequence mutation (ISM) attribution scores. Red shaded regions highlight the exact motif boundaries identified by FIMO scanning (threshold = 0.0001). Sequence context extends 20 base pairs upstream and downstream of each motif instance. Attribution values (Attr) on the y-axis represent the contribution of each nucleotide position to model predictions, with positive values indicating positions that increase replication origin probability. The importance score (imp) shown in each subplot title represents the sum of attribution values across the motif region. Only motifs with identified instances above the significance threshold are shown.

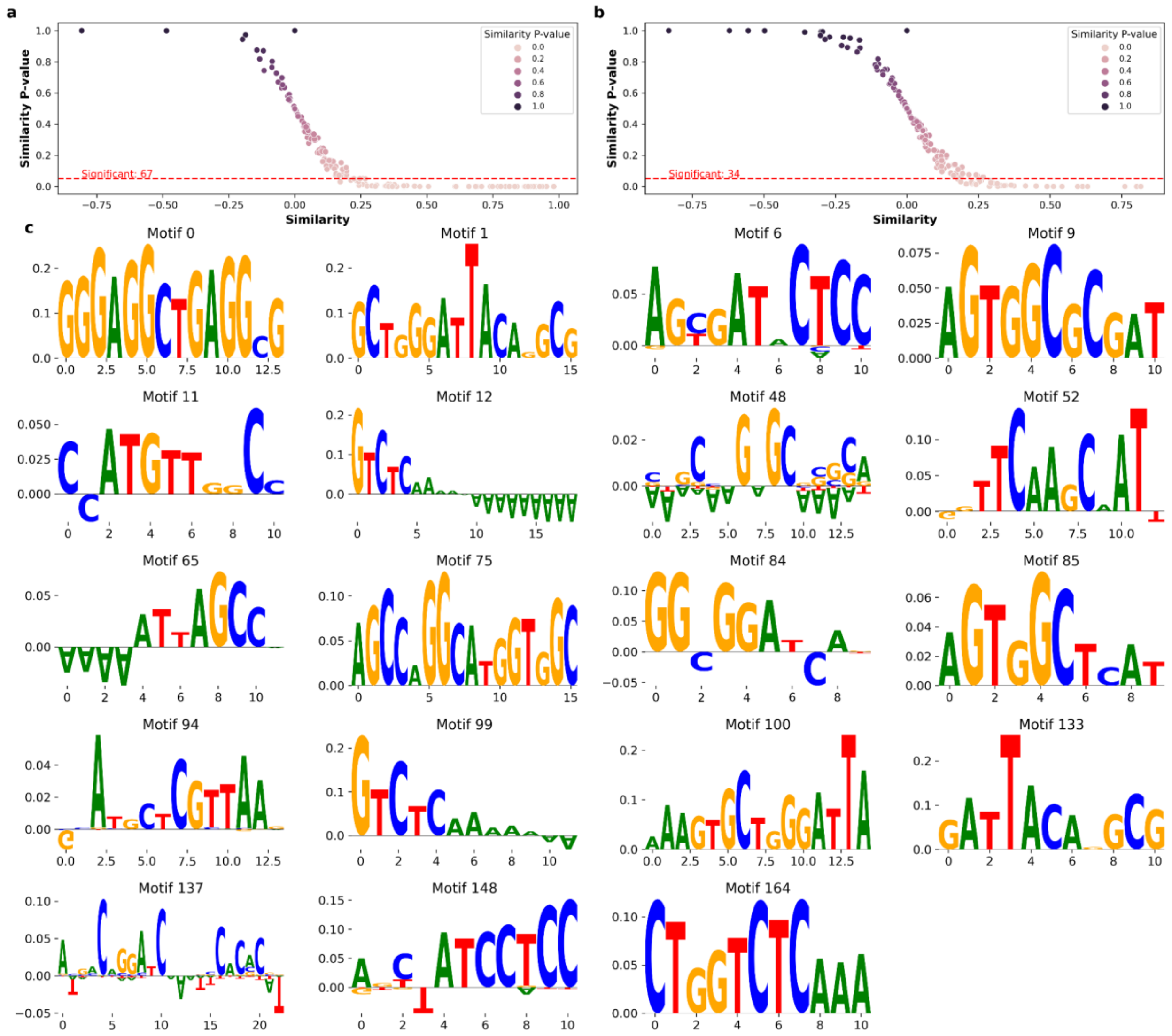


**Figure S5. Validating Motifs using *in silico mutagenesis.***
**(a - b)** Similarity scores of averaged attribution maps of each motif with its original PWMs (x-axis) vs empirical p-value, calculated from a baseline of random PWMs' similarity scores (see Methods). Analysis on top ranked 25,000 test SNS-Seq origins (a), and top ranked 10,000 Ini-Seq2 origins (b). Threshold at p = 0.05 is highlighted. **(c)** Attribution plots of the 19 motifs validating (FDR < 10%) on both SNS-Seq and Ini-Seq2 origins. Nucleotides are scaled by the maximum change in ORIFormer predictions, if they are mutated.

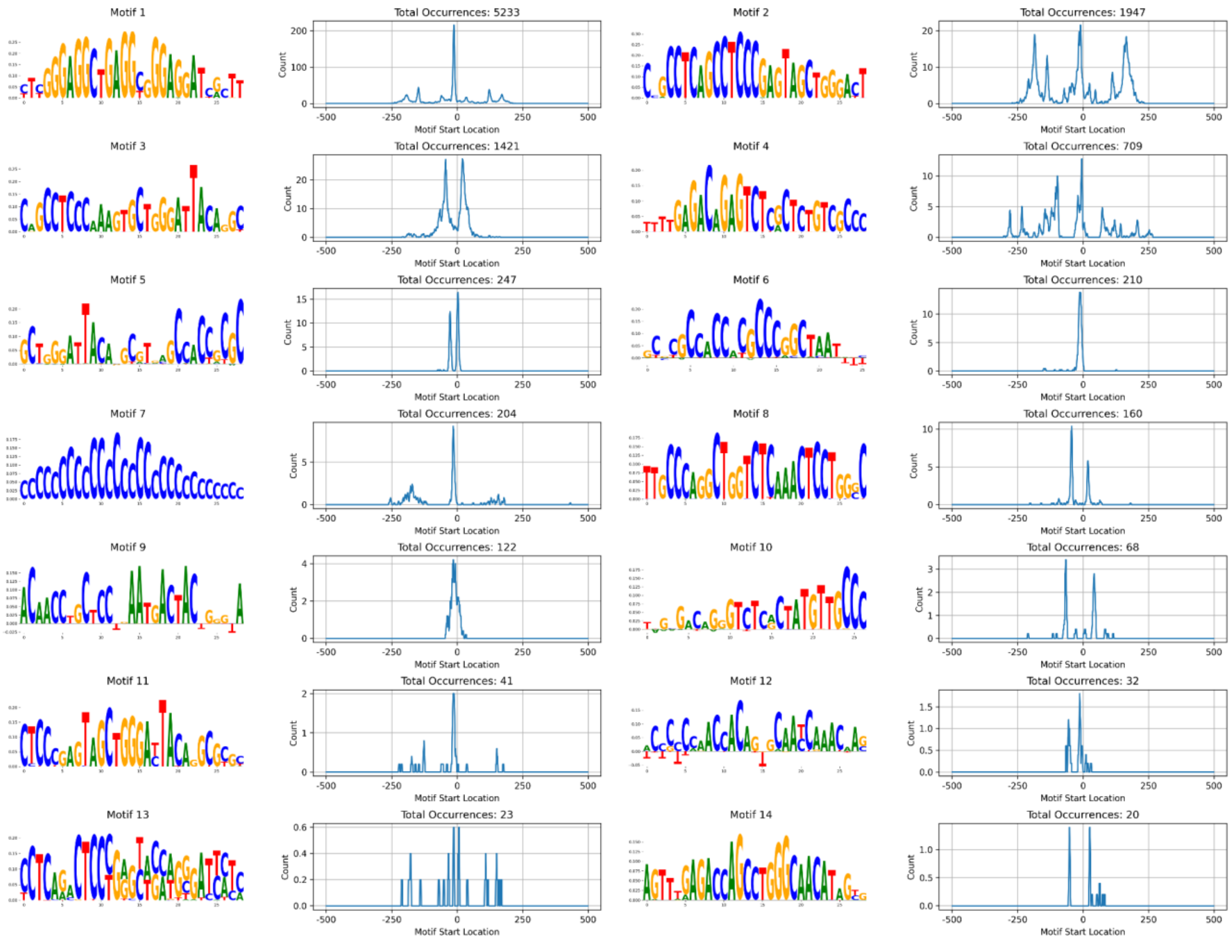


**Figure S6. Modisco motifs and their positional distributions.**
Motif logos of motifs obtained from TF-MoDISco using ISM attribution maps from the top ranked 10,000 test SNS-Seq peaks. Nucleotides are scaled by their importance to ORIFormer. Next to each motif, the number of instances (y-axis) at each position (x-axis) relative to the origin (center) is shown. Total number of instances displayed at the top of each positional curve.

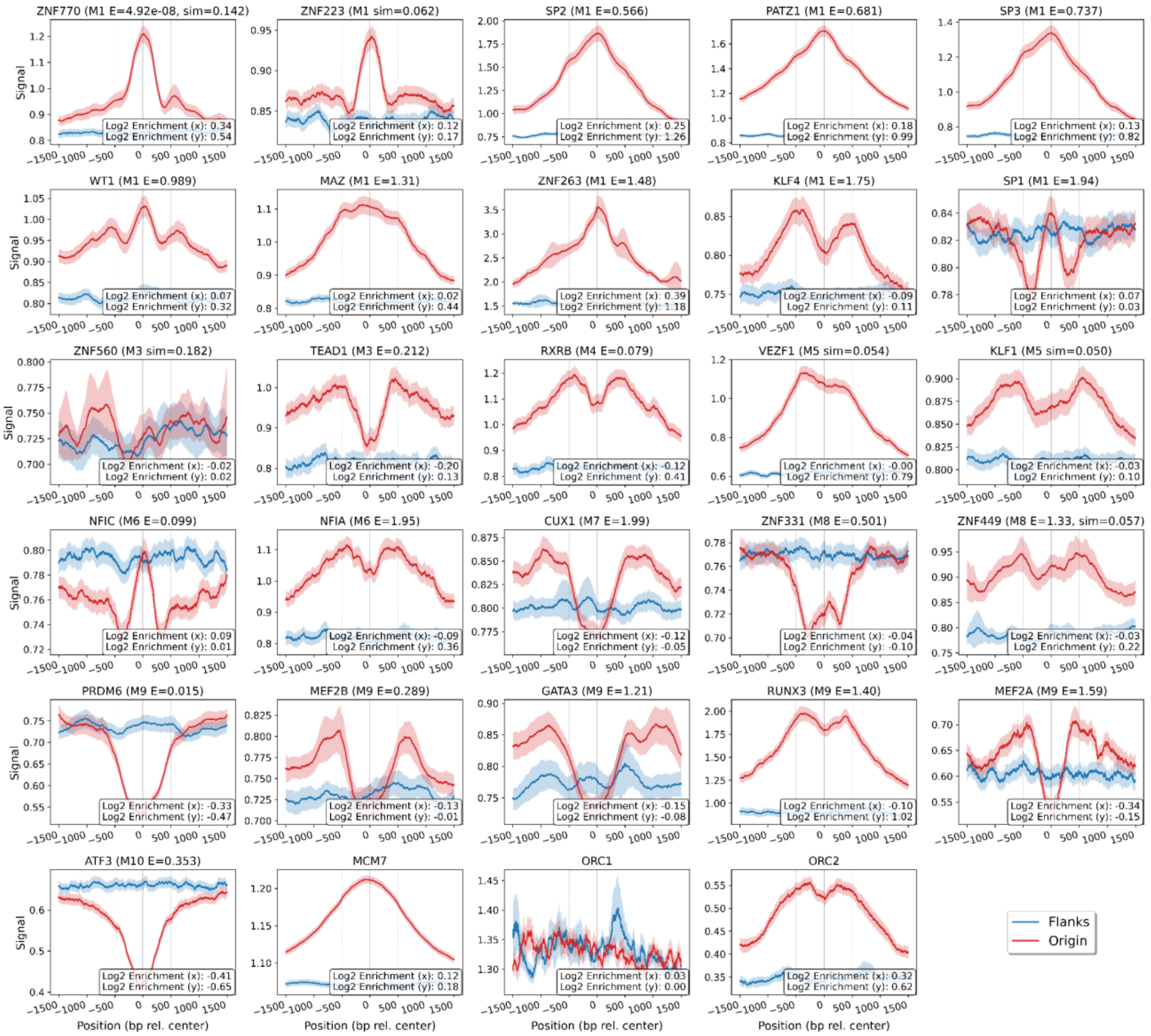


**Figure S7. ChIP-seq signal enrichment profiles of TF binding sites at replication origins**. Average ChIP-seq signal across 3 kb windows centered on predicted replication origins (red) versus control flanking regions (blue). Each panel shows a different transcription factor with motif cluster (M1-M10) and significance values. Shaded areas represent 95% confidence intervals. Log2 enrichment values shown: (x) origin center vs. left flank at -500bps, (y) origin center vs. control center. Notable enrichment observed for pre-replication complex components (MCM7, ORC1, ORC2), for these the flank is taken at distance -1500bps.

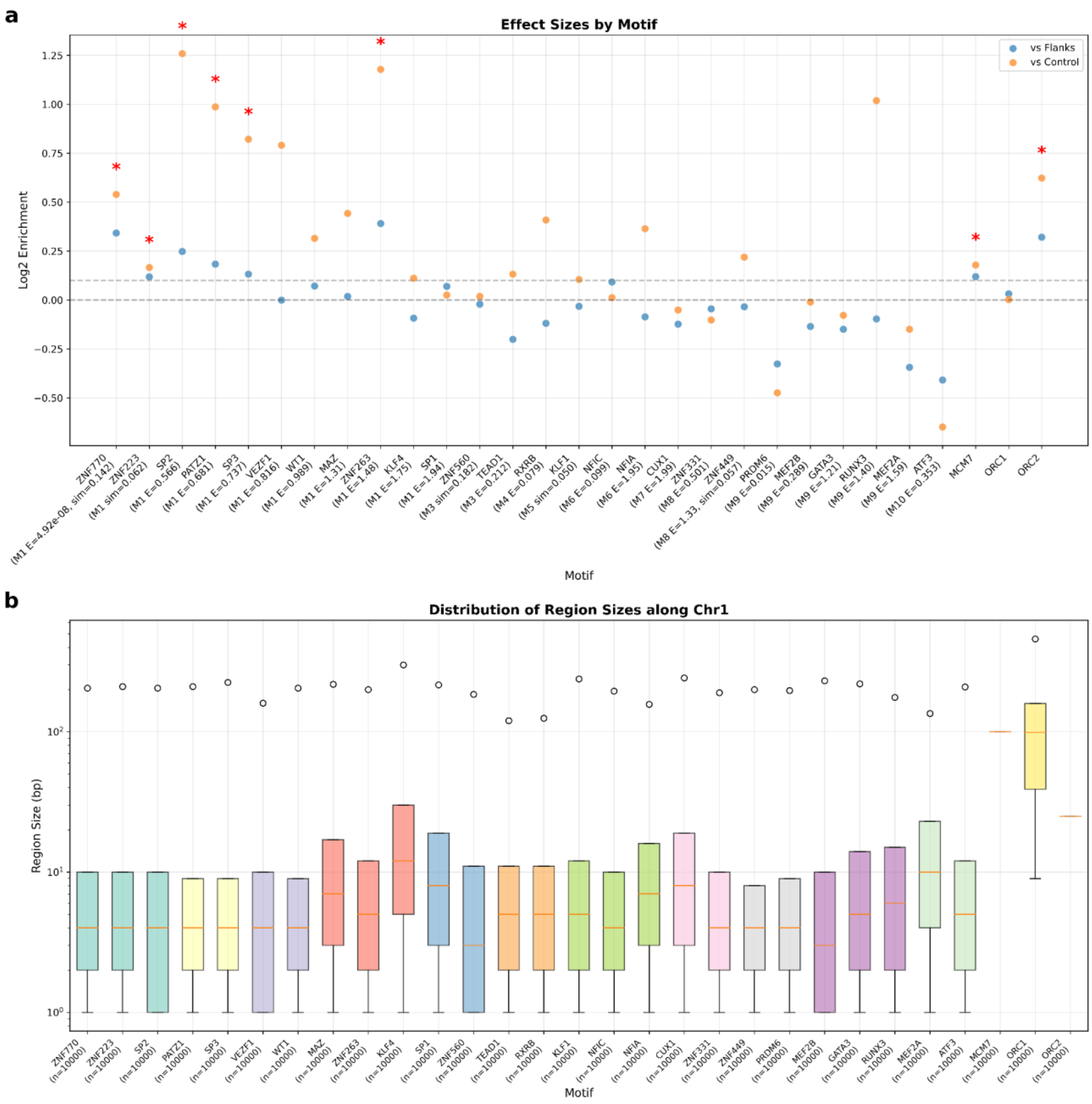


**Figure S8. ChIP-seq enrichment analysis and region size distributions**.
**(a)** Log2 enrichment values for transcription factor binding motifs, comparing enrichment within origins (center vs -500 bp, orange) and enrichment of origin centers relative to flanking regions (vs Flanks, blue). Motifs are ordered by similarity groups (M1-M10) with E-values and similarity scores shown, and are denoted by stars (*) if both enrichment values are above 0.1. **(b)** Distribution of sampled region sizes along chromosome 1 for each TF ChIP signal, displayed as box plots showing median, quartiles, and range of binding site region lengths. Sample sizes (n) are indicated for each.

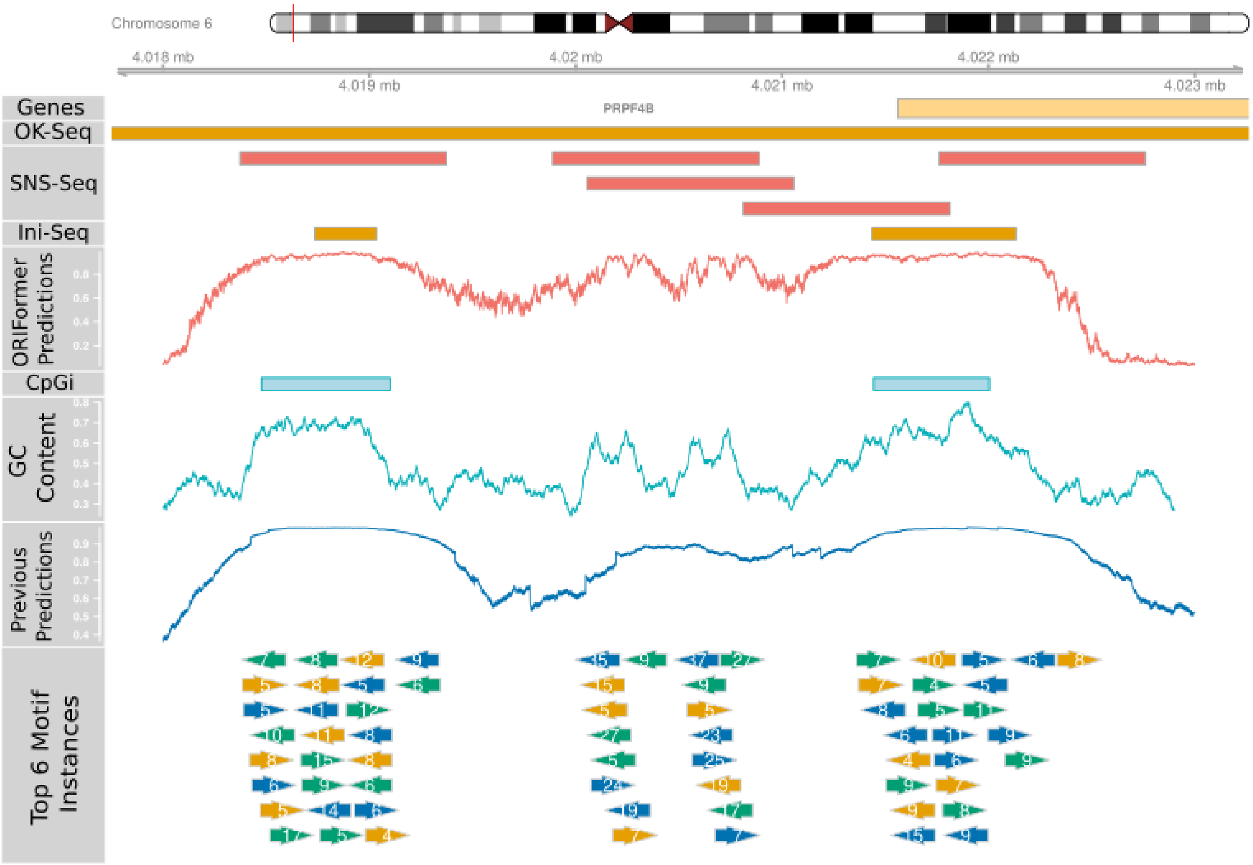


**Figure S9. Replication origin locus near the PRPF4B / PRP4K gene.**
Replication loci at the promoter region of PRPF4B / PRP4K transcript (light yellow bar, top) near at chr6 p25.2. The region is within an initiation zone (HR Repli-Seq H1) denoted by orange bars below the gene transcript, and contains multiple initiation sites (SNS-Seq in red, Ini-Seq2 in orange below). ORIFormer predictions, previous model predictions are shown in red and dark blue, respectively at basepair resolution. GC content and CpG islands are highlighted in light blue. Instances of top motifs, their matching score, and orientation are shown in coloured arrows (motif instances displayed from the 9th row of arrows onwards were omitted).

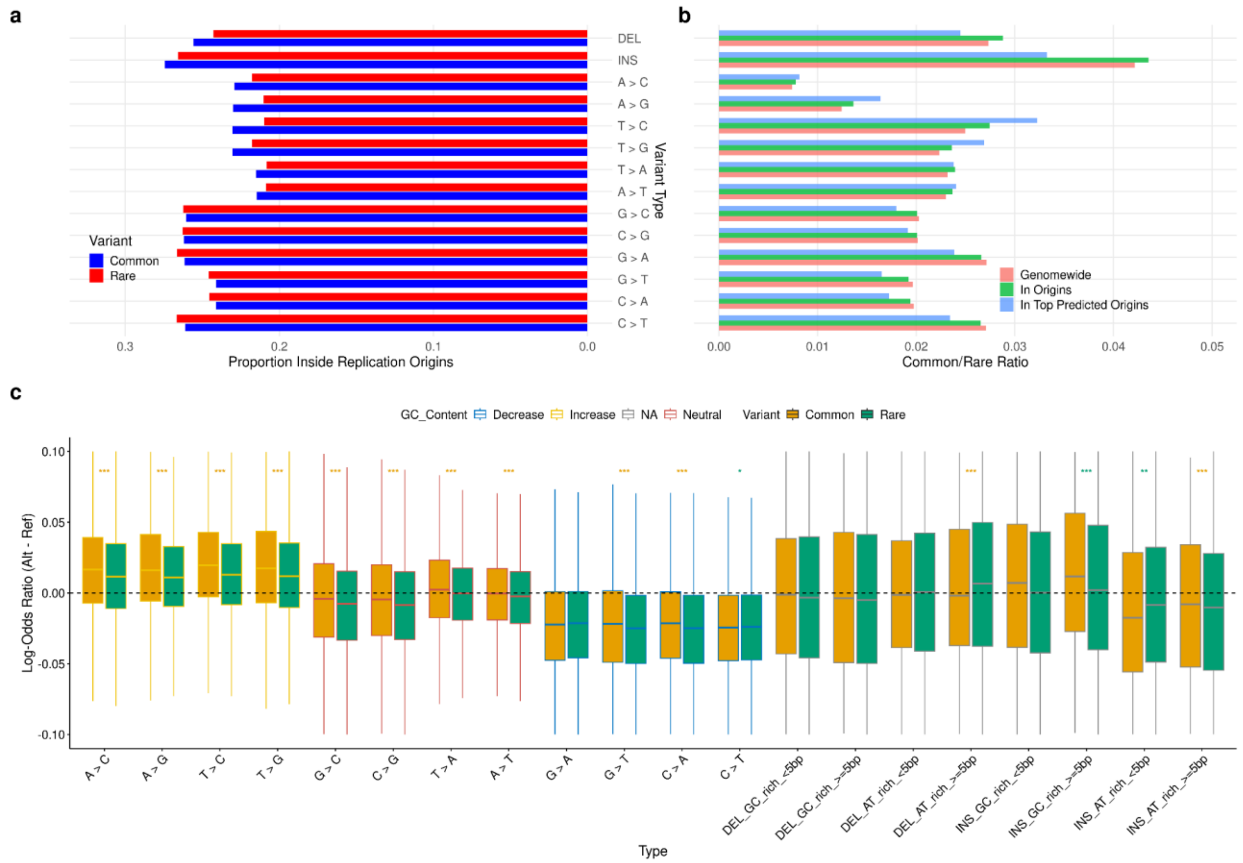


**Figure S10. Genomic variants in the population at replication origins.**

**(a)** Ratio of the number of variants inside SNS-Seq origins and all variants of that type. **(b)** Ratios of common to rare variants of the same type genomewide, in origins, and in top predicted origins highlight G/C > N variants to be less enriched in common variants, suggesting that they are not fixed in the population. **(c)** Predicting the effects of variants of each type inside top predicted replication origins on test chromosomes via ORIFormer, using the reference and the alternative alleles. The log-odds ratio (difference in logits alt - ref) is shown on the y-axis, while variant type on the x-axis. The effect of a variant on GC content is highlighted by the boxplot outlines, while the fill colour denotes occurrence. Outliers omitted, and the IQR is shortened to 10% for visual clarity. Significance levels from Wilcoxon tests are shown between each box, indicating via colour the group with higher mean value.

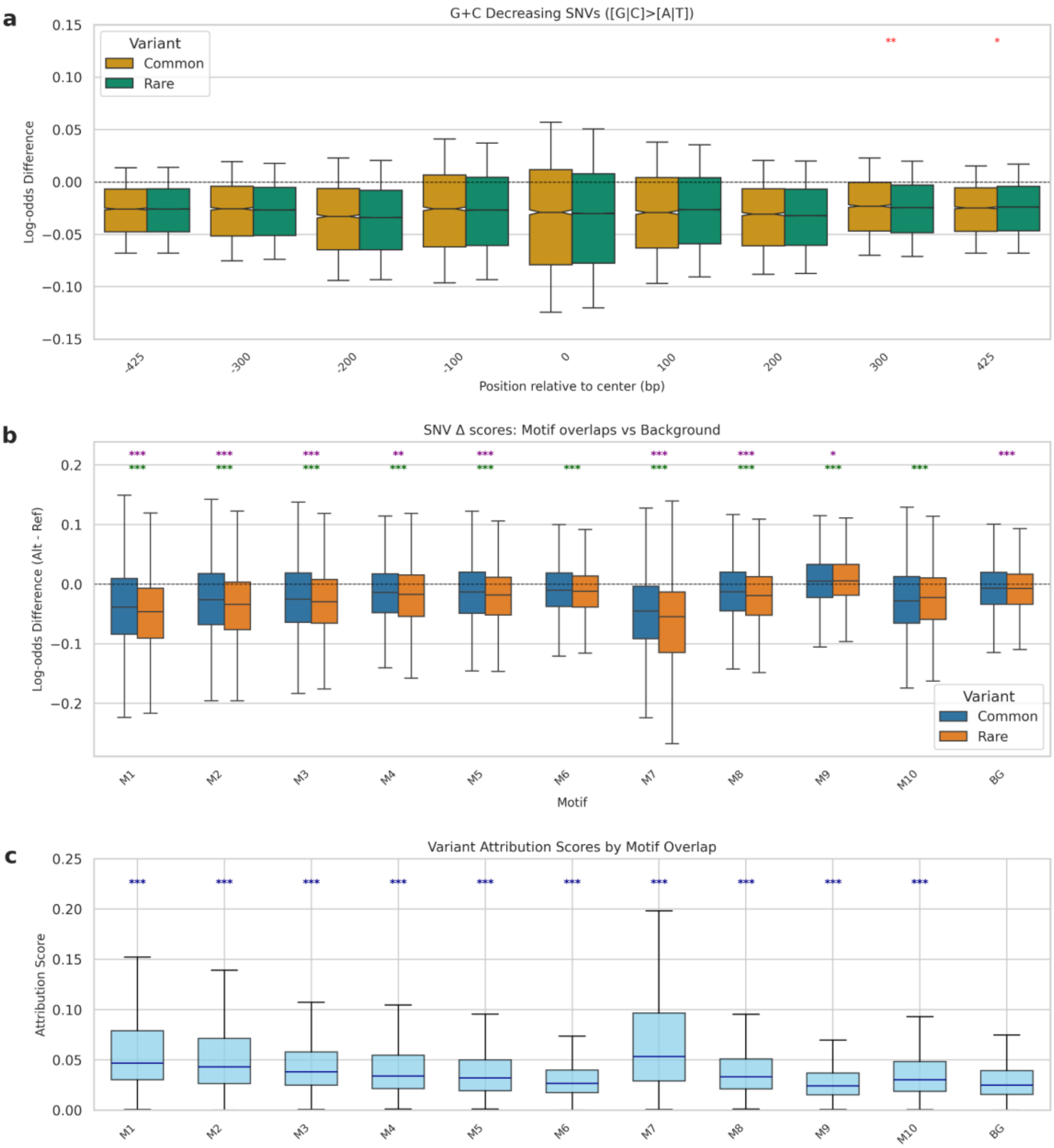


**Figure S11. Detailed analysis of germline variant effects on replication origin predictions. (a)** Position-dependent mutational effects on origin activity. Box plots show median log-odds differences (Alt - Ref) for SNVs binned by relative position within 1kb origin regions, stratified by mutation category (G+C increasing/neutral) and variant frequency (common/rare). Notched boxes indicate 95% confidence intervals. Whiskers shortened to 0.5 x IQR. **(b)** Distribution of log-odds difference scores for SNVs overlapping top ORIFormer motifs compared to background variants not overlapping any motif. Boxplots show median and quartiles, with statistical significance indicated for comparisons between Common and Rare variants (purple stars) and between each motif and background (green stars). **(c)** Distribution of hypothetical attribution scores for SNVs at the same loci of all variants combined. Statistical significance between each motif's attribution values and the background attribution values are shown by stars.

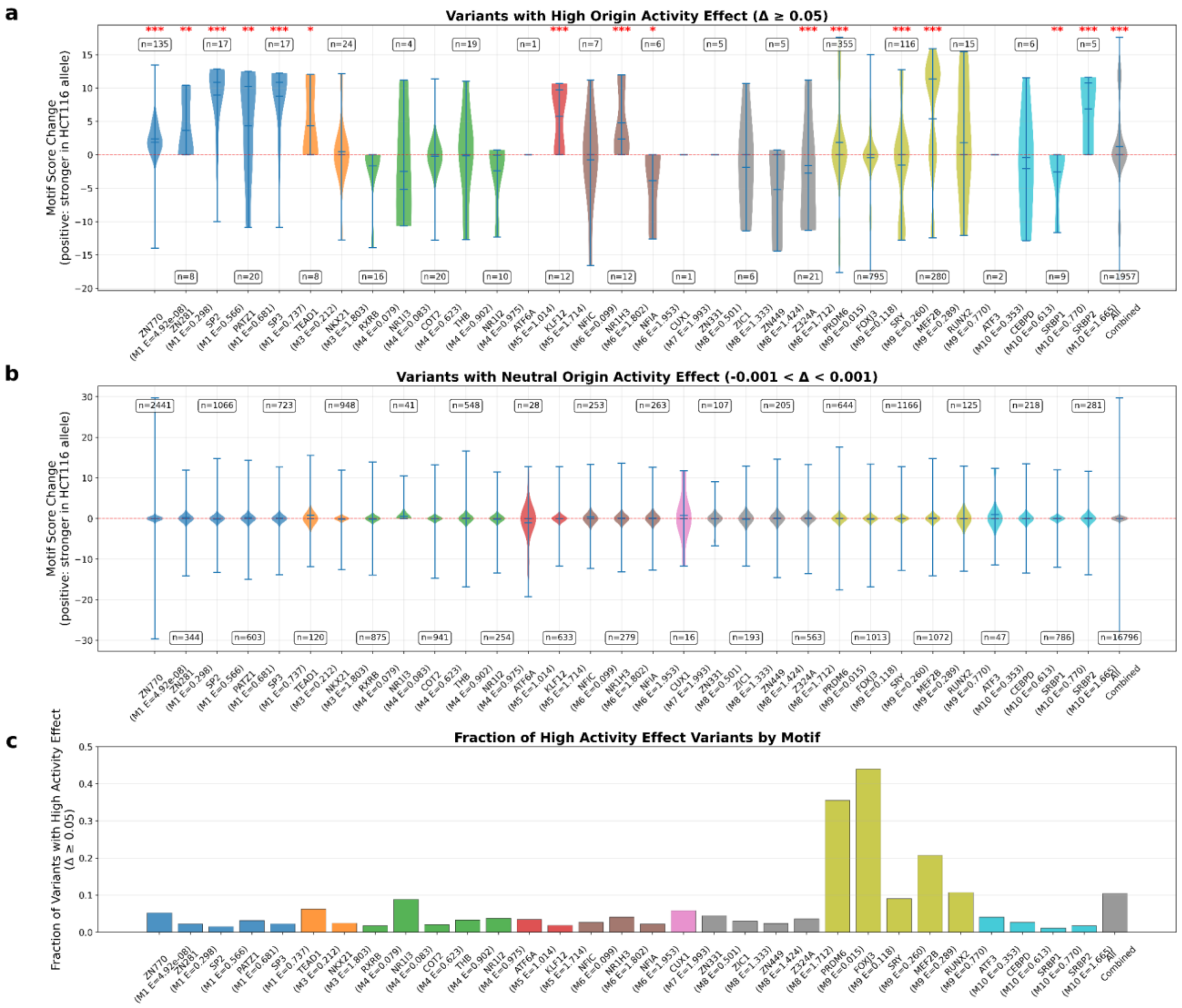


**Figure S12. Motif binding site disruption analysis.**

**(a)** Motif score changes in variants with high origin activity effect (Δ ≥ 0.05). Violin plots show motif binding score changes for variants that significantly increase origin activity. Each violin represents a transcription factor motif, colored by ORIFormer motif (M1-M10). Positive values indicate stronger binding in HCT116 alternative alleles. Sample sizes (n) and significance stars from Mann-Whitney U tests are shown above violins. **(b)** Motif score changes in variants with neutral origin activity effect (-0.001 < Δ < 0.001). Control comparison showing motif binding score changes for variants with minimal origin activity impact. Same color scheme and ordering as panel (a). **(c)** Fraction of high activity effect variants by motif. Proportion of high effect variants (Δ ≥ 0.05) relative to total variants for each motif. Higher fractions indicate motifs preferentially disrupted in variants strongly affecting origin activity. Motif annotations show ORIFormer motif identity (M1-M10) and TOMTOM E-values for top 5 transcription factor matches. Analysis excludes variants overlapping MCF7 origins and uses ±25bp windows around variants.

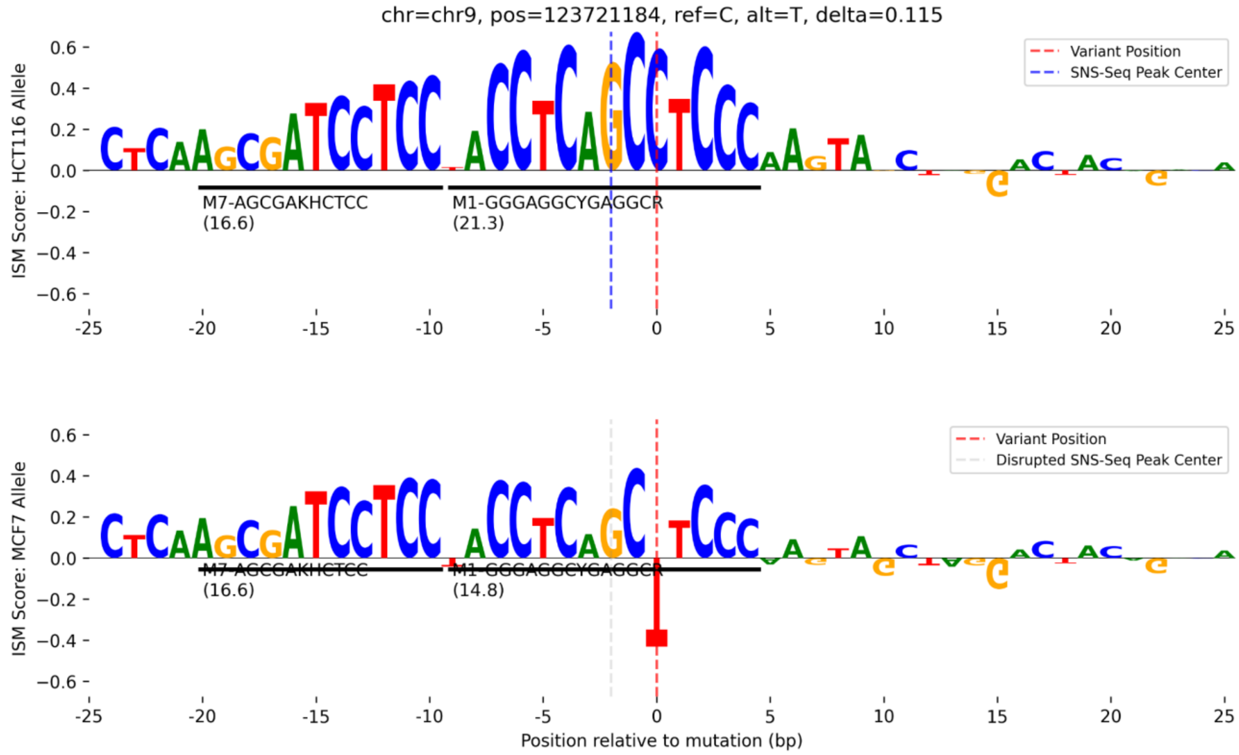


**Figure S13. In silico saturation mutagenesis of origin-disrupting C>T variant in MCF7.** Upper and lower panels show the ISM plots of reference and alternative alleles, respectively, with motif annotations overlaid. Red dashed lines mark variant positions, while blue and gray dashed lines show the disrupted SNS-seq peak centers. Motif matches via FIMO to ORIFormer motifs M1-M10 are also shown.

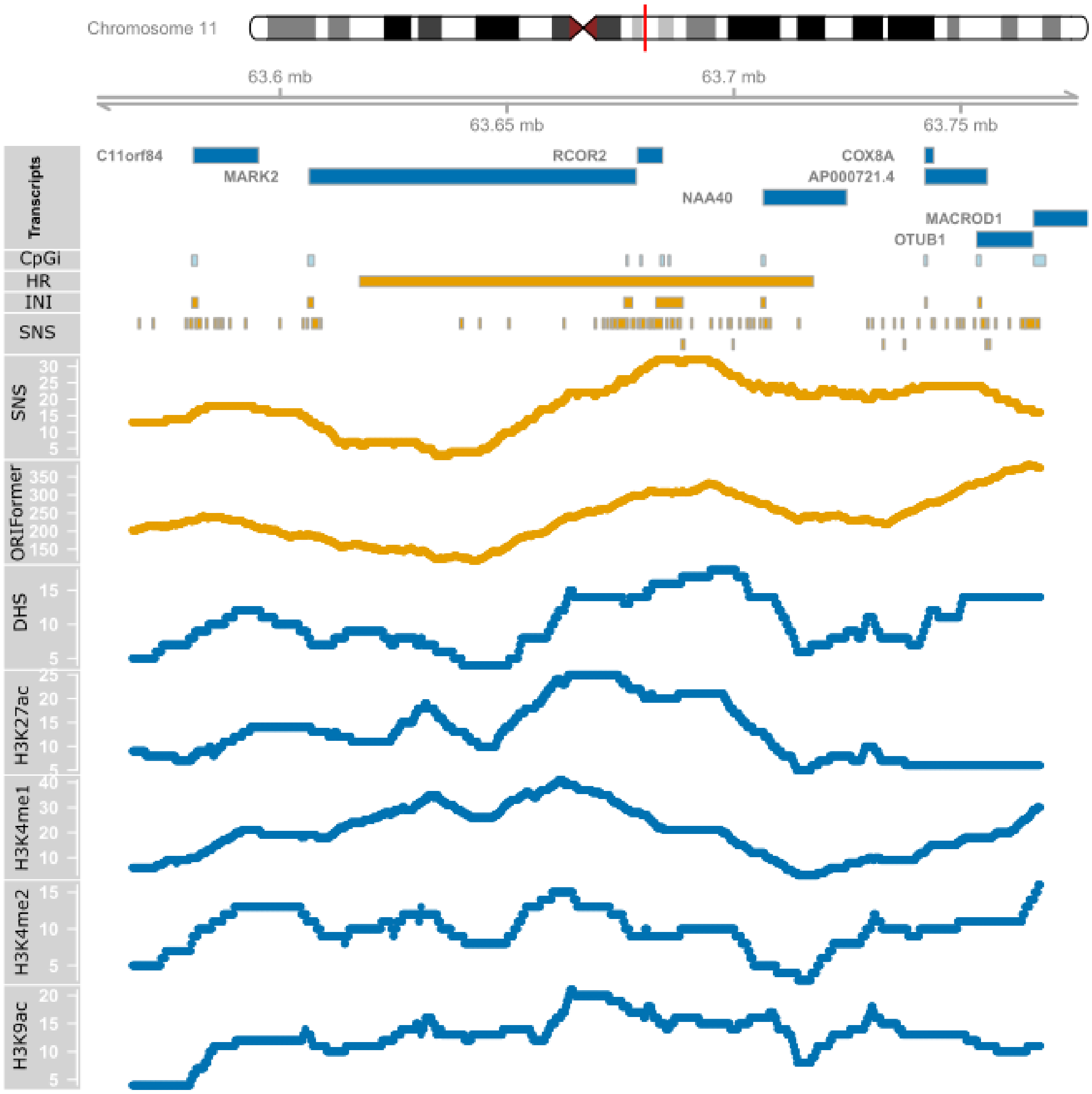


**Figure S14. Example initiation zone shows elevated density of initiation sites and marks.** The highlighted genomic region at chr 11q13.1 contains an HR Repli-Seq initiation zone (in H1). The region contains a number of SNS-Seq peaks and Ini-Seq2 peaks, namely at high density at the IZ. It also contains multiple gene bodies and CpG islands. SNS-Seq Count/Density at each position, as well as total signal per epigenetic track are shown in yellow and blue, respectively

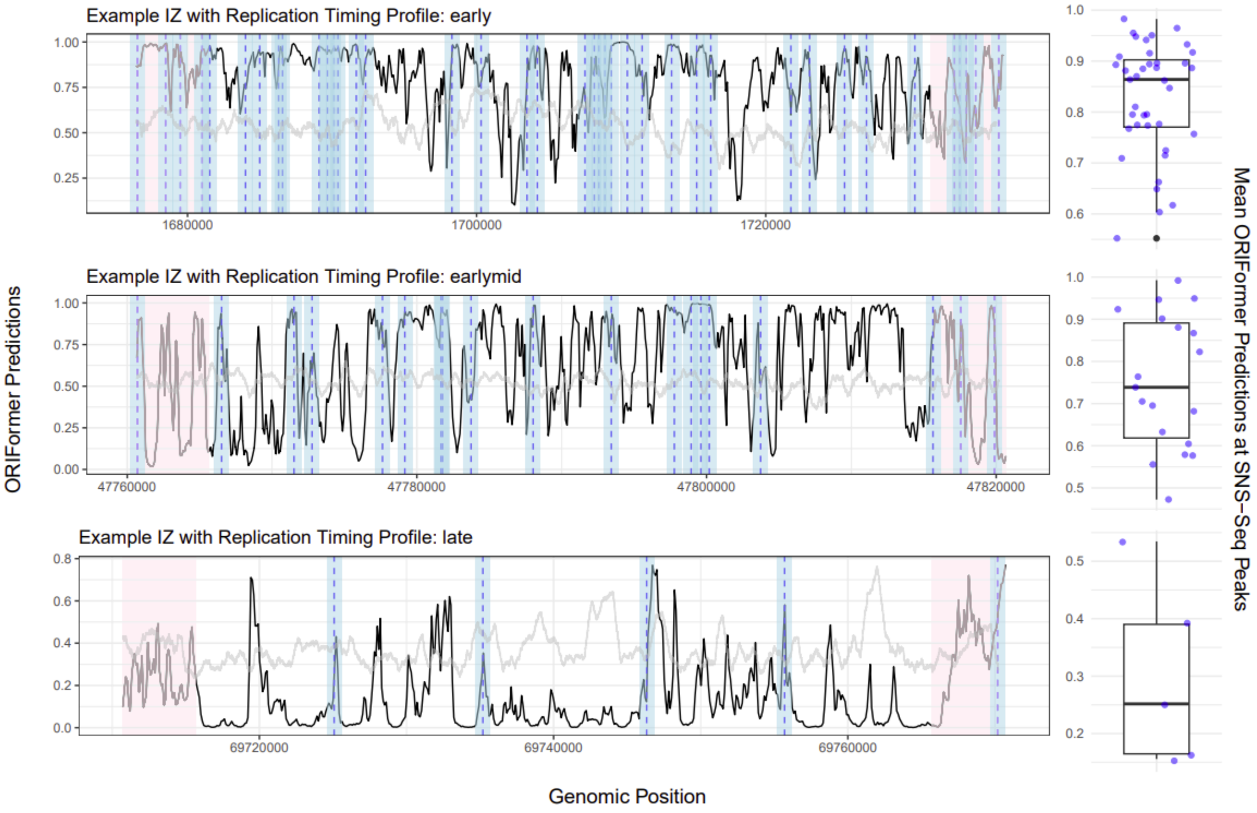


**Figure S15. Early replicating initiation zones contain more higher efficiency origins.**
3 example HR Repli-Seq initiation zones (H1) of different replication profiles are shown, regions between the shaded red region, which highlight the flanking 5kbs at each side of the IZ. Black lines show ORIFormer predictions at each genomic position, while light gray lines show the GC content. Shaded blue areas denote SNS-Seq peaks of 1kb, centered at the horizontal dotted lines. Boxplots next to each example IZ denote the distribution of ORIFormer predictions at measured SNS-Seq peaks within each IZ.

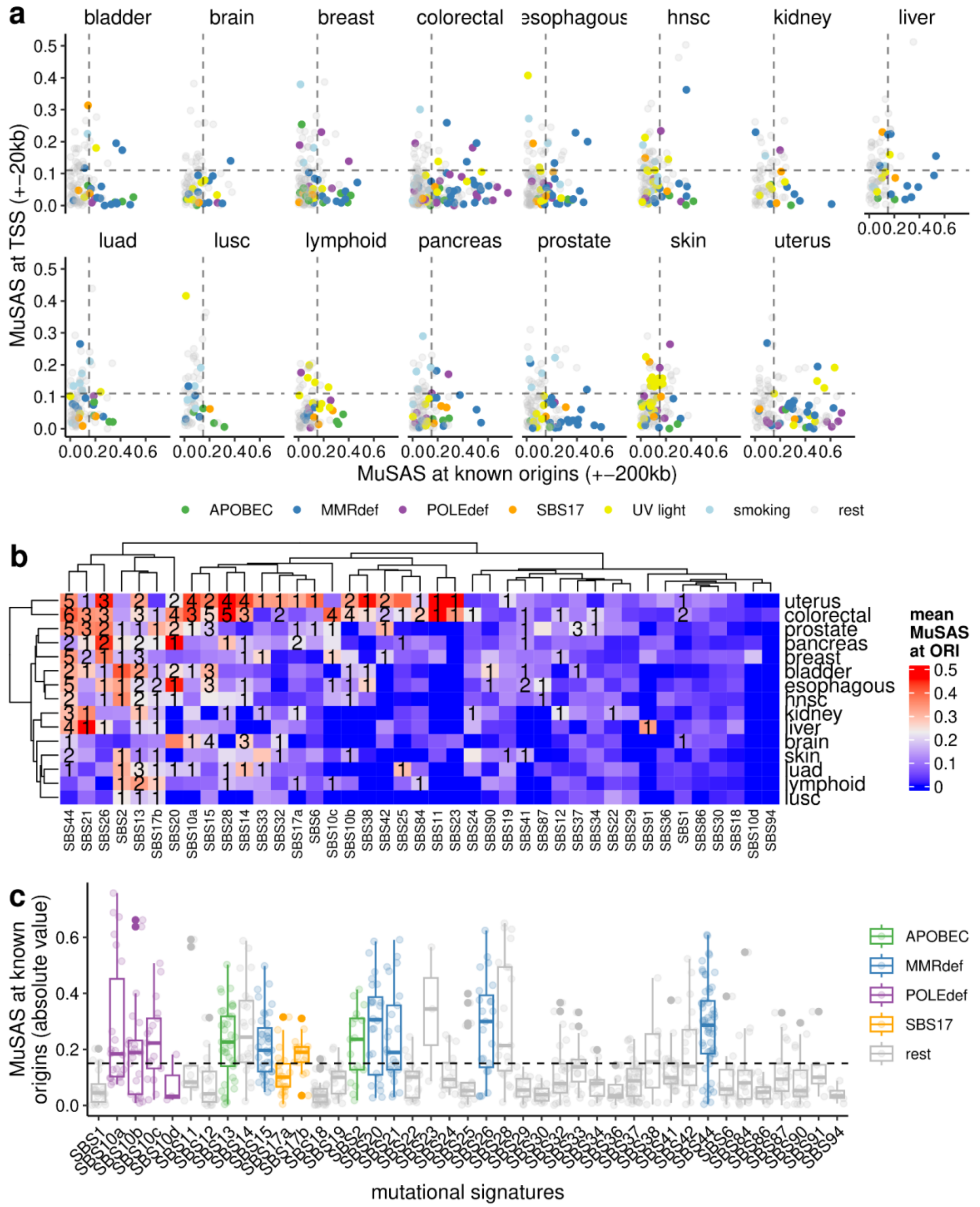


**Figure S16. Mutational strand asymmetries across tissue-mutational signature-substitution combinations. (a)** MuSAS score at origin loci vs at TSS loci for all signature-tissue-mutation_type combinations separated by cancer type. **(b)** Mean MuSAS at ORI per cancer type and per signature. The number represents the number of mutation types that pass the threshold (from 1 to maximum 6). **(c)** Summary of MuSAS score at ORI per mutational signature. Each dot is a combination of cancer type and mutation type. The groups with robust strand asymmetry are grouped by colors according to their mechanisms.

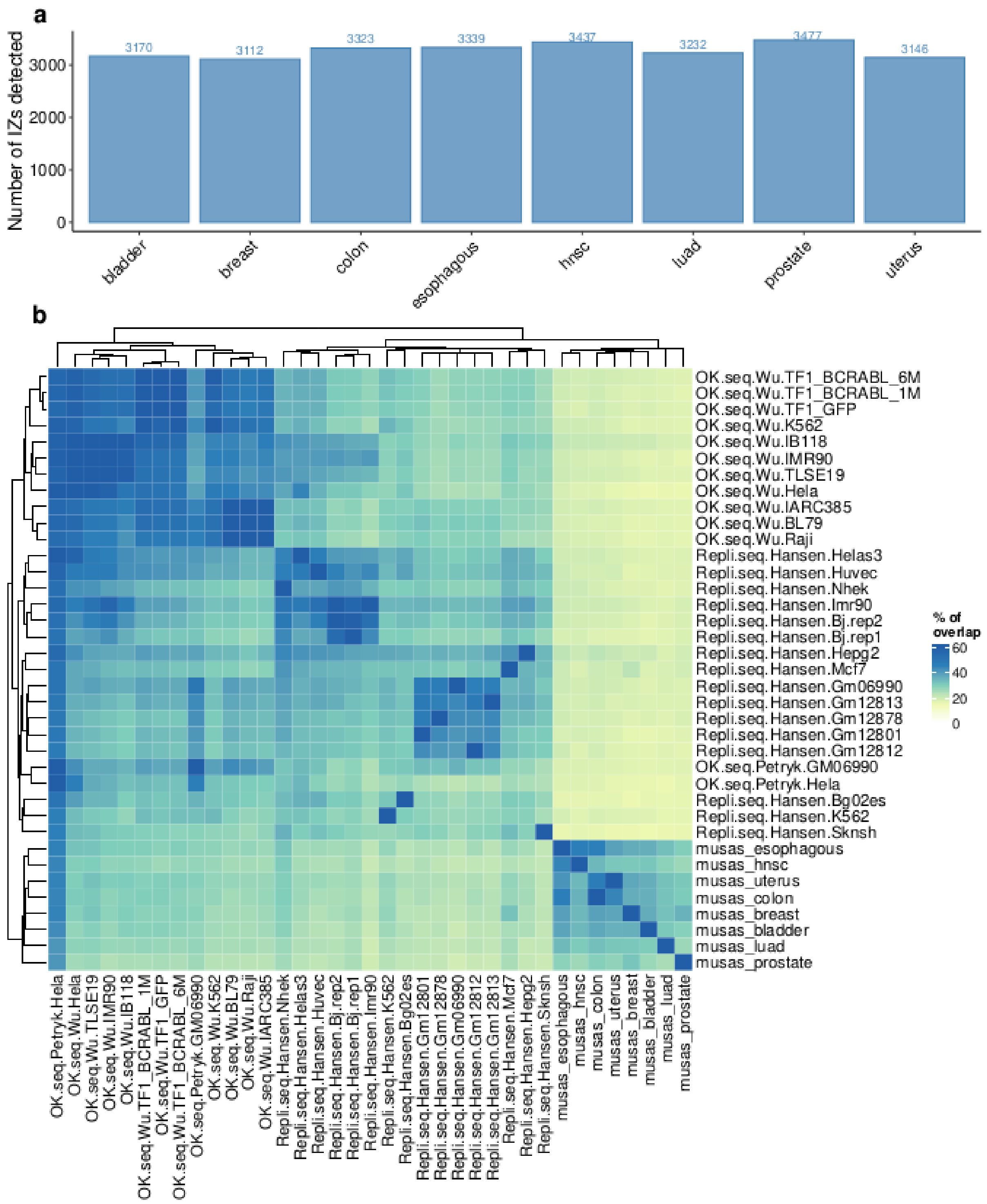


**Figure S17**. **MuSAS-IZ agreement with experimental datasets**. **(a)** Number of MuSAS-IZs detected in each tissue. **(b)** Percentage (%) of origins that overlap between different datasets (pairwise combinations). To equalize all the origin sizes, each origin is set as the middle point on the window +-30kb.

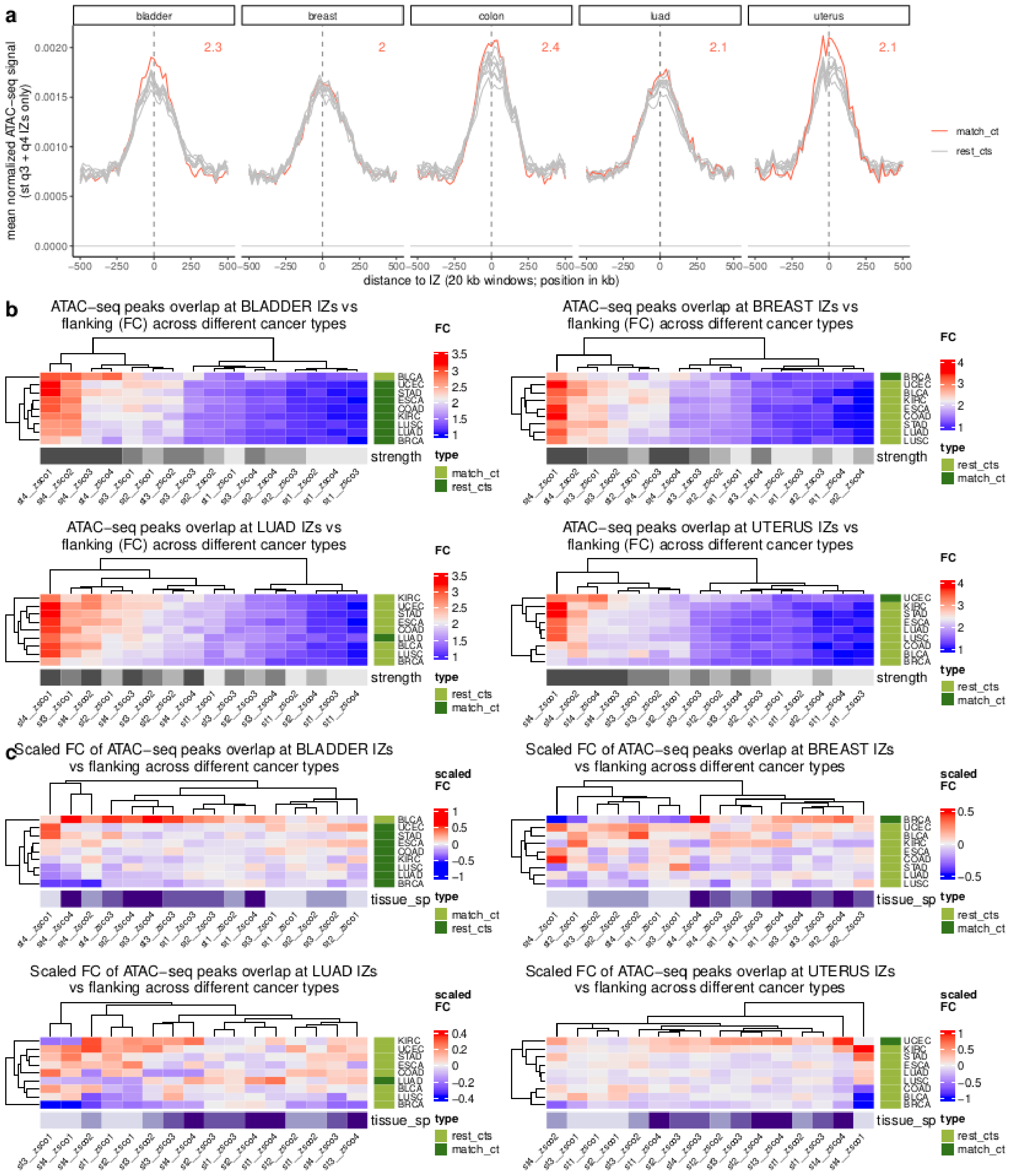


**Figure S18**. **Tissue-specific replication origins localize to regions of tissue-specific chromatin accessibility. (a)** Normalized frequency of ATAC-seq peaks and flanking windows across the different cancer types (matching cancer type in red) at the MuSAS-IZs for bladder, breast, colon, luad and uterus. The IZs are grouped in 4 categories according to their strength (the highest strength, quartile Q3+Q4, is shown here). Numbers within each panel show the fold-change (FC) between ATAC-seq intensity at the IZ centre normalized to the flanks for the matching tissue. **(b)** Fold-change (FC) between normalized frequency of ATAC-seq peaks at IZ (MuSAS peak) centre, versus flanking regions, for MuSAS-IZs subdivided into 4 quartiles of strength (st1-st4) and 4 quartiles of tissue-specificity (ts1-ts4), total 16 categories. Each row represents the chromatin accessibility (ATAC-seq) of a different tissue, positively associating with origin strength)**.** **(c)** Same as panel **b**, but FC is scaled across columns of the heatmap, adjusting for the effect of the IZ strength, emphasizing that ATAC-seq has the strongest signal in matching tissue**.**

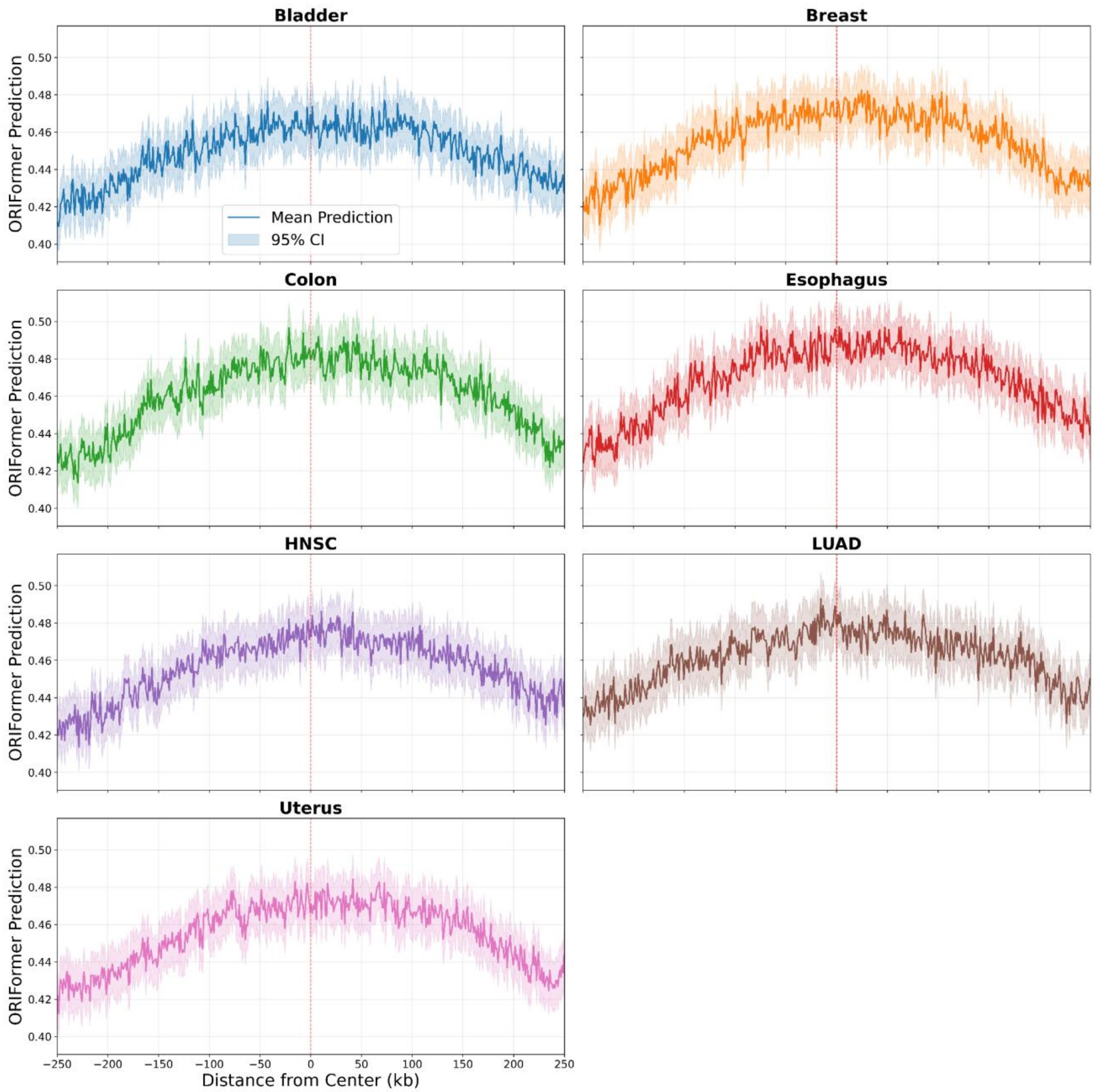


**Figure S19: ORIFormer predictions nearby MuSAS peak centers in multiple tissues.** ORIFormer prediction scores plotted as a function of distance from MuSAS peak centers across eight cancer types. Lines represent mean predictions with shaded regions indicating 95% confidence intervals. Red dashed lines mark the MuSAS peak center (0 kb). Predictions show consistent enrichment at peak centers across all tissues, demonstrating that ORIFormer-predicted replication initiation sites correspond to experimentally-determined mutation strand asymmetry signatures.

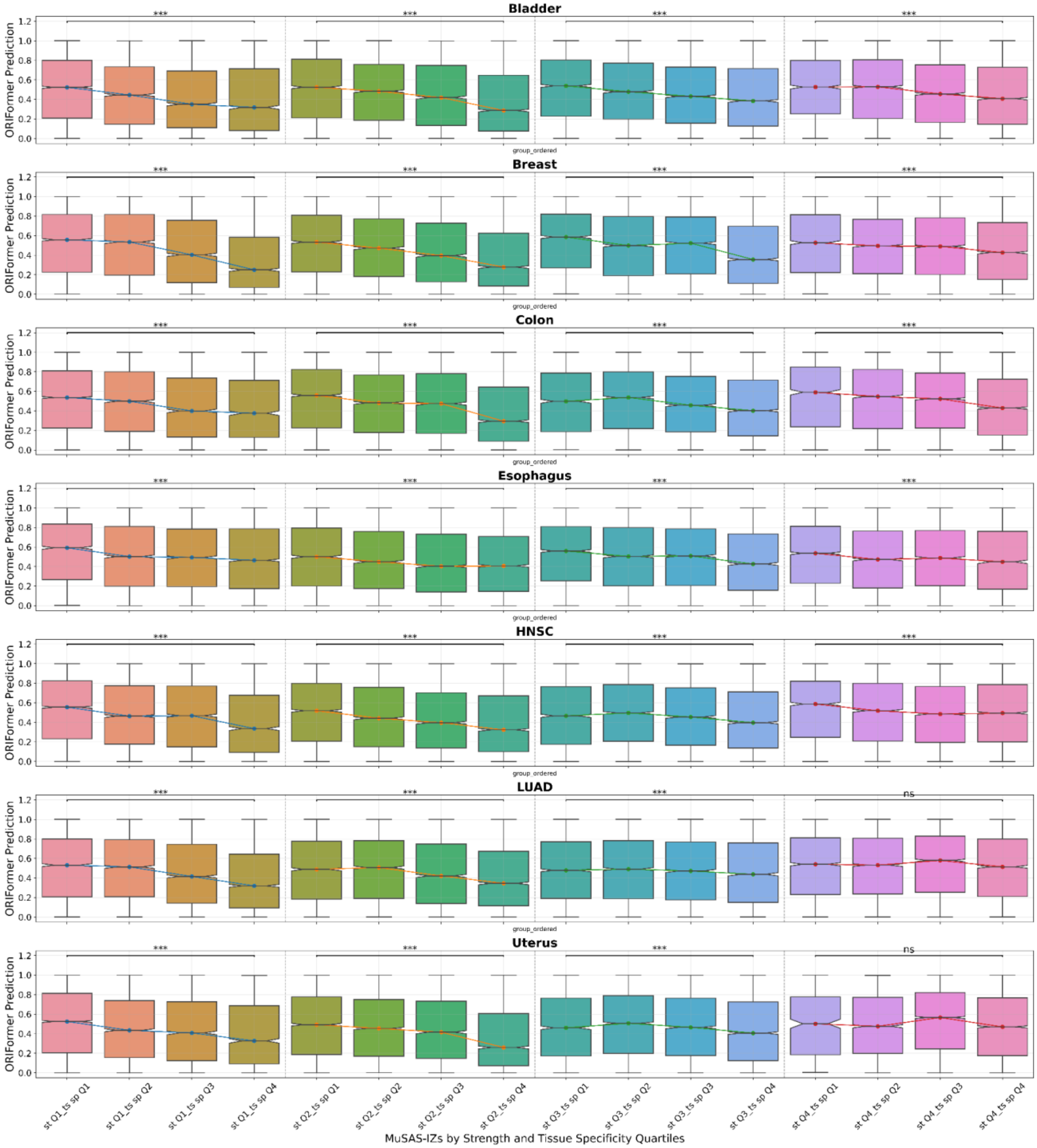


**Figure S20: ORIFormer predictions stratified by MuSAS peak strength across tissues.** Box plots showing ORIFormer prediction scores grouped by MuSAS peak strength (st Q1 to Q4) and tissue specificity (ts sp Q1 to Q4) across multiple tissues. Within each strength category, lines connect median values across z-score levels. Gray dashed vertical lines separate different strength categories. Statistical comparisons between zsco1 and zsco4 within each strength level are indicated by horizontal bars with significance stars (*** p<0.001, ** p<0.01, * p<0.05, ns: not significant). Higher strength peaks consistently show elevated ORIFormer predictions.

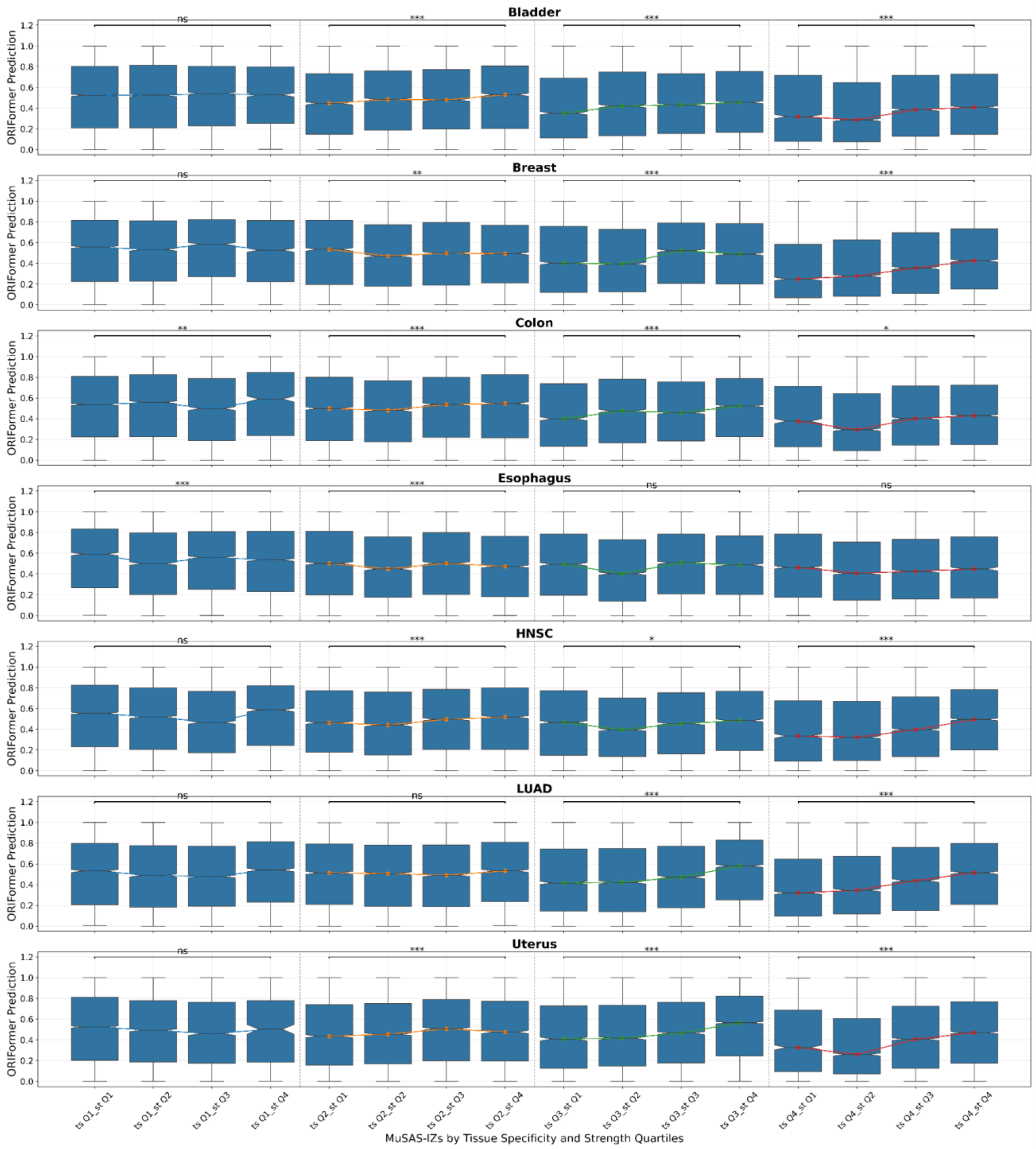


**Figure S21: ORIFormer predictions stratified by MuSAS peak z-score across tissues.** Box plots showing ORIFormer prediction scores grouped by MuSAS peak tissue specificity score (ts Q1 to Q4) and mutation strength (st Q1 to Q4) across multiple tissues. Within each z-score category, lines connect median values across strength levels. Gray dashed vertical lines separate different z-score categories. Statistical comparisons between st1 and st4 within each z-score level are indicated by horizontal bars with significance stars (*** $p<0.001$, ** $p<0.01$, * $p<0.05$, ns: not significant). Higher z-score peaks show increased ORIFormer predictions, confirming the model's ability to identify stronger replication origins.

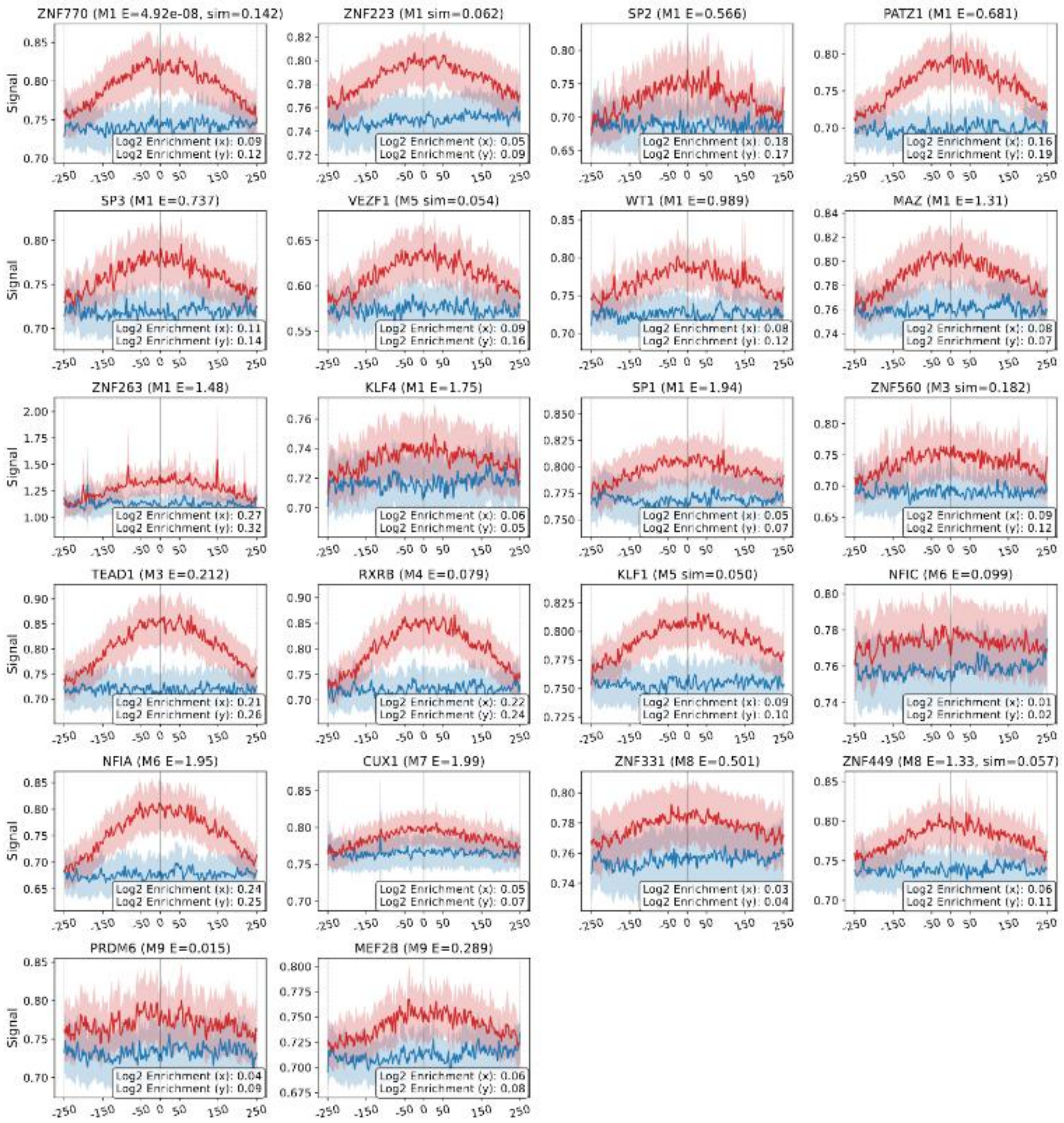


**Figure S22: ORIFormer predictions stratified by MuSAS peak z-score across tissues.** Smoothed ChIP-seq signal profiles for ORIFomer motif M1-M10 associated factors, as well as replication initiation associated factors MCM7, and ORC2, across MuSAS-detected IZs in colon cancer. Enrichment can be observed at MuSAS IZ centers (“Origin”) compared to non-origin, flanking genomic regions (“Negative”). Signals averaged over 500kb windows ± SEM shown. Log2 enrichment of ORIFormer signals stated in the plot indicate: (x) IZ center vs. upstream edge within IZs; (y) IZ center vs. non-origin, flanking regions center. Signal smoothed using a Gaussian filter with sigma 4, and window size 5kb.
