## Supplementary File S1-S4 for "Intrinsic DNA sequence determinants and tissue-specific regulation of human replication origins"

### STREME

Sensitive, Thorough, Rapid, Enriched Motif Elicitation

For further information on how to interpret these results please access https://meme-suite.org/meme/doc/streme.html.
To get a copy of the MEME software please access https://meme-suite.org.

If you use STREME in your research, please cite the following paper:
Timothy L. Bailey, "STREME: accurate and versatile sequence motif discovery", *Bioinformatics*, Mar. 24, 2021. [full text]

Discovered Motifs | Inputs & Settings | Program Information | Motifs in MEME Text Format ? | Matching Sequences ? | Matching Sites ? | Results in XML Format ?

#### Discovered Motifs

Next Top

| Motif ? | Logo ? | RC Logo ? | P-value ? | E-value ? | Sites ? | More ? | Submit/Download ? | Positional Distribution ? | Matches per Sequence ? |
|---|---|---|---|---|---|---|---|---|---|
| 1-GGGAGGCYGAGGCR | | | 5.0e-924 | 9.0e-922 | 23109 (92.4%) | ↧ | ⇢ | | |
| 2-GCTGGGAYTACAGGCG | | | 2.7e-818 | 4.8e-816 | 23514 (94.1%) | ↧ | ⇢ | | |
| 3-GCACTCCAGCCTGGGYGAC | | | 4.9e-814 | 8.8e-812 | 23192 (92.8%) | ↧ | ⇢ | | |
| 4-CTCCTGACCTCR | | | 9.6e-655 | 1.7e-652 | 22165 (88.7%) | ↧ | ⇢ | | |
| 5-CCCCGTCTCTAC | | | 1.3e-582 | 2.3e-580 | 21012 (84.0%) | ↧ | ⇢ | | |
| 6-GCAGTGAGCCGAG | | | 5.0e-523 | 9.0e-521 | 20033 (80.1%) | ↧ | ⇢ | | |
| 7-AGCGAKHCTCC | | | 3.8e-433 | 6.9e-431 | 19402 (77.6%) | ↧ | ⇢ | | |
| 8-CVAGACCAKCCY | | | 7.0e-402 | 1.3e-399 | 18447 (73.8%) | ↧ | ⇢ | | |
| 9-AAAAATACAAAAAWT | | | 4.8e-344 | 8.6e-342 | 12348 (49.4%) | ↧ | ⇢ | | |
| 10-AGTGGCGYGAT | | | 1.3e-295 | 2.2e-293 | 12360 (49.4%) | ↧ | ⇢ | | |
| 11-CCACCVCVCC | | | 2.3e-290 | 4.2e-288 | 17034 (68.1%) | ↧ | ⇢ | | |
| 12-CCRTGTTGGCC | | | 1.5e-235 | 2.6e-233 | 15946 (63.8%) | ↧ | ⇢ | | |
| 13-GTCTCAAAAAAAAAAAAAA | | | 2.7e-225 | 4.9e-223 | 9792 (39.2%) | ↧ | ⇢ | | |
| 14-CAGGAGCWGGKKS | | | 1.2e-169 | 2.2e-167 | 14008 (56.0%) | ↧ | ⇢ | | |
| 15-CTGGGWGSYS | | | 1.3e-126 | 2.4e-124 | 15322 (61.3%) | ↧ | ⇢ | | |
| 16-GGCAGATCACC | | | 9.4e-108 | 1.7e-105 | 4868 (19.5%) | ↧ | ⇢ | | |
| 17-RGGCWGAGGS | | | 1.1e-097 | 1.9e-095 | 13828 (55.3%) | ↧ | ⇢ | | |
| 18-CCAGGCCCHN | | | 1.1e-053 | 1.9e-051 | 10591 (42.4%) | ↧ | ⇢ | | |
| 19-GGGAGGAGRS | | | 2.2e-052 | 4.0e-050 | 11181 (44.7%) | ↧ | ⇢ | | |
| 20-CCCWGSCWGGG | | | 1.3e-045 | 2.4e-043 | 9625 (38.5%) | ↧ | ⇢ | | |
| 21-CAGCCCCWSC | | | 6.9e-045 | 1.2e-042 | 10074 (40.3%) | ↧ | ⇢ | | |

**Stopped because 3 consecutive motifs exceeded the p-value threshold (0.05).**
**STREME ran for 62351.38 seconds.**

| Motif | Logo | RC Logo | P-value | E-value | Sites | More | Submit/Download | Positional Distribution | Matches per Sequence |
|---|---|---|---|---|---|---|---|---|---|
| 22-AAAAAAAAAAAAAA | | | 5.3e-043 | 9.5e-041 | 4501 (18.0%) | | | | |
| 23-AGGYGGATCA | | | 4.5e-042 | 8.1e-040 | 1777 (7.1%) | | | | |
| 24-CYCAGCTCTGC | | | 1.2e-040 | 2.2e-038 | 5856 (23.4%) | | | | |
| 25-GTTATTTCTTGCCTTCTGCTAGCTTTTGAA | | | 2.7e-040 | 4.8e-038 | 1153 (4.6%) | | | | |
| 26-AATCCTGAGTTCTAGTTTGAT | | | 5.5e-040 | 9.9e-038 | 1139 (4.6%) | | | | |
| 27-AAGGCAGAAATAAAGATGTTCTTT | | | 1.1e-039 | 2.0e-037 | 1152 (4.6%) | | | | |
| 28-CAGTTTCCATGTAGCTGAGCGGTTT | | | 1.1e-039 | 2.0e-037 | 1138 (4.6%) | | | | |
| 29-AGATTCTGGTATGTTGTGTCTTTG | | | 1.1e-039 | 2.0e-037 | 1139 (4.6%) | | | | |
| 30-AAAGTCTCCCATTATTAWTGTGT | | | 2.3e-039 | 4.2e-037 | 1141 (4.6%) | | | | |
| 31-ATGTGGTCAATTTTGGAATA | | | 2.3e-039 | 4.2e-037 | 1118 (4.5%) | | | | |
| 32-CAGAGCAGAACTGAAGGARATAGAGACAC | | | 4.8e-039 | 8.6e-037 | 1144 (4.6%) | | | | |
| 33-AGCACTAAATGCCCACAAGAGAAAGCGG | | | 4.8e-039 | 8.6e-037 | 1126 (4.5%) | | | | |
| 34-AATCAACAGAATATACATTYTTY | | | 4.8e-039 | 8.6e-037 | 1110 (4.4%) | | | | |
| 35-AACAAGGATAYCCAGGAATTGAA | | | 9.8e-039 | 1.8e-036 | 1135 (4.5%) | | | | |
| 36-CTGTAGATGTCTATTA | | | 9.8e-039 | 1.8e-036 | 1113 (4.5%) | | | | |
| 37-ATTTGCTGAGGAGWGCTTTACTTCCAA | | | 4.1e-038 | 7.4e-036 | 1134 (4.5%) | | | | |
| 38-CATCACAATTAAAAGAACTA | | | 8.5e-038 | 1.5e-035 | 1129 (4.5%) | | | | |
| 39-AATGACTACTGGGTACATAA | | | 8.5e-038 | 1.5e-035 | 1121 (4.5%) | | | | |
| 40-GTTTGTTATAATTTCTGTTCTTTTA | | | 8.5e-038 | 1.5e-035 | 1083 (4.3%) | | | | |
| 41-AAAAACCCTTCAAAAAA | | | 3.6e-037 | 6.4e-035 | 1046 (4.2%) | | | | |
| 42-CATAAAGCAAGTCCTKAGWGACCTAC | | | 7.3e-037 | 1.3e-034 | 1111 (4.4%) | | | | |
| 43-CCCTTTATCATTTTTTA | | | 1.3e-036 | 2.3e-034 | 1146 (4.6%) | | | | |
| 44-ATTAGTCTTGCTAGCGGTCTAT | | | 2.6e-036 | 4.7e-034 | 1069 (4.3%) | | | | |
| 45-TAGGGTGTCAATTTTRGATCTTT | | | 3.1e-036 | 5.5e-034 | 1063 (4.3%) | | | | |

**Stopped because 3 consecutive motifs exceeded the p-value threshold (0.05).**
**STREME ran for 62351.38 seconds.**

| Motif | Logo | RC Logo | P-value | E-value | Sites | More | Submit/Download | Positional Distribution | Matches per Sequence |
|---|---|---|---|---|---|---|---|---|---|
| 46-TTGTTGATCYTTTCAAAA | | | 6.3e-036 | 1.1e-033 | 1036 (4.1%) | | | | |
| 47-GGGTGCATATATATTTAGGACAG | | | 1.9e-033 | 3.5e-031 | 978 (3.9%) | | | | |
| 48-GAAACCAAYGAGAA | | | 2.8e-033 | 5.0e-031 | 978 (3.9%) | | | | |
| 49-AAGCAAGAGCAAACA | | | 5.6e-033 | 1.0e-030 | 1054 (4.2%) | | | | |
| 50-AGAAGAATCAAATAGAYRC | | | 1.4e-032 | 2.6e-030 | 1046 (4.2%) | | | | |
| 51-RCCATTCTCC | | | 1.5e-032 | 2.6e-030 | 1553 (6.2%) | | | | |
| 52-RACTTTCTGTCTC | | | 4.6e-031 | 8.2e-029 | 980 (3.9%) | | | | |
| 53-GGTTCAAGCRATT | | | 3.2e-030 | 5.7e-028 | 1562 (6.2%) | | | | |
| 54-CAGTGGCTCA | | | 7.7e-030 | 1.4e-027 | 1228 (4.9%) | | | | |
| 55-TTAGCTCTTCTTGTTGAA | | | 1.0e-029 | 1.8e-027 | 853 (3.4%) | | | | |
| 56-CTGATGGTAGTTTGTATTTC | | | 8.6e-029 | 1.5e-026 | 802 (3.2%) | | | | |
| 57-CAGCCTCCYR | | | 3.6e-026 | 6.4e-024 | 7466 (29.9%) | | | | |
| 58-AGWGGGGGGC | | | 1.7e-025 | 3.1e-023 | 6184 (24.7%) | | | | |
| 59-CTGGGACACATTYAAA | | | 6.2e-024 | 1.1e-021 | 845 (3.4%) | | | | |
| 60-GTCAACATTA | | | 3.4e-023 | 6.1e-021 | 840 (3.4%) | | | | |
| 61-AAAGAARAAAAGAG | | | 5.0e-022 | 8.9e-020 | 911 (3.6%) | | | | |
| 62-AGACAAAGAAGGCCA | | | 4.3e-021 | 7.7e-019 | 578 (2.3%) | | | | |
| 63-CCHCACACMC | | | 2.0e-020 | 3.6e-018 | 4472 (17.9%) | | | | |
| 64-AGAATACTAYAAACA | | | 1.8e-019 | 3.2e-017 | 615 (2.5%) | | | | |
| 65-AGAGCGAGAC | | | 4.9e-019 | 8.7e-017 | 1187 (4.7%) | | | | |
| 66-AAAAATTAGCCA | | | 5.6e-018 | 1.0e-015 | 1091 (4.4%) | | | | |
| 67-AAATAAACTAGAAAATC | | | 8.4e-017 | 1.5e-014 | 505 (2.0%) | | | | |
| 68-AGAGTGAGACTCT | | | 1.4e-016 | 2.4e-014 | 736 (2.9%) | | | | |
| 69-GGCTGGAGTG | | | 1.4e-015 | 2.6e-013 | 4031 (16.1%) | | | | |

**Stopped because 3 consecutive motifs exceeded the p-value threshold (0.05).**
**STREME ran for 62351.38 seconds.**

| Motif | Logo | RC Logo | P-value | E-value | Sites | More | Submit/Download | Positional Distribution | Matches per Sequence |
|---|---|---|---|---|---|---|---|---|---|
| 70-CAGRGCYCTG | | | 3.8e-015 | 6.8e-013 | 6242 (25.0%) | ↧ | ⇢ | | |
| 71-AACCAACAAAGATCAAAAG | | | 1.8e-014 | 3.3e-012 | 489 (2.0%) | ↧ | ⇢ | | |
| 72-ACCAAGCAGACC | | | 3.6e-014 | 6.5e-012 | 397 (1.6%) | ↧ | ⇢ | | |
| 73-GARCCYRGGARGTGGAGG | | | 5.5e-013 | 9.9e-011 | 587 (2.3%) | ↧ | ⇢ | | |
| 74-TGAGCCACCA | | | 2.1e-012 | 3.8e-010 | 488 (2.0%) | ↧ | ⇢ | | |
| 75-ATAAATTTCC | | | 2.6e-012 | 4.7e-010 | 992 (4.0%) | ↧ | ⇢ | | |
| 76-AGCCAGGYATGRTGGC | | | 6.5e-012 | 1.2e-009 | 773 (3.1%) | ↧ | ⇢ | | |
| 77-AGGGCCAGAG | | | 9.7e-012 | 1.7e-009 | 4344 (17.4%) | ↧ | ⇢ | | |
| 78-AGGGGGACAG | | | 1.1e-010 | 2.1e-008 | 3058 (12.2%) | ↧ | ⇢ | | |
| 79-ATTTATCCATTTCTTCTA | | | 1.2e-010 | 2.2e-008 | 379 (1.5%) | ↧ | ⇢ | | |
| 80-AACATGGTGAAA | | | 2.0e-010 | 3.5e-008 | 355 (1.4%) | ↧ | ⇢ | | |
| 81-GATAAAACAGACTTTA | | | 2.5e-010 | 4.4e-008 | 369 (1.5%) | ↧ | ⇢ | | |
| 82-SCTGSSCAGS | | | 2.5e-010 | 4.5e-008 | 6043 (24.2%) | ↧ | ⇢ | | |
| 83-AGAGACTAGGATTG | | | 4.2e-010 | 7.6e-008 | 253 (1.0%) | ↧ | ⇢ | | |
| 84-CCTCACTTCCCAG | | | 2.5e-009 | 4.4e-007 | 400 (1.6%) | ↧ | ⇢ | | |
| 85-GGCGGATCAK | | | 2.8e-009 | 5.0e-007 | 646 (2.6%) | ↧ | ⇢ | | |
| 86-AGTGGCTCAT | | | 3.3e-009 | 5.8e-007 | 322 (1.3%) | ↧ | ⇢ | | |
| 87-CCCCTCCTGA | | | 6.2e-009 | 1.1e-006 | 4335 (17.3%) | ↧ | ⇢ | | |
| 88-CCTCCHTCCY | | | 1.1e-008 | 2.1e-006 | 1487 (5.9%) | ↧ | ⇢ | | |
| 89-ACTCAGTGCTCAAT | | | 2.8e-008 | 5.0e-006 | 187 (0.7%) | ↧ | ⇢ | | |
| 90-ATGTGTCGAGGA | | | 3.2e-008 | 5.7e-006 | 280 (1.1%) | ↧ | ⇢ | | |
| 91-CCCCAACCCW | | | 4.7e-008 | 8.4e-006 | 1681 (6.7%) | ↧ | ⇢ | | |
| 92-AAAGAGACTTAGACTCCC | | | 5.6e-008 | 1.0e-005 | 229 (0.9%) | ↧ | ⇢ | | |
| 93-ATTGATCATTCTTGGG | | | 1.1e-007 | 2.0e-005 | 234 (0.9%) | ↧ | ⇢ | | |

**Stopped because 3 consecutive motifs exceeded the p-value threshold (0.05).**
**STREME ran for 62351.38 seconds.**

| Motif | Logo | RC Logo | P-value | E-value | Sites | More | Submit/Download | Positional Distribution | Matches per Sequence |
|---|---|---|---|---|---|---|---|---|---|
| 94-CATTTTGTTCTGTACTAAGAAGAA | | | 1.1e-007 | 2.0e-005 | 242 (1.0%) | | | | |
| 95-GCATGCTCGTTAAG | | | 1.1e-007 | 2.0e-005 | 227 (0.9%) | | | | |
| 96-ATTGTCCTATGACCCTG | | | 2.3e-007 | 4.1e-005 | 232 (0.9%) | | | | |
| 97-GTTTGTGTCCCTGGGTACTTGAGA | | | 2.3e-007 | 4.1e-005 | 228 (0.9%) | | | | |
| 98-CACAGCACATGTTTCAGA | | | 2.3e-007 | 4.1e-005 | 234 (0.9%) | | | | |
| 99-TCAGCACCACATYRCACT | | | 2.3e-007 | 4.1e-005 | 197 (0.8%) | | | | |
| 100-GTCTCAAAAAAA | | | 3.2e-007 | 5.7e-005 | 494 (2.0%) | | | | |
| 101-AAAGTGCTGGGATTA | | | 4.1e-007 | 7.3e-005 | 355 (1.4%) | | | | |
| 102-TAAAAATACAAAA | | | 4.4e-007 | 7.8e-005 | 969 (3.9%) | | | | |
| 103-GTGGGGAAAAGATTGAG | | | 9.2e-007 | 1.6e-004 | 206 (0.8%) | | | | |
| 104-CTAAACCAGGAAGA | | | 9.2e-007 | 1.6e-004 | 192 (0.8%) | | | | |
| 105-GCTTGAAGGCA | | | 9.2e-007 | 1.6e-004 | 213 (0.9%) | | | | |
| 106-MAGAGCGAGA | | | 1.2e-006 | 2.1e-004 | 669 (2.7%) | | | | |
| 107-ATAAACAAGTGAAC | | | 1.8e-006 | 3.3e-004 | 227 (0.9%) | | | | |
| 108-AATGAGCTGTTGGG | | | 2.7e-006 | 4.9e-004 | 224 (0.9%) | | | | |
| 109-WAAATAAATAAATAAATAA | | | 3.9e-006 | 7.0e-004 | 744 (3.0%) | | | | |
| 110-AGGAGAATCA | | | 6.7e-006 | 1.2e-003 | 307 (1.2%) | | | | |
| 111-GTGGATCACY | | | 7.0e-006 | 1.2e-003 | 410 (1.6%) | | | | |
| 112-CCAAGCAAATGGAAARCAAAAA | | | 7.4e-006 | 1.3e-003 | 187 (0.7%) | | | | |
| 113-TTAAATGGATTAA | | | 1.0e-005 | 1.8e-003 | 198 (0.8%) | | | | |
| 114-AGAGCAAGATCC | | | 1.0e-005 | 1.8e-003 | 246 (1.0%) | | | | |
| 115-AGCAATCCACC | | | 2.0e-005 | 3.5e-003 | 313 (1.3%) | | | | |
| 116-GGAAGTGAGGA | | | 2.9e-005 | 5.1e-003 | 382 (1.5%) | | | | |
| 117-AATACAGGAGCACCMAGAT | | | 3.0e-005 | 5.3e-003 | 166 (0.7%) | | | | |

**Stopped because 3 consecutive motifs exceeded the p-value threshold (0.05).**
**STREME ran for 62351.38 seconds.**

| Motif | Logo | RC Logo | P-value | E-value | Sites | More | Submit/Download | Positional Distribution | Matches per Sequence |
|---|---|---|---|---|---|---|---|---|---|
| 118-ATCGTCTGAGAT | | | 3.0e-005 | 5.3e-003 | 140 (0.6%) | | | | |
| 119-ATCTGTTTAACAAAGC | | | 3.2e-005 | 5.7e-003 | 232 (0.9%) | | | | |
| 120-CCTTCCCTTC | | | 5.4e-005 | 9.7e-003 | 1472 (5.9%) | | | | |
| 121-AGTTGAATCTCTGAA | | | 6.0e-005 | 1.1e-002 | 126 (0.5%) | | | | |
| 122-CATAGGCTCAAAATAAA | | | 6.0e-005 | 1.1e-002 | 119 (0.5%) | | | | |
| 123-CCCTTTCCCC | | | 6.0e-005 | 1.1e-002 | 2747 (11.0%) | | | | |
| 124-AGAGCAAAACTC | | | 7.6e-005 | 1.4e-002 | 217 (0.9%) | | | | |
| 125-AGCAATCTGCC | | | 7.6e-005 | 1.4e-002 | 221 (0.9%) | | | | |
| 126-TCACAGATCAACAGGATCCC | | | 1.2e-004 | 2.2e-002 | 154 (0.6%) | | | | |
| 127-AAAGAAAAGAAAAGAAA | | | 1.9e-004 | 3.4e-002 | 448 (1.8%) | | | | |
| 128-CTTTCTTTCTTTCTTT | | | 2.6e-004 | 4.7e-002 | 554 (2.2%) | | | | |
| 129-ACTGCAAGCTCCAC | | | 4.8e-004 | 8.6e-002 | 159 (0.6%) | | | | |
| 130-GATATCACCACYGATCCCA | | | 4.8e-004 | 8.6e-002 | 132 (0.5%) | | | | |
| 131-ATTTTATTTCTCCTTCACTTATGAAGC | | | 4.8e-004 | 8.6e-002 | 97 (0.4%) | | | | |
| 132-AGGCAGAAGAA | | | 5.7e-004 | 1.0e-001 | 233 (0.9%) | | | | |
| 133-ACTTTGGGAGGCC | | | 9.9e-004 | 1.8e-001 | 441 (1.8%) | | | | |
| 134-GATTACAGGCG | | | 1.3e-003 | 2.3e-001 | 197 (0.8%) | | | | |
| 135-CCCCTCCWWG | | | 1.4e-003 | 2.6e-001 | 2584 (10.3%) | | | | |
| 136-CCTGTTATTGGTCTA | | | 1.7e-003 | 3.0e-001 | 91 (0.4%) | | | | |
| 137-TGTCTGTGTAGAAAGAA | | | 1.9e-003 | 3.5e-001 | 104 (0.4%) | | | | |
| 138-ATGACAGGATCAAATTCATCCAT | | | 1.9e-003 | 3.5e-001 | 90 (0.4%) | | | | |
| 139-AAGAGCTCCTGAAGGAAGCACTA | | | 1.9e-003 | 3.5e-001 | 87 (0.3%) | | | | |
| 140-AACTGCATCAACTAAYGAGCAAAATAACCA | | | 1.9e-003 | 3.5e-001 | 81 (0.3%) | | | | |
| 141-AACACAGTGAAA | | | 1.9e-003 | 3.5e-001 | 96 (0.4%) | | | | |

**Stopped because 3 consecutive motifs exceeded the p-value threshold (0.05).**
**STREME ran for 62351.38 seconds.**

| Motif | Logo | RC Logo | P-value | E-value | Sites | More | Submit/Download | Positional Distribution | Matches per Sequence |
|---|---|---|---|---|---|---|---|---|---|
| 142-AGAGATCCTCC | | | 2.1e-003 | 3.7e-001 | 156 (0.6%) | ↓ | ⇢ | | |
| 143-AGAGGGAGAAG | | | 2.1e-003 | 3.8e-001 | 1194 (4.8%) | ↓ | ⇢ | | |
| 144-AGTGGCACTAT | | | 3.1e-003 | 5.6e-001 | 88 (0.4%) | ↓ | ⇢ | | |
| 145-ATATTCAACATTCTTAAAGAAA | | | 3.9e-003 | 7.0e-001 | 83 (0.3%) | ↓ | ⇢ | | |
| 146-AAGCAAATGCTGAGAGATTTTGT | | | 3.9e-003 | 7.0e-001 | 90 (0.4%) | ↓ | ⇢ | | |
| 147-AGCTGTGATGGC | | | 5.8e-003 | 1.0e+000 | 135 (0.5%) | ↓ | ⇢ | | |
| 148-TACACCTCCCA | | | 5.8e-003 | 1.0e+000 | 93 (0.4%) | ↓ | ⇢ | | |
| 149-AGCTATCCTCC | | | 5.8e-003 | 1.0e+000 | 147 (0.6%) | ↓ | ⇢ | | |
| 150-ATGGTGAAACCCYR | | | 6.3e-003 | 1.1e+000 | 219 (0.9%) | ↓ | ⇢ | | |
| 151-GTCAATATTAGACAGATCAA | | | 7.8e-003 | 1.4e+000 | 77 (0.3%) | ↓ | ⇢ | | |
| 152-AAGCCCATCAGACTAA | | | 7.8e-003 | 1.4e+000 | 75 (0.3%) | ↓ | ⇢ | | |
| 153-AGTTTGGCTGGATATGAAATTCTGGGTT | | | 7.8e-003 | 1.4e+000 | 91 (0.4%) | ↓ | ⇢ | | |
| 154-AAAAAAAAAAAATT | | | 9.5e-003 | 1.7e+000 | 196 (0.8%) | ↓ | ⇢ | | |
| 155-TCACCCCTTACCA | | | 1.1e-002 | 1.9e+000 | 133 (0.5%) | ↓ | ⇢ | | |
| 156-ACTGCACCACT | | | 1.1e-002 | 2.0e+000 | 120 (0.5%) | ↓ | ⇢ | | |
| 157-ACAAATGTTTCTTCTAACAWCCCCACAA | | | 1.6e-002 | 2.8e+000 | 87 (0.3%) | ↓ | ⇢ | | |
| 158-ACACATAATYGTCAGATTCACCAAR | | | 1.6e-002 | 2.8e+000 | 86 (0.3%) | ↓ | ⇢ | | |
| 159-CCACTCTTAACTC | | | 1.6e-002 | 2.8e+000 | 86 (0.3%) | ↓ | ⇢ | | |
| 160-AAGGTCAAGTTGTCTGGAC | | | 1.6e-002 | 2.8e+000 | 76 (0.3%) | ↓ | ⇢ | | |
| 161-ACCCTGTAGCATT | | | 1.6e-002 | 2.8e+000 | 74 (0.3%) | ↓ | ⇢ | | |
| 162-GTTGAAATGAAGGAAAAAATGTTA | | | 1.6e-002 | 2.8e+000 | 70 (0.3%) | ↓ | ⇢ | | |
| 163-AAGAAAATAGATTTTGGAAGTTATGA | | | 1.6e-002 | 2.8e+000 | 63 (0.3%) | ↓ | ⇢ | | |
| 164-GGACTGGACA | | | 1.9e-002 | 3.5e+000 | 64 (0.3%) | ↓ | ⇢ | | |
| 165-CTGGTCTCAAA | | | 2.6e-002 | 4.7e+000 | 232 (0.9%) | ↓ | ⇢ | | |

**Stopped because 3 consecutive motifs exceeded the p-value threshold (0.05).**
**STREME ran for 62351.38 seconds.**

| Motif | Logo / RC Logo | P-value | E-value | Sites | More | Submit/Download | Positional Distribution / Matches per Sequence |
|---|---|---|---|---|---|---|---|
| 166-AGAAGGAGAAAAACAGGTATWAAAGGW | 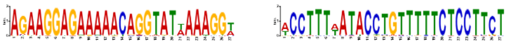 | 3.1e-002 | 5.6e+000 | 94 (0.4%) | ↧ | ⇢ | 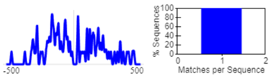 |
| 167-AAACAGTGTAAACCGGCAGTGT | 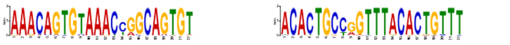 | 3.1e-002 | 5.6e+000 | 81 (0.3%) | ↧ | ⇢ | 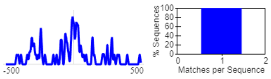 |
| 168-ATGAGTAGTTGAGAAYGGTGAATA | 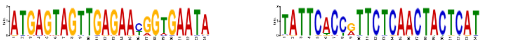 | 3.1e-002 | 5.6e+000 | 81 (0.3%) | ↧ | ⇢ | 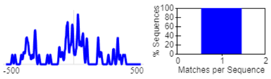 |
| 169-CTTGTCATCCCTA | 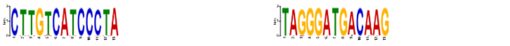 | 3.1e-002 | 5.6e+000 | 75 (0.3%) | ↧ | ⇢ | 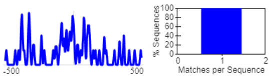 |
| 170-AGCTCCTGTATAGAYGCTCCTTTT | 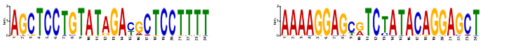 | 3.1e-002 | 5.6e+000 | 68 (0.3%) | ↧ | ⇢ | 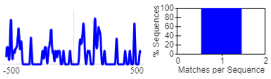 |
| 171-ATTAGAGAGTGCC | 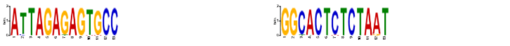 | 3.1e-002 | 5.6e+000 | 68 (0.3%) | ↧ | ⇢ | 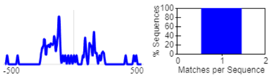 |
| 172-GATCTCAAACATGCTTTCTTTACTAT | 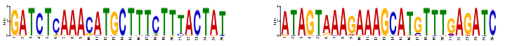 | 3.1e-002 | 5.6e+000 | 60 (0.2%) | ↧ | ⇢ | 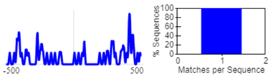 |
| 173-AGAGATTAAGCT | 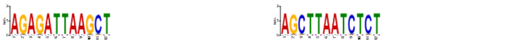 | 3.1e-002 | 5.6e+000 | 68 (0.3%) | ↧ | ⇢ | 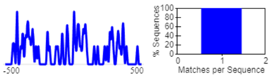 |
| 174-CTACAAGCCAGA | 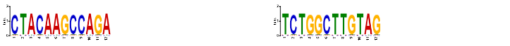 | 3.1e-002 | 5.6e+000 | 63 (0.3%) | ↧ | ⇢ | 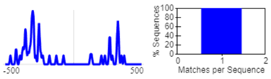 |
| 175-CATTCCATTCCATTC | 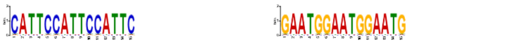 | 3.5e-002 | 6.3e+000 | 58 (0.2%) | ↧ | ⇢ | 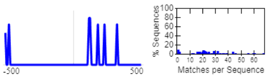 |
| 176-ATGGGTCTTGA | 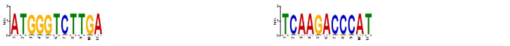 | 3.5e-002 | 6.3e+000 | 64 (0.3%) | ↧ | ⇢ | 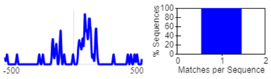 |
| 177-AAAAAATTTAAAAATT | 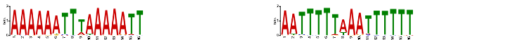 | 5.3e-002 | 9.6e+000 | 232 (0.9%) | ↧ | ⇢ | 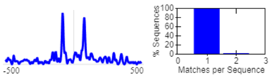 |
| 178-ACCTGWCCTAAAACC | 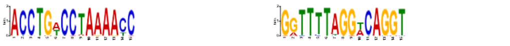 | 6.2e-002 | 1.1e+001 | 120 (0.5%) | ↧ | ⇢ | 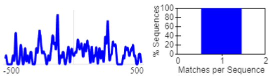 |
| 179-TTAGTTTYTCAATTCATACAAAA | 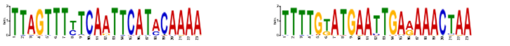 | 6.2e-002 | 1.1e+001 | 87 (0.3%) | ↧ | ⇢ | 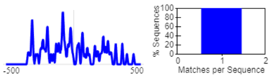 |

**Stopped because 3 consecutive motifs exceeded the p-value threshold (0.05).**
**STREME ran for 62351.38 seconds.**

#### Inputs & Settings

Previous Next Top

##### Sequences

| Role | Source | Alphabet | Sequence Count | Total Size |
|---|---|---|---|---|
| Positive (primary) Sequences | /tmp/Rtmp38v5rK/file6a94263b50df6.fa | DNA | 25000 | 25000000 |
| Negative (control) Sequences | /tmp/Rtmp38v5rK/file6a9425357e970.fa | DNA | 25000 | 25000000 |

##### Background Model

**Source:** built from the negative (control) sequences

**Order:** 2 (only order-0 shown)

| Name | Freq. | Bg. | | | | Bg. | Freq. | Name |
|---|---|---|---|---|---|---|---|---|
| Adenine | 0.254 | 0.337 | A | ~ | T | 0.337 | 0.254 | Thymine |
| Cytosine | 0.246 | 0.163 | C | ~ | G | 0.163 | 0.246 | Guanine |

##### Other Settings

| | |
|---|---|
| **Strand Handling** | Both the given and reverse complement strands are processed. |
| **Objective Function** | Differential Enrichment |
| **Statistical Test** | Fisher Exact Test |
| **Minimum Motif Width** | 10 |
| **Maximum Motif Width** | 30 |
| **Test Set** | 10% of the input sequences were randomly assigned to the test set. |
| **Word Evaluation** | Up to 25 words of each width from 10 to 30 were evaluated to find seeds. |
| **Seed Refinement** | Up to 4 seeds of each width from 10 to 30 were further refined. |
| **Refinement Iterations** | Up to 20 iterations were allowed when refining a seed. |
| **Random Number Seed** | 0 |
| **Trimming of Control Sequences** | Trimming of control sequences was allowed. |
| **Total Length** | The total length of the sequence set(s) was not limited. |
| **Maximum Motif p-value** | Stop when the p-value is greater than 0.05 for 3 consecutive motifs. |
| **Maximum Motifs to Find** | No maximum number of motifs. |
| **Maximum Run Time** | No maximum running time. |

Previous Top

**STREME version**
5.5.5 (Release date: Thu Sep 14 08:48:04 2023 +1000)

**Reference**
Timothy L. Bailey, "STREME: accurate and versatile sequence motif discovery", *Bioinformatics*, Mar. 24, 2021. [full text]

**Command line**
```
streme --p /tmp/Rtmp38v5rK/file6a94263b50df6.fa --n /tmp/Rtmp38v5rK/file6a9425357e970.fa --oc /g/strcombio/fsupek_home/mveiner/Projects/ORI/out/Motifs/top/streme --minw 10 --maxw 30 --dna
```

**Tomtom**
Motif Comparison Tool

For further information on how to interpret these results please access http://meme-suite.org/doc/tomtom-output-format.html.
To get a copy of the MEME software please access http://meme-suite.org.

If you use TOMTOM in your research, please cite the following paper:
Shobhit Gupta, JA Stamatoyannopolous, Timothy Bailey and William Stafford Noble, "Quantifying similarity between motifs", *Genome Biology*, **8**(2):R24, 2007. [full text]



#### Query Motifs

Next Top

| Database ? | ID ? | Alt. ID ? | Preview ? | Matches ? | List ? |
|---|---|---|---|---|---|
| motif_01 | 1-GGGAGGCYGAGGCR | STREME-1 | 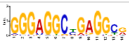 | 33 | ZN770_HUMAN.H11MO.0.C, ZN281_HUMAN.H11MO.0.A, SP2_HUMAN.H11MO.0.A, PATZ1_HUMAN.H11MO.0.C, SP3_HUMAN.H11MO.0.B, VEZF1_HUMAN.H11MO.0.C, WT1_HUMAN.H11MO.0.C, ZBT17_HUMAN.H11MO.0.A, TAF1_HUMAN.H11MO.0.A, MAZ_HUMAN.H11MO.0.A, ZN263_HUMAN.H11MO.0.A, ZN467_HUMAN.H11MO.0.C, KLF15_HUMAN.H11MO.0.A, SP4_HUMAN.H11MO.0.A, |

#### Target Databases

Previous Next Top

| Database ? | Used ? | Matched ? |
|---|---|---|
| HOCOMOCOv11_core_HUMAN_mono_meme_format | 401 | 33 |

#### Matches to 1-GGGAGGCYGAGGCR (STREME-1)

Previous Next Top

**Summary ?**

| | |
|---|---|
| **Name** | ZN770_HUMAN.H11MO.0.C |
| **Database** | HOCOMOCOv11_core_HUMAN_mono_meme_format |
| ***p*-value** | 1.23e-10 |
| ***E*-value** | 4.92e-08 |
| ***q*-value** | 9.64e-08 |
| **Overlap** | 14 |
| **Offset** | 0 |
| **Orientation** | Normal |

Show logo download options

**Optimal Alignment ?**

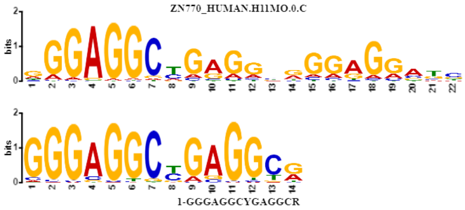


**Summary ?**

| | |
|---|---|
| **Name** | ZN281_HUMAN.H11MO.0.A |
| **Database** | HOCOMOCOv11_core_HUMAN_mono_meme_format |
| ***p*-value** | 7.43e-04 |
| ***E*-value** | 2.98e-01 |
| ***q*-value** | 2.24e-01 |
| **Overlap** | 14 |
| **Offset** | 0 |
| **Orientation** | Normal |

Show logo download options

**Optimal Alignment ?**

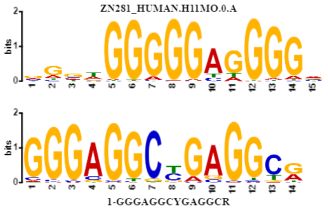


**Summary ?**

| | |
|---|---|
| **Name** | SP2_HUMAN.H11MO.0.A |
| **Database** | HOCOMOCOv11_core_HUMAN_mono_meme_format |
| ***p*-value** | 1.41e-03 |
| ***E*-value** | 5.66e-01 |
| ***q*-value** | 2.24e-01 |
| **Overlap** | 14 |
| **Offset** | 4 |
| **Orientation** | Normal |

Show logo download options

**Optimal Alignment ?**

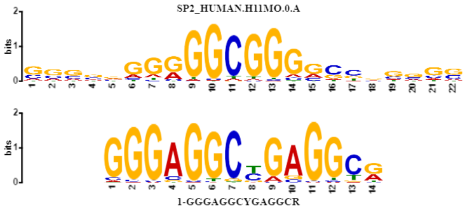


**Summary ?**

| | |
|---|---|
| **Name** | PATZ1_HUMAN.H11MO.0.C |
| **Database** | HOCOMOCOv11_core_HUMAN_mono_meme_format |
| ***p*-value** | 1.70e-03 |
| ***E*-value** | 6.81e-01 |
| ***q*-value** | 2.24e-01 |
| **Overlap** | 14 |
| **Offset** | 1 |
| **Orientation** | Normal |

Show logo download options

**Optimal Alignment ?**

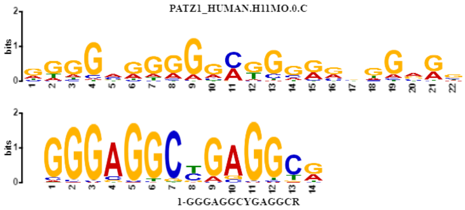


**Summary ?**

| | |
|---|---|
| **Name** | SP3_HUMAN.H11MO.0.B |
| **Database** | HOCOMOCOv11_core_HUMAN_mono_meme_format |
| ***p*-value** | 1.84e-03 |
| ***E*-value** | 7.37e-01 |
| ***q*-value** | 2.24e-01 |
| **Overlap** | 14 |
| **Offset** | 3 |
| **Orientation** | Normal |

Show logo download options

**Optimal Alignment ?**

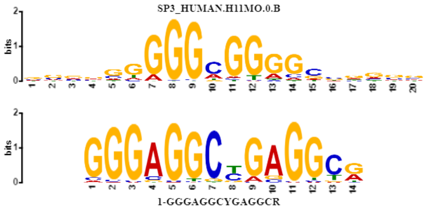


**Summary ?**

| | |
|---|---|
| **Name** | VEZF1_HUMAN.H11MO.0.C |
| **Database** | HOCOMOCOv11_core_HUMAN_mono_meme_format |
| ***p*-value** | 2.04e-03 |
| ***E*-value** | 8.16e-01 |
| ***q*-value** | 2.24e-01 |
| **Overlap** | 14 |
| **Offset** | 8 |
| **Orientation** | Normal |

Show logo download options

**Optimal Alignment ?**

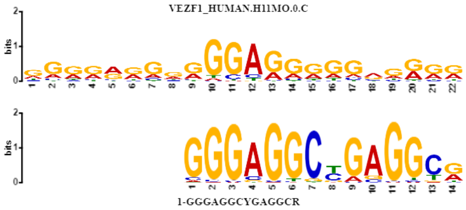


**Summary ?**

| | |
|---|---|
| **Name** | WT1_HUMAN.H11MO.0.C |
| **Database** | HOCOMOCOv11_core_HUMAN_mono_meme_format |
| ***p*-value** | 2.47e-03 |
| ***E*-value** | 9.89e-01 |
| ***q*-value** | 2.24e-01 |
| **Overlap** | 14 |
| **Offset** | 0 |
| **Orientation** | Normal |

Show logo download options

**Optimal Alignment ?**

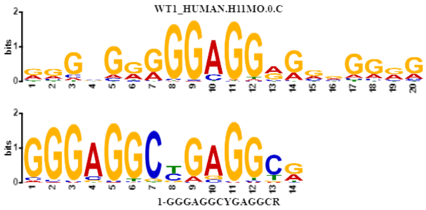


**Summary ?**

**Optimal Alignment ?**

| | |
|---|---|
| **Name** | ZBT17_HUMAN.H11MO.0.A |
| **Database** | HOCOMOCOv11_core_HUMAN_mono_meme_format |
| ***p*-value** | 2.57e-03 |
| ***E*-value** | 1.03e+00 |
| ***q*-value** | 2.24e-01 |
| **Overlap** | 14 |
| **Offset** | 2 |
| **Orientation** | Normal |

Show logo download options

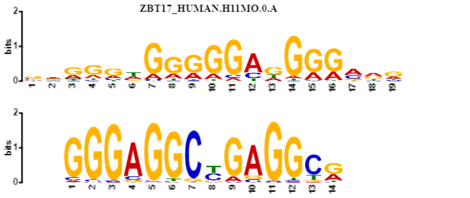


**Summary** **Optimal Alignment**

| | |
|---|---|
| **Name** | TAF1_HUMAN.H11MO.0.A |
| **Database** | HOCOMOCOv11_core_HUMAN_mono_meme_format |
| ***p*-value** | 2.83e-03 |
| ***E*-value** | 1.13e+00 |
| ***q*-value** | 2.24e-01 |
| **Overlap** | 14 |
| **Offset** | 1 |
| **Orientation** | Normal |

Show logo download options

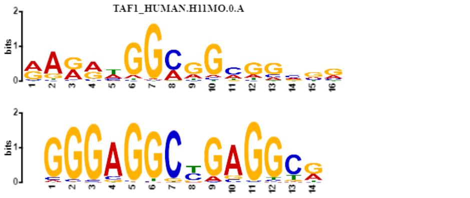


**Summary** **Optimal Alignment**

| | |
|---|---|
| **Name** | MAZ_HUMAN.H11MO.0.A |
| **Database** | HOCOMOCOv11_core_HUMAN_mono_meme_format |
| ***p*-value** | 3.26e-03 |
| ***E*-value** | 1.31e+00 |
| ***q*-value** | 2.24e-01 |
| **Overlap** | 14 |
| **Offset** | 2 |
| **Orientation** | Normal |

Show logo download options

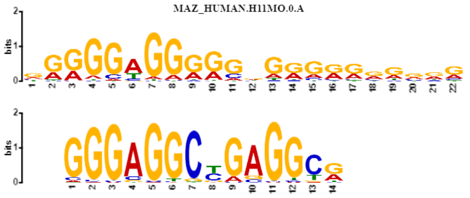


**Summary** **Optimal Alignment**

| | |
|---|---|
| **Name** | ZN263_HUMAN.H11MO.0.A |
| **Database** | HOCOMOCOv11_core_HUMAN_mono_meme_format |
| ***p*-value** | 3.69e-03 |
| ***E*-value** | 1.48e+00 |
| ***q*-value** | 2.24e-01 |
| **Overlap** | 14 |
| **Offset** | 0 |
| **Orientation** | Normal |

Show logo download options

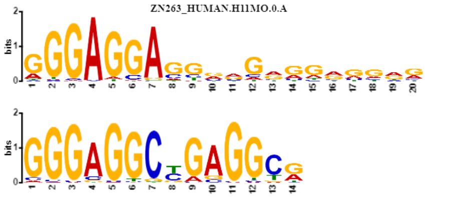


**Summary** **Optimal Alignment**

| | |
|---|---|
| **Name** | ZN467_HUMAN.H11MO.0.C |
| **Database** | HOCOMOCOv11_core_HUMAN_mono_meme_format |
| ***p*-value** | 3.87e-03 |
| ***E*-value** | 1.55e+00 |
| ***q*-value** | 2.24e-01 |
| **Overlap** | 14 |
| **Offset** | 1 |
| **Orientation** | Normal |

Show logo download options

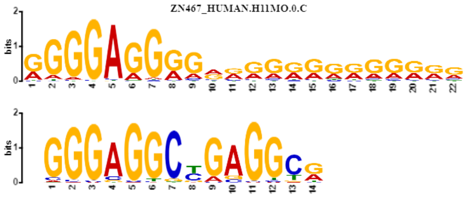


**Summary** **Optimal Alignment**

| | |
|---|---|
| **Name** | KLF15_HUMAN.H11MO.0.A |
| **Database** | HOCOMOCOv11_core_HUMAN_mono_meme_format |
| ***p*-value** | 4.06e-03 |
| ***E*-value** | 1.63e+00 |
| ***q*-value** | 2.24e-01 |
| **Overlap** | 14 |
| **Offset** | 1 |
| **Orientation** | Normal |

Show logo download options

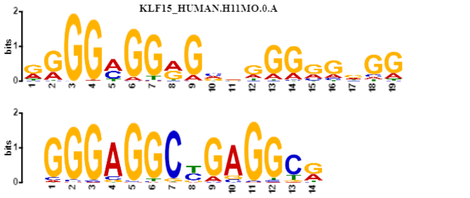


**Summary** **Optimal Alignment**

| | |
|---|---|
| **Name** | SP4_HUMAN.H11MO.0.A |
| **Database** | HOCOMOCOv11_core_HUMAN_mono_meme_format |
| ***p*-value** | 4.13e-03 |
| ***E*-value** | 1.65e+00 |
| ***q*-value** | 2.24e-01 |
| **Overlap** | 14 |
| **Offset** | 4 |
| **Orientation** | Normal |

Show logo download options

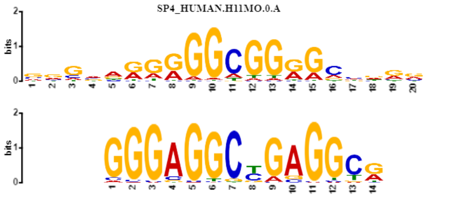


**Summary** **Optimal Alignment**

| | |
|---|---|
| **Name** | KLF4_HUMAN.H11MO.0.A |
| **Database** | HOCOMOCOv11_core_HUMAN_mono_meme_format |
| ***p*-value** | 4.37e-03 |
| ***E*-value** | 1.75e+00 |
| ***q*-value** | 2.24e-01 |
| **Overlap** | 10 |
| **Offset** | -3 |
| **Orientation** | Normal |

Show logo download options

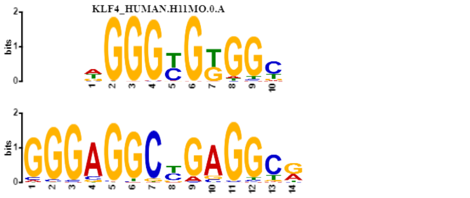


**Summary** **Optimal Alignment**

| | |
|---|---|
| **Name** | SRBP2_HUMAN.H11MO.0.B |
| **Database** | HOCOMOCOv11_core_HUMAN_mono_meme_format |
| ***p*-value** | 4.74e-03 |
| ***E*-value** | 1.90e+00 |
| ***q*-value** | 2.24e-01 |
| **Overlap** | 12 |
| **Offset** | 1 |
| **Orientation** | Normal |

Show logo download options

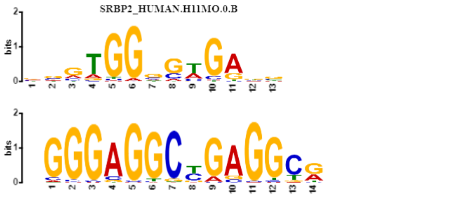


**Summary** **Optimal Alignment**

**Name** SP1_HUMAN.H11MO.0.A
**Database** HOCOMOCOv11_core_HUMAN_mono_meme_format
***p*-value** 4.85e-03
***E*-value** 1.94e+00
***q*-value** 2.24e-01
**Overlap** 14
**Offset** 5
**Orientation** Normal
Show logo download options

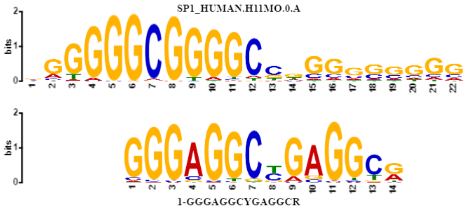


**Summary** **Optimal Alignment**

**Name** ELF5_HUMAN.H11MO.0.A
**Database** HOCOMOCOv11_core_HUMAN_mono_meme_format
***p*-value** 6.19e-03
***E*-value** 2.48e+00
***q*-value** 2.70e-01
**Overlap** 14
**Offset** 0
**Orientation** Normal
Show logo download options

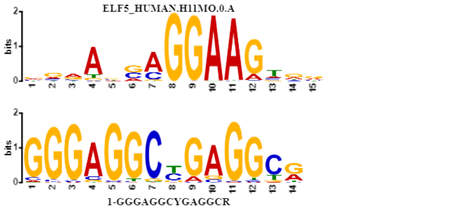


**Summary** **Optimal Alignment**

**Name** KLF3_HUMAN.H11MO.0.B
**Database** HOCOMOCOv11_core_HUMAN_mono_meme_format
***p*-value** 6.66e-03
***E*-value** 2.67e+00
***q*-value** 2.76e-01
**Overlap** 14
**Offset** 2
**Orientation** Normal
Show logo download options

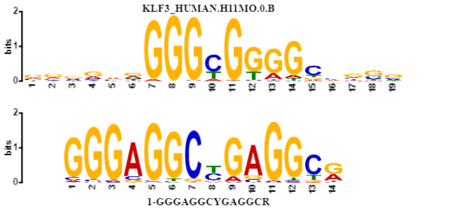


**Summary** **Optimal Alignment**

**Name** KLF6_HUMAN.H11MO.0.A
**Database** HOCOMOCOv11_core_HUMAN_mono_meme_format
***p*-value** 8.26e-03
***E*-value** 3.31e+00
***q*-value** 3.24e-01
**Overlap** 14
**Offset** 0
**Orientation** Normal
Show logo download options

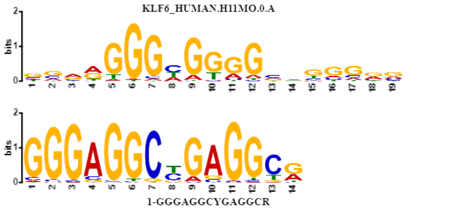


**Summary** **Optimal Alignment**

**Name** GABPA_HUMAN.H11MO.0.A
**Database** HOCOMOCOv11_core_HUMAN_mono_meme_format
***p*-value** 8.86e-03
***E*-value** 3.55e+00
***q*-value** 3.31e-01
**Overlap** 13
**Offset** -1
**Orientation** Normal
Show logo download options

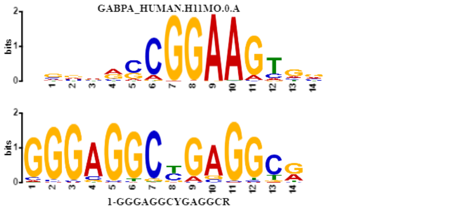


**Summary** **Optimal Alignment**

**Name** EGR2_HUMAN.H11MO.0.A
**Database** HOCOMOCOv11_core_HUMAN_mono_meme_format
***p*-value** 9.80e-03
***E*-value** 3.93e+00
***q*-value** 3.50e-01
**Overlap** 14
**Offset** 1
**Orientation** Normal
Show logo download options

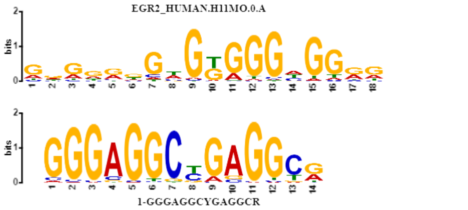


**Summary** **Optimal Alignment**

**Name** SALL4_HUMAN.H11MO.0.B
**Database** HOCOMOCOv11_core_HUMAN_mono_meme_format
***p*-value** 1.06e-02
***E*-value** 4.23e+00
***q*-value** 3.59e-01
**Overlap** 10
**Offset** -1
**Orientation** Normal
Show logo download options

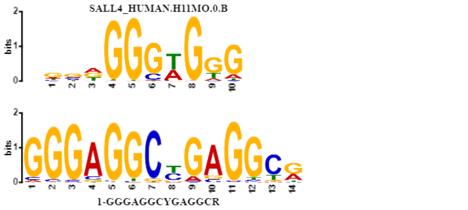


**Summary** **Optimal Alignment**

**Name** RXRA_HUMAN.H11MO.0.A
**Database** HOCOMOCOv11_core_HUMAN_mono_meme_format
***p*-value** 1.10e-02
***E*-value** 4.40e+00
***q*-value** 3.59e-01
**Overlap** 14
**Offset** 2
**Orientation** Normal
Show logo download options

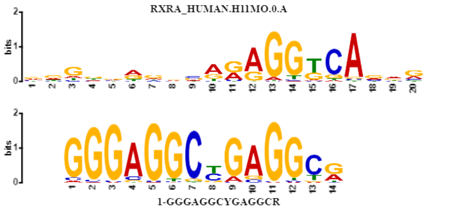


**Summary** **Optimal Alignment**

**Name** E2F6_HUMAN.H11MO.0.A
**Database** HOCOMOCOv11_core_HUMAN_mono_meme_format
***p*-value** 1.18e-02
***E*-value** 4.74e+00
***q*-value** 3.71e-01
**Overlap** 12
**Offset** 1
**Orientation** Normal
Show logo download options

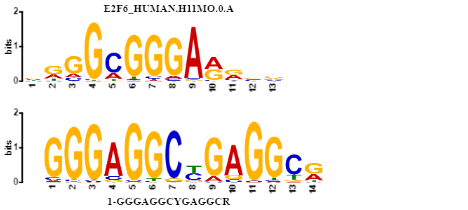


**Summary** **Optimal Alignment**

**Name** [EGR1_HUMAN.H11MO.0.A]
**Database** HOCOMOCOv11_core_HUMAN_mono_meme_format

***p*-value** 1.31e-02
***E*-value** 5.26e+00
***q*-value** 3.97e-01

**Overlap** 13
**Offset** -1
**Orientation** Normal
Show logo download options

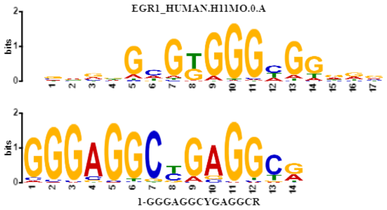


**Summary** ? **Optimal Alignment** ?

**Name** [KLF9_HUMAN.H11MO.0.C]
**Database** HOCOMOCOv11_core_HUMAN_mono_meme_format

***p*-value** 1.40e-02
***E*-value** 5.63e+00
***q*-value** 4.06e-01

**Overlap** 14
**Offset** 1
**Orientation** Normal
Show logo download options

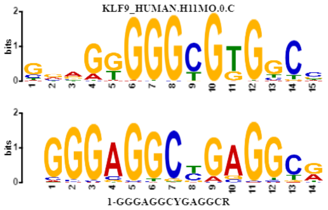


**Summary** ? **Optimal Alignment** ?

**Name** [CRX_HUMAN.H11MO.0.B]
**Database** HOCOMOCOv11_core_HUMAN_mono_meme_format

***p*-value** 1.45e-02
***E*-value** 5.80e+00
***q*-value** 4.06e-01

**Overlap** 12
**Offset** 1
**Orientation** Normal
Show logo download options

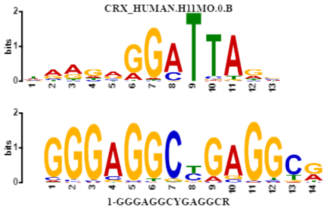


**Summary** ? **Optimal Alignment** ?

**Name** [ZBTB6_HUMAN.H11MO.0.C]
**Database** HOCOMOCOv11_core_HUMAN_mono_meme_format

***p*-value** 1.94e-02
***E*-value** 7.80e+00
***q*-value** 5.12e-01

**Overlap** 13
**Offset** -1
**Orientation** Normal
Show logo download options

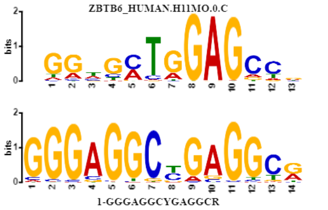


**Summary** ? **Optimal Alignment** ?

**Name** [P63_HUMAN.H11MO.0.A]
**Database** HOCOMOCOv11_core_HUMAN_mono_meme_format

***p*-value** 1.96e-02
***E*-value** 7.86e+00
***q*-value** 5.12e-01

**Overlap** 14
**Offset** 0
**Orientation** Normal
Show logo download options

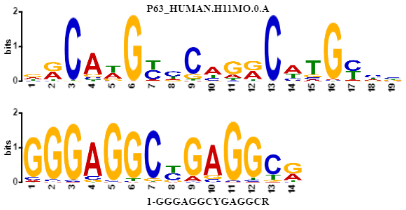


**Summary** ? **Optimal Alignment** ?

**Name** [ETS2_HUMAN.H11MO.0.B]
**Database** HOCOMOCOv11_core_HUMAN_mono_meme_format

***p*-value** 2.04e-02
***E*-value** 8.19e+00
***q*-value** 5.12e-01

**Overlap** 13
**Offset** 0
**Orientation** Normal
Show logo download options

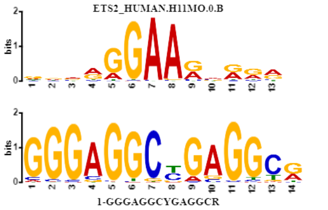


**Summary** ? **Optimal Alignment** ?

**Name** [ZFX_HUMAN.H11MO.0.A]
**Database** HOCOMOCOv11_core_HUMAN_mono_meme_format

***p*-value** 2.09e-02
***E*-value** 8.37e+00
***q*-value** 5.12e-01

**Overlap** 10
**Offset** -1
**Orientation** Normal
Show logo download options

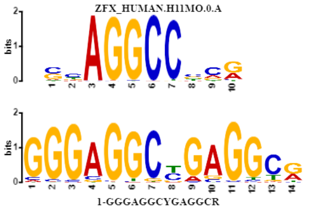


**Summary** ? **Optimal Alignment** ?

**Name** [MYC_HUMAN.H11MO.0.A]
**Database** HOCOMOCOv11_core_HUMAN_mono_meme_format

***p*-value** 2.45e-02
***E*-value** 9.83e+00
***q*-value** 5.84e-01

**Overlap** 11
**Offset** -2
**Orientation** Reverse Complement
Show logo download options

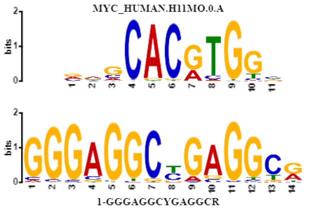


#### SETTINGS

Previous Next Top

**Alphabet**

**Source:** the query file

| Name ? | Bg. ? | | | | Bg. ? | Name ? |
|---|---|---|---|---|---|---|
| Adenine | 0.25 | A | ~ | T | 0.25 | Thymine |
| Cytosine | 0.25 | C | ~ | G | 0.25 | Guanine |

**Other Settings**

| | |
|---|---|
| **Strand Handling** | Motifs may be reverse complemented before comparison to find a better match. |
| **Distance Measure** | Pearson correlation coefficient |
| **Match Threshold** | Matches must have a *E*-value of 10 or smaller. |

Previous Top

**TOMTOM version**
5.0.5 (Release date: Mon Mar 18 20:12:19 2019 -0700)

**Reference**
Shobhit Gupta, JA Stamatoyannopolous, Timothy Bailey and William Stafford Noble, "Quantifying similarity between motifs", *Genome Biology*, **8**(2):R24, 2007.

**Command line**
```
tomtom -no-ssc -oc /g/strcombio/fsupek_home/mveiner/Projects/ORI/out/Motifs/top/tomtom/motif_01_results -verbosity 1 -min-overlap 5 -mi 1 -dist pearson -evalue -thresh 10.0 /g/strcombio/fsupek_home/mveiner/Projects/ORI/out/Motifs/top/tomtom/motif_01
```

Result calculation took 0.857 seconds

### CentriMo
Local Motif Enrichment Analysis

For further information on how to interpret these results please access https://meme-suite.org/meme/doc/centrimo-output-format.html.
To get a copy of the MEME software please access https://meme-suite.org.

If you use CentriMo in your research, please cite the following paper:
Timothy L. Bailey and Philip Machanick, "Inferring direct DNA binding from ChIP-seq", *Nucleic Acids Research*, **40**:e128, 2012. [Full Text] If you use CentriMo in your research, please cite the following paper:
Timothy L. Bailey and Philip Machanick, "Inferring direct DNA binding from ChIP-seq", *Nucleic Acids Research*, **40**:e128, 2012. [Full Text]



#### Results

##### Motif Probability Graph (motif score ≥ optimal Score Threshold for given motif ≥ 0) ?

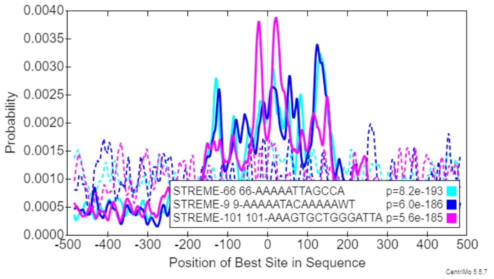


##### Options

**Plotting** ?

66-AAAAATTAGCCA
9-AAAAATACAAAAAWT
101-AAAGTGCTGGGATTA

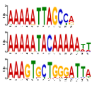

**Unused Colors** ?

**Graph** ?
Smoothing: Weighted Moving Average
Window: 20
Legend: Enabled (click on graph to move)
X-axis: Position of Best Site in Sequence
Negative sequences: Plotted as dashed line
Zoom: Undo Zoom | Center on 0

Download EPS (for publication)

##### Enriched motifs (*E*-value ≤ 10 using the binomial test )

| ☒ | ID ? | Alt ID ? | Consensus ? | Concentration ? | *E*-value ? | Fisher *E*-value ? | Region Width ? | Region Matches ? | Negative Region Matches ? | Score Threshold ? |
|---|---|---|---|---|---|---|---|---|---|---|
| ■ | 66-AAAAATTAGCCA | STREME-66 | AAAAATTAGCCA | 0.0422 | 1.5e-190 | 1.9e-56 | 339 | 1566 | 419 | 4.61e+0 |
| ☐ | 134-GATTACAGGCG | STREME-134 | GATTACAGGCG | 0.0443 | 2.1e-152 | 3.3e-51 | 332 | 1606 | 341 | 7.98e+0 |
| ☐ | 76-AGCCAGGYATGRTGGC | STREME-76 | AGCCAGGYATGRTGGC | 0.0652 | 4.9e-173 | 2.4e-50 | 365 | 1640 | 275 | 1.27e+1 |
| ☐ | 16-GGCAGATCACC | STREME-16 | GGCAGATCACC | 0.0415 | 6.8e-138 | 8.3e-44 | 268 | 1544 | 510 | 4.51e+0 |
| ☐ | 2-GCTGGGAYTACAGGCG | STREME-2 | GCTGGGAYTACAGGCG | 0.0485 | 6.9e-169 | 2.0e-42 | 337 | 1590 | 269 | 1.45e+1 |
| ☐ | 80-AACATGGTGAAA | STREME-80 | AACATGGTGAAA | 0.0205 | 9.0e-170 | 2.8e-42 | 379 | 1378 | 217 | 5.59e+0 |
| ☐ | 150-ATGGTGAAACCCYR | STREME-150 | ATGGTGAAACCCYR | 0.0292 | 8.2e-173 | 2.9e-41 | 359 | 1397 | 234 | 9.89e+0 |
| ☐ | 1-GGGAGGCYGAGGCR | STREME-1 | GGGAGGCYGAGGCR | 0.0784 | 1.3e-152 | 5.3e-40 | 431 | 1767 | 306 | 1.59e+1 |
| ■ | 9-AAAAATACAAAAAWT | STREME-9 | AAAAATACAAAAAWT | 0.0480 | 1.1e-183 | 7.7e-40 | 326 | 1339 | 241 | 1.34e+1 |
| ■ | 101-AAAGTGCTGGGATTA | STREME-101 | AAAGTGCTGGGATTA | 0.0364 | 1.0e-182 | 2.3e-39 | 316 | 1425 | 254 | 1.10e+1 |
| ☐ | 102-TAAAAATACAAAA | STREME-102 | TAAAAATACAAAA | 0.0534 | 9.5e-156 | 5.3e-38 | 330 | 1232 | 209 | 1.46e+1 |
| ☐ | 133-ACTTTGGGAGGCC | STREME-133 | ACTTTGGGAGGCC | 0.0627 | 1.9e-172 | 2.7e-37 | 298 | 1371 | 230 | 1.31e+1 |
| ☐ | 5-CCCCGTCTCTAC | STREME-5 | CCCCGTCTCTAC | 0.0453 | 7.9e-182 | 6.5e-37 | 339 | 1440 | 218 | 1.38e+1 |
| ☐ | 4-CTCCTGACCTCR | STREME-4 | CTCCTGACCTCR | 0.0209 | 2.5e-176 | 4.0e-36 | 249 | 1287 | 191 | 1.32e+1 |
| ☐ | 12-CCRTGTTGGCC | STREME-12 | CCRTGTTGGCC | 0.0258 | 5.2e-139 | 9.1e-36 | 246 | 1504 | 328 | 1.01e+1 |
| ☐ | 11-CCACCVCVCC | STREME-11 | CCACCVCVCC | 0.0641 | 1.5e-117 | 2.0e-33 | 361 | 1635 | 334 | 1.23e+1 |
| ☐ | 8-CVAGACCAKCCY | STREME-8 | CVAGACCAKCCY | 0.0211 | 3.4e-152 | 3.2e-28 | 227 | 1212 | 182 | 1.23e+1 |
| ☐ | 57-CAGCCTCCYR | STREME-57 | CAGCCTCCYR | 0.0862 | 1.3e-136 | 5.1e-28 | 439 | 1440 | 240 | 1.42e+1 |
| ☐ | 53-GGTTCAAGCRATT | STREME-53 | GGTTCAAGCRATT | 0.0290 | 1.8e-115 | 1.5e-27 | 458 | 1735 | 474 | 8.03e+0 |
| ☐ | 17-RGGCWGAGGS | STREME-17 | RGGCWGAGGS | 0.0717 | 2.7e-93 | 1.5e-24 | 425 | 1379 | 233 | 1.38e+1 |
| ☐ | 85-GGCGGATCAK | STREME-85 | GGCGGATCAK | 0.0427 | 1.3e-120 | 3.3e-24 | 273 | 1041 | 180 | 1.11e+1 |
| ☐ | 111-GTGGATCACY | STREME-111 | GTGGATCACY | 0.0479 | 2.8e-110 | 2.7e-21 | 291 | 1119 | 209 | 9.92e+0 |
| ☐ | 23-AGGYGGATCA | STREME-23 | AGGYGGATCA | 0.0450 | 7.4e-125 | 9.3e-21 | 285 | 1022 | 188 | 9.94e+0 |
| ☐ | 54-CAGTGGCTCA | STREME-54 | CAGTGGCTCA | 0.0492 | 3.9e-75 | 1.7e-19 | 355 | 1490 | 460 | 5.09e+0 |
| ☐ | 73-GARCCYRGGARGTGGAGG | STREME-73 | GARCCYRGGARGTGGAGG | 0.0178 | 3.0e-103 | 2.2e-18 | 491 | 1751 | 404 | 1.28e+1 |
| ☐ | 7-AGCGAKHCTCC | STREME-7 | AGCGAKHCTCC | 0.0444 | 1.6e-63 | 8.2e-18 | 596 | 2088 | 656 | 1.02e+1 |
| ☐ | 141-AACACAGTGAAA | STREME-141 | AACACAGTGAAA | 0.0165 | 6.6e-71 | 3.0e-17 | 241 | 849 | 324 | 1.14e+1 |
| ☐ | 70-CAGRGCYCTG | STREME-70 | CAGRGCYCTG | 0.0309 | 8.8e-35 | 3.2e-17 | 373 | 2349 | 1818 | 3.62e+0 |
| ☐ | 20-CCCWGSCWGGG | STREME-20 | CCCWGSCWGGG | 0.0280 | 4.3e-37 | 3.8e-17 | 584 | 2865 | 1296 | 7.73e+0 |
| ☐ | 74-TGAGCCACCA | STREME-74 | TGAGCCACCA | 0.0677 | 5.3e-79 | 4.7e-17 | 353 | 1007 | 205 | 7.65e+0 |
| ☐ | 21-CAGCCCCWSC | STREME-21 | CAGCCCCWSC | 0.0370 | 7.7e-31 | 1.6e-15 | 405 | 2417 | 1540 | 7.05e+0 |
| ☐ | 132-AGGCAGAAGAA | STREME-132 | AGGCAGAAGAA | 0.0617 | 4.4e-67 | 2.0e-15 | 440 | 1123 | 282 | 1.12e+1 |
| ☐ | 58-AGWGGGGGGC | STREME-58 | AGWGGGGGGC | 0.0350 | 3.5e-25 | 2.5e-15 | 385 | 2357 | 1856 | 4.17e+0 |
| ☐ | 14-CAGGAGCWGGKKS | STREME-14 | CAGGAGCWGGKKS | 0.0325 | 3.8e-49 | 4.0e-15 | 444 | 2666 | 1633 | 7.55e+0 |
| ☐ | 18-CCAGGCCCHN | STREME-18 | CCAGGCCCHN | 0.0367 | 4.4e-44 | 8.1e-14 | 335 | 2000 | 1085 | 7.58e+0 |
| ☐ | 129-ACTGCAAGCTCCAC | STREME-129 | ACTGCAAGCTCCAC | 0.0136 | 2.4e-59 | 1.3e-13 | 497 | 1683 | 640 | 1.11e+0 |
| ☐ | 15-CTGGGWGSYS | STREME-15 | CTGGGWGSYS | 0.0309 | 1.1e-33 | 2.7e-12 | 355 | 2183 | 1401 | 5.95e+0 |
| ☐ | 165-CTGGTCTCAAA | STREME-165 | CTGGTCTCAAA | 0.0274 | 1.0e-66 | 3.7e-12 | 226 | 849 | 206 | 7.70e+0 |
| ☐ | 86-AGTGGCTCAT | STREME-86 | AGTGGCTCAT | 0.0357 | 7.4e-33 | 1.0e-11 | 549 | 1540 | 704 | 1.87e+0 |
| ☐ | 106-MAGAGCGAGA | STREME-106 | MAGAGCGAGA | 0.0284 | 6.3e-31 | 2.0e-11 | 587 | 1998 | 988 | 7.44e+0 |
| ☐ | 3-GCACTCCAGCCTGGGYGAC | STREME-3 | GCACTCCAGCCTGGGYGAC | 0.0282 | 7.2e-77 | 2.3e-9 | 584 | 2074 | 492 | 1.19e+1 |
| ☐ | 6-GCAGTGAGCCGAG | STREME-6 | GCAGTGAGCCGAG | 0.0113 | 1.2e-77 | 9.8e-9 | 530 | 1802 | 439 | 1.29e+1 |
| ☐ | 69-GGCTGGAGTG | STREME-69 | GGCTGGAGTG | 0.0226 | 9.5e-63 | 2.3e-8 | 579 | 2058 | 611 | 1.16e+1 |
| ☐ | 51-RCCATTCTCC | STREME-51 | RCCATTCTCC | 0.0468 | 8.9e-68 | 4.8e-8 | 449 | 993 | 226 | 1.04e+1 |
| ☐ | 65-AGAGCGAGAC | STREME-65 | AGAGCGAGAC | 0.0269 | 3.5e-38 | 4.9e-8 | 615 | 2430 | 1257 | 8.45e+0 |
| ☐ | 177-AAAAAATTTAAAAATT | STREME-177 | AAAAAATTTAAAAATT | 0.0294 | 5.2e-22 | 8.1e-8 | 311 | 996 | 661 | 4.76e+0 |
| ☐ | 147-AGCTGTGATGGC | STREME-147 | AGCTGTGATGGC | 0.0316 | 2.3e-27 | 2.5e-7 | 497 | 1313 | 551 | 1.65e+0 |
| ☐ | 10-AGTGGCGYGAT | STREME-10 | AGTGGCGYGAT | 0.0129 | 1.2e-53 | 1.8e-6 | 550 | 1538 | 432 | 1.06e+1 |
| ☐ | 63-CCHCACACMC | STREME-63 | CCHCACACMC | 0.0322 | 1.3e-7 | 2.8e-6 | 553 | 2966 | 2261 | 5.32e+0 |
| ☐ | 100-GTCTCAAAAAAA | STREME-100 | GTCTCAAAAAAA | 0.0288 | 2.0e-28 | 5.3e-6 | 673 | 1607 | 566 | 9.52e+0 |
| ☐ | 82-SCTGSSCAGS | STREME-82 | SCTGSSCAGS | 0.0310 | 4.2e-23 | 5.8e-6 | 385 | 2266 | 1637 | 5.70e+0 |
| ☐ | 110-AGGAGAATCA | STREME-110 | AGGAGAATCA | 0.0439 | 1.4e-20 | 3.1e-5 | 445 | 794 | 317 | 1.31e+1 |
| ☐ | 142-AGAGATCCTCC | STREME-142 | AGAGATCCTCC | 0.0308 | 1.5e-18 | 2.0e-4 | 460 | 2221 | 1673 | 2.95e+0 |
| ☐ | 68-AGAGTGAGACTCT | STREME-68 | AGAGTGAGACTCT | 0.0352 | 7.2e-30 | 3.1e-4 | 624 | 1262 | 421 | 7.00e+0 |
| ☐ | 13-GTCTCAAAAAAAAAAAAA | STREME-13 | GTCTCAAAAAAAAAAAAA | 0.0278 | 4.4e-28 | 3.6e-4 | 706 | 1874 | 690 | 1.33e+1 |
| ☐ | 154-AAAAAAAAAAAATT | STREME-154 | AAAAAAAAAAAATT | 0.0217 | 6.5e-12 | 8.4e-4 | 655 | 1860 | 1130 | 1.40e+1 |
| ☐ | 124-AGAGCAAAACTC | STREME-124 | AGAGCAAAACTC | 0.0325 | 1.2e-16 | 3.6e-3 | 609 | 1259 | 574 | 5.12e-1 |
| ☐ | 115-AGCAATCCACC | STREME-115 | AGCAATCCACC | 0.0410 | 1.7e-22 | 4.7e-3 | 294 | 846 | 429 | 8.13e-1 |
| ☐ | 19-GGGAGGAGRS | STREME-19 | GGGAGGAGRS | 0.0320 | 4.7e-8 | 2.0e-2 | 441 | 2479 | 1993 | 5.71e+0 |
| ☐ | 77-AGGGCCAGAG | STREME-77 | AGGGCCAGAG | 0.0261 | 2.9e-6 | 3.9e-2 | 455 | 2556 | 2232 | 3.56e+0 |
| ☐ | 87-CCCCTCCTGA | STREME-87 | CCCCTCCTGA | 0.0286 | 1.3e-20 | 1.0e-1 | 313 | 1937 | 1568 | 5.21e+0 |
| ☐ | 38-CATCACAATTAAAAGAACTA | STREME-38 | CATCACAATTAAAAGAACTA | 0.0290 | 8.8e-11 | 1.5e-1 | 323 | 995 | 746 | 3.45e-1 |
| ☐ | 22-AAAAAAAAAAAAAA | STREME-22 | AAAAAAAAAAAAAA | 0.0215 | 1.1e-10 | 6.0e-1 | 657 | 1694 | 858 | 6.37e+0 |
| ☐ | 105-GCTTGAAGGCA | STREME-105 | GCTTGAAGGCA | 0.0190 | 2.1e-7 | 1.6 | 512 | 2631 | 2226 | 4.62e+0 |
| ☐ | 135-CCCCTCCWWG | STREME-135 | CCCCTCCWWG | 0.0283 | 2.1e-11 | 8.8 | 423 | 2141 | 1456 | 7.41e+0 |
| ☐ | 24-CYCAGCTCTGC | STREME-24 | CYCAGCTCTGC | 0.0227 | 2.3e-2 | 18.3 | 516 | 2758 | 2266 | 5.02e+0 |
| ☐ | 144-AGTGGCACTAT | STREME-144 | AGTGGCACTAT | 0.0145 | 4.0e-7 | 22.3 | 560 | 1186 | 670 | 8.93e+0 |

##### Matching sequences (out of 5000) ?

**Union: 2076 sequences (42%).**
**Intersection: 872 sequences (17%).**

```
chr14:35496174-35497174
chr14:92531436-92532436
chr14:74629549-74630549
chr12:20899604-20900604
chr14:38562934-38563934
chr13:114108614-114109614
chr10:43603062-43604062
chr14:63807562-63808562
chr10:72594704-72595704
chr10:103413284-103414284
```

##### Filter & Sort

**Filters** ?
- ☐ Top 10
- ☐ Database is streme
- ☐ ID matches .*
- ☐ Alt ID matches .*
- ☐ *E*-value ≤ 1
- ☐ Fisher *E*-value≤ 1
- ☐ Region Width ≤ 200
- ☐ Negative set *E*-value ≥ 1

**Sort** ?
Motifs: Fisher E-value

Update

##### Columns to display

- ☐ Show Database
- ☑ Show ID
- ☑ Show Alt ID
- ☑ Show Consensus
- ☑ Show Concentration
- ☑ Show *E*-value
- ☐ Show *p*-value
- ☑ Show Region Width
- ☑ Show Region Matches
- ☐ Show Sequence Matches
- ☐ Show Negative *p*-value
- ☑ Show Negative Region Matches
- ☐ Show Negative Sequence Matches
- ☐ Show Max Probability
- ☐ Show Max Probability Location
- ☐ Show Multiple Tests
- ☑ Show Score Threshold

| ☒ | ID | Alt ID | Consensus | Concentration | E-value | Fisher E-value | Region Width | Region Matches | Negative Region Matches | Score Threshold |
|---|---|---|---|---|---|---|---|---|---|---|
| ☐ | 148-TACACCTCCCA | STREME-148 | TACACCTCCCA | 0.0153 | 3.0e-3 | 34.0 | 472 | 966 | 564 | 6.36e+0 |
| ☐ | 78-AGGGGGACAG | STREME-78 | AGGGGGACAG | 0.0298 | 8.6e-8 | 70.8 | 597 | 1661 | 766 | 1.04e+1 |
| ☐ | 26-AATCCTGAGTTCTAGTTTGAT | STREME-26 | AATCCTGAGTTCTAGTTTGAT | 0.0678 | 4.2e-17 | 92.2 | 342 | 57 | 3 | 1.18e+1 |
| ☐ | 99-TCAGCACCACATYRCACT | STREME-99 | TCAGCACCACATYRCACT | 0.0156 | 5.0e-5 | 141.5 | 573 | 62 | 22 | 1.02e+1 |
| ☐ | 29-AGATTCTGGTATGTTGTGTCTTTG | STREME-29 | AGATTCTGGTATGTTGTGTCTTTG | 0.0833 | 4.6e-10 | 160.0 | 307 | 35 | 0 | 4.00e+1 |
| ☐ | 34-AATCAACAGAATATACATTYTTY | STREME-34 | AATCAACAGAATATACATTYTTY | 0.0000 | 1.1e-3 | 161.8 | 614 | 56 | 3 | 2.45e+1 |
| ☐ | 37-ATTTGCTGAGGAGWGCTTTACTTCCAA | STREME-37 | ATTTGCTGAGGAGWGCTTTACTTCCAA | 0.0536 | 1.4e-9 | 178.9 | 484 | 55 | 4 | 3.46e+1 |
| ☐ | 40-GTTTGTTATAATTTCTGTTCTTTTA | STREME-40 | GTTTGTTATAATTTCTGTTCTTTTA | 0.0175 | 1.9e-10 | 178.9 | 430 | 55 | 5 | 1.44e+1 |
| ☐ | 149-AGCTATCCTCC | STREME-149 | AGCTATCCTCC | 0.0807 | 7.8e-10 | 179.0 | 40 | 66 | 8 | 1.03e+1 |
| ☐ | 127-AAAGAAAAGAAAAGAAA | STREME-127 | AAAGAAAAGAAAAGAAA | 0.0152 | 2.1e-5 | 179.0 | 708 | 1361 | 744 | 1.04e+1 |
| ☐ | 88-CCTCCHTCCY | STREME-88 | CCTCCHTCCY | 0.0454 | 4.0e-3 | 179.0 | 31 | 107 | 23 | 1.19e+1 |
| ☐ | 28-CAGTTTCCATGTAGCTGAGCGGTTT | STREME-28 | CAGTTTCCATGTAGCTGAGCGGTTT | 0.1356 | 1.6e-18 | 179.0 | 248 | 53 | 3 | 1.56e+1 |
| ☐ | 39-AATGACTACTGGGTACATAA | STREME-39 | AATGACTACTGGGTACATAA | 0.0385 | 1.3e-15 | 179.0 | 205 | 44 | 1 | 2.61e+1 |
| ☐ | 27-AAGGCAGAAATAAAGATGTTCTTT | STREME-27 | AAGGCAGAAATAAAGATGTTCTTT | 0.0667 | 9.0e-10 | 179.0 | 195 | 35 | 1 | 3.85e+1 |
| ☐ | 98-CACAGCACATGTTTCAGA | STREME-98 | CACAGCACATGTTTCAGA | 0.1500 | 4.1e-9 | 179.0 | 85 | 17 | 0 | 2.41e+1 |
| ☐ | 126-TCACAGATCAACAGGATCCC | STREME-126 | TCACAGATCAACAGGATCCC | 0.1364 | 3.2e-6 | 179.0 | 161 | 19 | 0 | 1.04e+1 |
| ☐ | 48-GAAACCAAYGAGAA | STREME-48 | GAAACCAAYGAGAA | 0.0816 | 9.7e-6 | 179.0 | 269 | 38 | 1 | 1.87e+1 |
| ☐ | 31-ATGTGGTCAATTTTGGAATA | STREME-31 | ATGTGGTCAATTTTGGAATA | 0.0000 | 1.5e-5 | 179.0 | 535 | 59 | 6 | 1.40e+1 |
| ☐ | 113-TTAAATGGATTAA | STREME-113 | TTAAATGGATTAA | 0.2000 | 2.6e-4 | 179.0 | 156 | 14 | 0 | 2.35e+1 |
| ☐ | 59-CTGGGACACATTYAAA | STREME-59 | CTGGGACACATTYAAA | 0.0750 | 4.3e-4 | 179.0 | 347 | 35 | 2 | 2.38e+1 |
| ☐ | 119-ATCTGTTTAACAAAGC | STREME-119 | ATCTGTTTAACAAAGC | 0.0000 | 7.6e-4 | 179.0 | 199 | 16 | 0 | 2.47e+1 |
| ☐ | 94-CATTTTGTTCTGTACTAAGAAGAA | STREME-94 | CATTTTGTTCTGTACTAAGAAGAA | 0.0000 | 8.6e-4 | 179.0 | 241 | 20 | 0 | 2.82e+1 |
| ☐ | 33-AGCACTAAATGCCCACAAGAGAAAGCGG | STREME-33 | AGCACTAAATGCCCACAAGAGAAAGCGG | 0.0000 | 6.8e-3 | 179.0 | 469 | 46 | 3 | 3.70e+1 |
| ☐ | 97-GTTTGTGTCCCTGGGTACTTGAGA | STREME-97 | GTTTGTGTCCCTGGGTACTTGAGA | 0.0000 | 7.8e-3 | 179.0 | 327 | 19 | 0 | 1.57e+1 |
| ☐ | 95-GCATGCTCGTTAAG | STREME-95 | GCATGCTCGTTAAG | 0.0556 | 1.8e-2 | 179.0 | 251 | 17 | 0 | 2.05e+1 |
| ☐ | 156-ACTGCACCACT | STREME-156 | ACTGCACCACT | 0.0203 | 6.7e-2 | 179.0 | 610 | 948 | 587 | 9.21e-1 |
| ☐ | 167-AAACAGTGTAAACCGGCAGTGT | STREME-167 | AAACAGTGTAAACCGGCAGTGT | 0.3636 | 2.2e-1 | 179.0 | 71 | 8 | 0 | 3.90e+0 |
| ☐ | 160-AAGGTCAAGTTGTCTGGAC | STREME-160 | AAGGTCAAGTTGTCTGGAC | 0.0201 | 5.2e-1 | 179.0 | 366 | 452 | 326 | 1.97e+0 |
| ☐ | 125-AGCAATCTGCC | STREME-125 | AGCAATCTGCC | 0.0335 | 6.3e-1 | 179.0 | 238 | 86 | 38 | 9.00e+0 |
| ☐ | 137-TGTCTGTGTAGAAAGAA | STREME-137 | TGTCTGTGTAGAAAGAA | 0.0000 | 1.5 | 179.0 | 286 | 18 | 1 | 1.43e+1 |
| ☐ | 36-CTGTAGATGTCTATTA | STREME-36 | CTGTAGATGTCTATTA | 0.0135 | 1.6 | 179.0 | 681 | 70 | 31 | 8.60e+0 |
| ☐ | 103-GTGGGGAAAAGATTGAG | STREME-103 | GTGGGGAAAAGATTGAG | 0.0000 | 1.6 | 179.0 | 380 | 22 | 1 | 1.72e+1 |
| ☐ | 45-TAGGGTGTCAATTTTRGATCTTT | STREME-45 | TAGGGTGTCAATTTTRGATCTTT | 0.0222 | 3.3 | 179.0 | 522 | 40 | 1 | 3.47e+1 |

#### Input Files

##### Alphabet

**Background source:** the file 'pos.fa.bg'

**Background source:** the file 'pos.fa.bg'

| Name | Bg. | | | | Bg. | Name |
|---|---|---|---|---|---|---|
| Adenine | 0.2784 | A | ~ | T | 0.2784 | Thymine |
| Cytosine | 0.2216 | C | ~ | G | 0.2216 | Guanine |


| Name | Bg. | | | | Bg. | Name |
|---|---|---|---|---|---|---|
| Adenine | 0.2784 | A | ~ | T | 0.2784 | Thymine |
| Cytosine | 0.2216 | C | ~ | G | 0.2216 | Guanine |


##### Sequences

| Database | Source | Sequence Count |
|---|---|---|
| pos | pos.fa | 5000 |

##### Negative sequences

| Database | Source | Sequence Count |
|---|---|---|
| neg | neg.fa | 5000 |

##### Motifs

| Database | Source | Motif Count |
|---|---|---|
| streme | streme.txt | 179 |

##### Other Settings

| | |
|---|---|
| **Objective Function** | central region enrichment (CE) |
| **Convert Motifs to Different Alphabet?** | No |
| **Motif Pseudo-Counts** | 0.1 |
| **Required sequence length** | None |
| **Site Scoring Method** | log-odds scores |
| **Score Threshold** | 0 (bits) |
| **Optimize Score Threshold?** | Yes |
| **Minimum Region Size** | 0 |
| **Maximum Region Size** | 0 |
| **Strand Handling** | scan both strands if alphabet is complementable |
| **Plotting of Matches on Negative Strand** | same as for positive strand |
| **Sequence IDs Included in Output?** | Yes |

**CentriMo version**
5.5.7 (Release date: Wed Jun 19 13:59:04 2024 -0700)


**Reference**
Timothy L. Bailey and Philip Machanick, "Inferring direct DNA binding from ChIP-seq", *Nucleic Acids Research*, **40**:e128, 2012. [Full Text]


**Reference**
Timothy L. Bailey and Philip Machanick, "Inferring direct DNA binding from ChIP-seq", *Nucleic Acids Research*, **40**:e128, 2012. [Full Text]


**Command line summary**

```
centrimo --oc . --verbosity 1 --optimize-score --score 0.0 --ethresh 10.0 --bfile pos.fa.bg --neg neg.fa pos.fa streme.txt
```

### SpaMo
Spaced Motif Analysis Tool

For further information on how to interpret these results please access https://meme-suite.org/meme/doc/spamo-output-format.html.
To get a copy of the MEME software please access https://meme-suite.org.

If you use SpaMo in your research, please cite the following paper:
Tom Whitington, Martin C. Frith, James Johnson and Timothy L. Bailey "Inferring transcription factor complexes from ChIP-seq data", *Nucleic Acids Res.* **39**(15):e98, 2011. [full text] If you use SpaMo in your research, please cite the following paper:
Tom Whitington, Martin C. Frith, James Johnson and Timothy L. Bailey "Inferring transcription factor complexes from ChIP-seq data", *Nucleic Acids Res.* **39**(15):e98, 2011. [full text]

Primary Motifs | Sequence Database | Secondary Motif Databases | Spacing Analysis | Inputs and Settings | Program information | Results in TSV Format ? | Contributing Sequence IDs [Download Plain] [Download BED] ?

#### Primary Motifs

Next Top

| Database ? | Name ? | Preview ? | Significant Secondaries ? | List ? |
|---|---|---|---|---|
| top_motifs.txttop_motifs.txt | 1-GGGAGGCYGAGGCR (STREME-1)1-GGGAGGCYGAGGCR (STREME-1) | 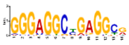 | 2020 | 16-GGCAGATCACC (STREME-16), 7-AGCGAKHCTCC (STREME-7), 2-GCTGGGAYTACAGGCG (STREME-2), 4-CTCCTGACCTCR (STREME-4), 11-CCACCVCVCC (STREME-11), 8-CVAGACCAKCCY (STREME-8), 12-CCRTGTTGGCC (STREME-12), 5-CCCCGTCTCTAC (STREME-5), 15-CTGGGWGSYS (STREME-15), 3-GCACTCCAGCCTGGGYGAC (STREME-3), 6- |

##### Alphabet

**Background source:** built from the (primary) sequences

**Background source:** built from the (primary) sequences

| Name ? | Bg. ? | | | | Bg. ? | Name ? |
|---|---|---|---|---|---|---|
| Adenine | 0.278416 | A | ~ | T | 0.278416 | Thymine |
| Cytosine | 0.221584 | C | ~ | G | 0.221584 | Guanine |

| Name ? | Bg. ? | | | | Bg. ? | Name ? |
|---|---|---|---|---|---|---|
| Adenine | 0.278416 | A | ~ | T | 0.278416 | Thymine |
| Cytosine | 0.221584 | C | ~ | G | 0.221584 | Guanine |

#### Sequence Database

Next Previous Top

| Name ? | Last Modified ? | Contained ? | Too Short ? | Too Masked ? | No Primary ? | Too Similar ? | Used ? |
|---|---|---|---|---|---|---|---|
| pospos | Wed Aug 28 03:13:25 2024Wed Aug 28 03:13:25 2024 | 50005000 | 00 | 00 | 16271627 | 19131913 | 14601460 |

#### Secondary Motif Databases

Next Previous Top

| Name ? | Last Modified ? | Number of Motifs ? | Motifs Significant ? | Motifs Redundant ? |
|---|---|---|---|---|
| secondariessecondaries | Wed Aug 28 03:13:25 2024Wed Aug 28 03:13:25 2024 | 2020 | 1717 | 33 |

#### Settings

Next Previous Top

**Match Score Threshold** ? 7 (bits)
**Margin size** ? 150
Show Advanced Settings

#### Spacing Analysis for 1-GGGAGGCYGAGGCR (STREME-1)

Next Previous Top

**Secondary Motif:** 16-GGCAGATCACC (STREME-16) ? **Cluster:** 16-GGCAGATCACC (STREME-16) ? **E-value:** 0.00e+0 ? **Best Gap:** 0 ? **Best Orientation:** downstream / same strand ?

**Primary Motif Logo** ?

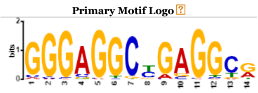

**Secondary Motif Logo** ?

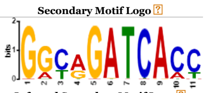

**Inferred Secondary Motif Logo** ?

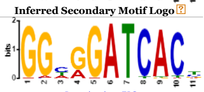

Download as EPS

**Alignment Logo** ?

Up+
Up-
Dn+
Dn-

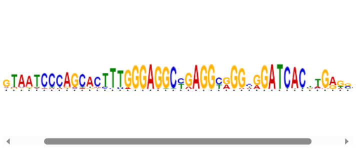

Download as EPS

**Spacings** ?

| Gap | Orientation | *p*-value |
|---|---|---|
| 0 | dn+ | 0.00e+0 |
| 0 | up-/dn+ | 1.92e-300 |
| 0 | dn+/dn- | 1.92e-300 |
| 0 | all | 2.67e-255 |
| 110 | up+ | 1.33e-6 |

Highlight: Selected All

**Overview Graph** ?

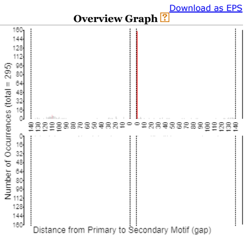


Download as EPS

**Selected Orientation Graph** ?

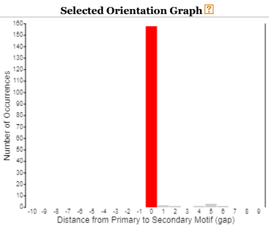


Download as EPS

**Contributing Sequence IDs (158)** ?

```
chr10:100001286-100002286
chr10:100814020-100815020
chr10:101439966-101440966
chr10:1014910-1015910
chr10:101546014-101547014
chr10:101857798-101858798
chr10:102844308-102845308
chr10:103152076-103153076
chr10:103165452-103166452
chr10:103564080-103565080
chr10:103596464-103597464
chr10:103856146-103857146
chr10:105056176-105057176
chr10:105134108-105135108
chr10:106047322-106048322
chr10:108425820-108426820
chr10:108577772-108578772
chr10:11003456-11004456
chr10:111070024-111071024
chr10:111514494-111515494
```

Plain Format Download

##### Secondaries ?

| Lock ? | ID ? | Name ? | Cluster ? | E-value ? | Best Gap ? | Best Orientation ? | Spacings ? |
|---|---|---|---|---|---|---|---|
| ☐ | 16-GGCAGATCACC | STREME-16 | 16-GGCAGATCACC | 0.00e+0 | 0 | downstream / same strand | |
| ☐ | 7-AGCGAKHCTCC | STREME-7 | 7-AGCGAKHCTCC | 0.00e+0 | 0 | downstream / opposite strand | |
| ☐ | 2-GCTGGGAYTACAGGCG | STREME-2 | 2-GCTGGGAYTACAGGCG | 0.00e+0 | 5 | upstream / opposite strand | |
| ☐ | 4-CTCCTGACCTCR | STREME-4 | 4-CTCCTGACCTCR | 0.00e+0 | 11 | downstream / opposite strand | |
| ☐ | 11-CCACCVCVCC | STREME-11 | 11-CCACCVCVCC | 7.78e-303 | 25 | upstream / opposite strand | |
| ☐ | 8-CVAGACCAKCCY | STREME-8 | 8-CVAGACCAKCCY | 9.06e-184 | 25 | downstream / same strand | |
| ☐ | 12-CCRTGTTGGCC | STREME-12 | 12-CCRTGTTGGCC | 1.99e-134 | 37 | downstream / opposite strand | |
| ☐ | 5-CCCCGTCTCTAC | STREME-5 | 5-CCCCGTCTCTAC | 8.65e-122 | 53 | downstream / same strand | |
| ☐ | 15-CTGGGWGSYS | STREME-15 | 15-CTGGGWGSYS | 1.24e-110 | 28 | upstream / primary palindromic | |
| ☐ | 3-GCACTCCAGCCTGGGYGAC | STREME-3 | 3-GCACTCCAGCCTGGGYGAC | 5.02e-108 | 25 | downstream / same strand | |
| ☐ | 6-GCAGTGAGCCGAG | STREME-6 | 6-GCAGTGAGCCGAG | 7.88e-102 | 32 | downstream / same strand | |
| ☐ | 10-AGTGGCGYGAT | STREME-10 | 10-AGTGGCGYGAT | 2.59e-101 | 45 | downstream / opposite strand | |
| ☐ | 20-CCCWGSCWGGG | STREME-20 | 3-GCACTCCAGCCTGGGYGAC | 3.82e-101 | 29 | downstream / secondary palindromic | |
| ☐ | 18-CCAGGCCCHN | STREME-18 | 18-CCAGGCCCHN | 2.16e-78 | 31 | downstream / opposite strand | |
| ☐ | 9-AAAAATACAAAAAWT | STREME-9 | 9-AAAAATACAAAAAWT | 8.48e-66 | 66 | downstream / same strand | |
| ☐ | 14-CAGGAGCWGGKKS | STREME-14 | 2-GCTGGGAYTACAGGCG | 2.34e-58 | 0 | upstream / opposite strand | |
| ☐ | 21-CAGCCCCWSC | STREME-21 | 21-CAGCCCCWSC | 6.75e-47 | 31 | downstream / same strand | |
| ☐ | 13-GTCTCAAAAAAAAAAAAAA | STREME-13 | 13-GTCTCAAAAAAAAAAAAAA | 4.41e-43 | 88 | downstream / same strand | |
| ☐ | 17-RGGCWGAGGS | STREME-17 | 2-GCTGGGAYTACAGGCG | 4.37e-42 | 17 | upstream / opposite strand | |
| ☐ | 19-GGGAGGAGRS | STREME-19 | 19-GGGAGGAGRS | 4.41e-14 | 57 | downstream / opposite strand | |

**Filter** ?
☐ Top 10
☐ ID matches .*
☐ Name matches .*
☐ Cluster matches .*
☐ *E*-value ≤ 1
☐ Gaps (ranges allowed) 0-10

**Sort** ?
Sort by E-value
Update

Previous Top

**SpaMo version**
5.5.75.5.7 (Release date: Wed Jun 19 13:59:04 2024 -0700Wed Jun 19 13:59:04 2024 -0700)

**Reference**
Tom Whitington, Martin C. Frith, James Johnson and Timothy L. Bailey "Inferring transcription factor complexes from ChIP-seq data", *Nucleic Acids Res.* **39**(15):e98, 2011. [full text]
**Reference**
Tom Whitington, Martin C. Frith, James Johnson and Timothy L. Bailey "Inferring transcription factor complexes from ChIP-seq data", *Nucleic Acids Res.* **39**(15):e98, 2011. [full text]

**Command line**

```
spamo -verbosity 1 -oc . -numgen 1 pos.fa top_motifs.txt secondaries.meme
```

Result calculation took 1010 seconds
